# MAXWELL: Calibrating the probabilistic outputs of protein language models to the mutation-induced stability change landscape

**DOI:** 10.64898/2026.07.29.741631

**Authors:** Mingchen Li, Xiaoran Cheng, Meiyao Hong, Fan Jiang, Liang Hong, Yuanxi Yu

**Author notes:** These authors contributed equally to this work (Mingchen Li, Xiaoran Cheng).

## Abstract

Designing mutations that enhance protein stability is a central goal in protein engineering. However, experimentally screening large numbers of candidate mutations is costly and time-consuming, creating a strong need for computational methods that can identify potentially stabilizing mutations. Among these approaches, protein language models are particularly promising because they learn context-dependent amino acid preferences from large-scale sequence and structure datasets. Nevertheless, most existing stability prediction methods use these models primarily as feature extractors and do not fully exploit the amino acid probability distributions they encode. Here, we introduce MAXWELL (Matrix-wise Landscape Learning), a novel post-training method that calibrates the probabilistic outputs learned by protein language models during pretraining to generate mutational landscapes that quantify the effects of individual amino acid substitutions on protein stability. When applied to ProteinMPNN, MAXWELL yields a state-of-the-art predictor of the effects of protein mutations on stability, outperforming ThermoMPNN and other representative methods on a curated benchmark of experimentally measured stability changes. We next applied MAXWELL to the design of ten single-point mutations in the DhaA dehalogenase, seven of which (70%) increased thermal stability. Among them, G171W showed the largest improvement, with a measured **Δ*T***_***m***_ of 4.91 °C. These experimental results establish MAXWELL as a novel post-training strategy for protein language models and a practical framework for designing stabilizing mutations.

## 1 Introduction

Improving protein stability is a key objective in protein engineering because stability determines whether designed proteins remain folded, soluble and active under operational conditions [1–3]. For enzymes, stabilizing mutations can expand usable temperature ranges and support further functional optimization, whereas for biologics they affect expression, storage, manufacturability and developability [4–6]. Because sequence space is large and stability assays are costly, time consuming and hard to scale, computational filters are needed before wet-lab testing [7, 8].

Computational stability prediction has progressed from energy functions and empirical statistical models to deep learning and pretrained protein language models. Energy and statistical methods provide interpretable estimates, but they can depend on structure quality, hand-designed descriptors and empirical potentials [9– 14]. Deep learning methods replaced many such descriptors with learned sequence or structural representations [15–20]. More recently, pretrained protein models have enabled zero-shot mutation-effect prediction [21, 22]. Representative approaches include structure-conditioned inverse folding with ESM-IF1 [23], retrieval-enhanced scoring with VenusREM [24], conservation and solvent-accessibility scoring with RSALOR [25], joint sequence–structure encoding with ProtSSN [26], and expert-guided prediction with VespaG [27]. Large-scale stability datasets have also supported supervised predictors such as ThermoMPNN [28]. Whereas zero-shot methods use pre-trained or evolutionary signals without fitting to the stability labels evaluated here, MAXWELL directly calibrates native amino-acid probabilities against experimental stability measurements.

To address this limitation, we introduce MAXWELL, a general post-training frame-work that directly harnesses the probabilistic outputs of protein language models for predicting mutation effects on protein stability. Rather than using a pretrained model solely as a feature extractor, MAXWELL fine-tunes its position-resolved amino-acid scores against experimental stability measurements and converts them into a complete single-mutation landscape in one forward pass. Each landscape entry provides a stability-oriented score for a specific amino-acid substitution, enabling the systematic ranking of all single-site variants in a protein. By retaining the native sequence-to-landscape output interface of the underlying model, MAXWELL can be applied to proteins that are not represented in the training set.

We evaluated MAXWELL across multiple protein language models and found that ProteinMPNN [29] provided the strongest backbone. We therefore post-trained ProteinMPNN on cDNA-display stability data [30] and evaluated the resulting model on a curated benchmark compiled from public protein-stability and mutation-effect resources [20, 31–35]. Across this benchmark, MAXWELL outperformed ThermoMPNN [28] and other representative baselines. To assess its practical utility for protein engineering, we applied MAXWELL to the haloalkane dehalogenase DhaA [36]. Of ten single-point variants selected from the predicted landscape, seven showed increased melting temperatures, with activity measurements identifying variants that retained substantial catalytic function. These findings support MAXWELL as a general post-training strategy for adapting diverse protein language models to mutational thermostability prediction and prioritizing candidates for experimental protein engineering.

## 2 Results

### 2.1 MAXWELL calibrates the probabilistic outputs of protein language models to the mutational stability change landscape

#### Concept of calibration

Deep learning-based stability predictors typically use protein language models only as feature extractors, regressing mutation effects on protein stability from their embeddings to guide variant selection. In contrast, MAXWELL directly transforms the output probability distributions of a protein language model into a mutation-stability landscape under supervision from experimentally measured stability effects, a process we term “calibration” (Fig. 1a). Before calibration, the protein language model outputs a position-by-amino-acid probability matrix that specifies, at each sequence position, the predicted probabilities of the 20 standard amino acids. After calibration, the matrix retains the same dimensions, but each entry represents the predicted effect of the corresponding single-point mutation on protein stability. This calibration procedure can be applied to any protein language model that provides position-aligned amino-acid probabilities. Among the models evaluated, ProteinMPNN [29] was selected as the backbone for the main MAXWELL implementation because it achieved the best predictive performance on the curated benchmark (Table 2).

**Fig. 1.**
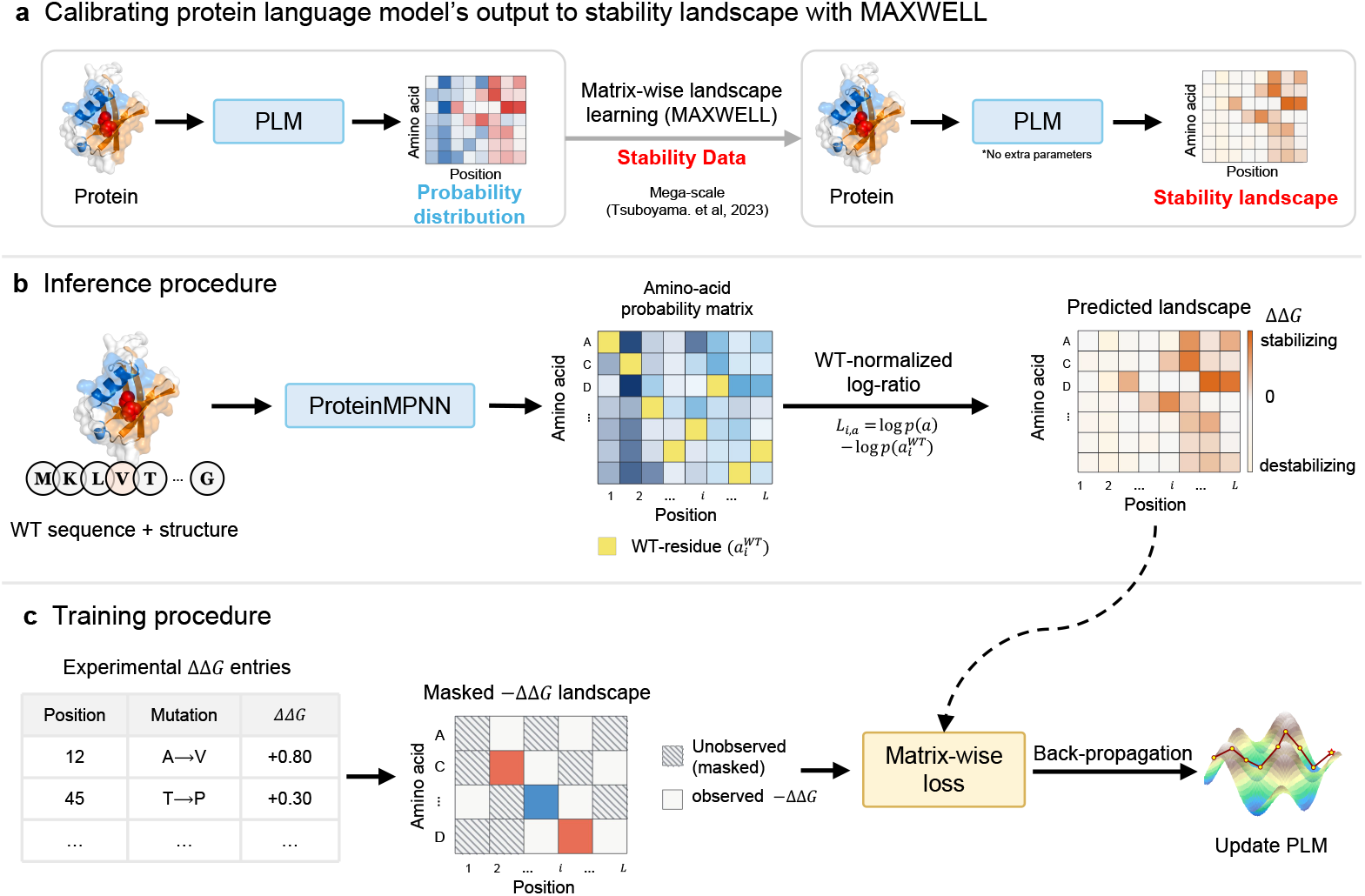
Calibration of protein language model outputs for predicting mutation effects on protein stability using MAXWELL. All matrix schematics in panels a–c are shown transposed relative to the mathematical convention in the text: rows represent amino-acid identities and columns represent sequence positions. **a**, Conceptual overview of MAXWELL. A protein language model produces an amino-acid probability matrix, with each entry representing the predicted probability of an amino acid at a given sequence position. MAXWELL calibrates these probabilistic outputs into a landscape of the same dimensions, where each entry predicts the effect of the corresponding single-point mutation on protein stability relative to the wild type. **b**, Forward computation and inference. The wild-type sequence and structural context are provided to the protein language model backbone, which outputs a position-by-amino-acid log-probability matrix. Each entry gives the log probability assigned to an amino acid at a given sequence position. This matrix is converted into a wild-type-normalized log-probability-ratio landscape according to Eq. 1. After calibration, each entry is a stability-oriented mutation score, with higher values indicating greater predicted stability. These scores are not estimates of absolute ΔΔ*G* in physical units. **c**, Training procedure. For each training protein, experimentally measured mutations and their associated ΔΔ*G* values are organized into a mutation table. The values are sign-reversed to construct a position-by-amino-acid target landscape of *−*ΔΔ*G*, so that larger target values indicate greater stabilization. Entries without measurements are masked and excluded from the loss. A masked Pearson-correlation loss between the predicted score landscape and the experimental target landscape is backpropagated to optimize the parameters of the protein language model.

#### Forward computation and inference process

Given a wild-type sequence and structural context, the protein language model (ProteinMPNN in this case) outputs a position-by-amino-acid log-probability matrix **Q**. Here, *Q*_*ij*_ is the log-probability the model assigns to amino acid *j* at position *i*. In the mathematical formulation, rows correspond to sequence positions and columns to amino-acid identities. For visualization, the matrix schematics in Fig. 1 are shown transposed, with positions on the horizontal axis and amino-acid identities on the vertical axis. MAXWELL converts **Q** into a mutation landscape **L** by subtracting, at each position *i*, the log-probability of the wild-type residue *a*_*i*_ from the log-probabilities of all amino acids:

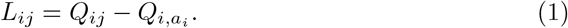

This subtraction is implemented as a single differentiable matrix operation for efficient computation and training (Methods, Eq. (7)). Each entry *L*_*ij*_ scores the substitution of amino acid *j* at position *i* relative to the wild-type residue. **L** is therefore a wild-type-normalized, log-probability-ratio mutation landscape over all positions and amino-acid identities (Fig. 1b).

#### Training and calibration process

We trained MAXWELL on a protein mutation stability dataset curated from cDNA display experiments [30], following the training process illustrated in Fig. 1c. For each protein in the dataset, we assembled a mutation table containing its measured ΔΔ*G* values. Following the conventional definition ΔΔ*G* = Δ*G*_mut_ −Δ*G*_WT_, negative ΔΔ*G* values indicate stabilizing mutations; therefore, for training and evaluation we used the sign-reversed target, −ΔΔ*G*, so that larger target values consistently correspond to higher mutant stability. We then converted this sign-reversed table into a stability landscape, masking out unmeasured entries so that they did not contribute to the loss.

Since both the masking operation and the landscape transformation in Eq. (1) are differentiable, MAXWELL’s parameters can be optimized end-to-end by gradient descent. We minimized a masked Pearson-correlation loss between the predicted land-scape and the observed entries of the experimental target landscape (see more details in Methods). After training, MAXWELL produces a complete stability-oriented land-scape that ranks all single amino-acid substitutions in a protein through one single forward pass.

Below, we show that MAXWELL achieves state-of-the-art thermostability prediction performance on a curated benchmark and enables real-world protein design for thermostability enhancement.

### 2.2 MAXWELL achieves state-of-the-art performance in predicting mutation effects on protein stability

#### Benchmark setup

To evaluate the performance of MAXWELL, we constructed a benchmark dataset from multiple sources of protein mutation stability data [20, 31–35], referred to as Test12K in the following text, containing 12,102 mutations and their measured ΔΔ*G* values across 308 proteins, with training proteins filtered below 30% sequence identity to any Test12K protein to avoid data leakage. We benchmarked MAXWELL against eleven baselines: ThermoMPNN[28], HERMES[37], Mutate Everything[38], FoldX[10], Rosetta[9], ProteinMPNN zero-shot scoring[29], ESMIF1[23], VenusREM[24], RSALOR[25], ProtSSN[26], and VespaG[27]. All correlation metrics were calculated against −ΔΔ*G*, so that higher experimental and predicted values both indicated stronger stabilization. Evaluation used average per-protein correlation metrics, computed by correlating predicted and measured stability scores within each protein before averaging across proteins so that proteins with many mutations do not dominate the score. Detailed dataset construction, filtering, leakage control and baseline inclusion procedures are described in Methods.

#### Benchmark results

Table 1 summarizes performance on the curated benchmark. MAXWELL achieved the highest mean performance across all five reported metrics. Per-protein paired comparisons showed higher Spearman correlations than all eleven baselines (two-sided paired Wilcoxon tests, all *P <* 0.001; Supplementary Table S7). With 1.66 million parameters, MAXWELL used approximately 62% fewer parameters than ThermoMPNN (4.34 million). By calibrating ProteinMPNN’s native amino-acid distributions, MAXWELL provides a compact approach to stability prediction without introducing a separate stability-regression network. Protein-length-stratified results are reported in Supplementary Table S8.

**Table 1.**
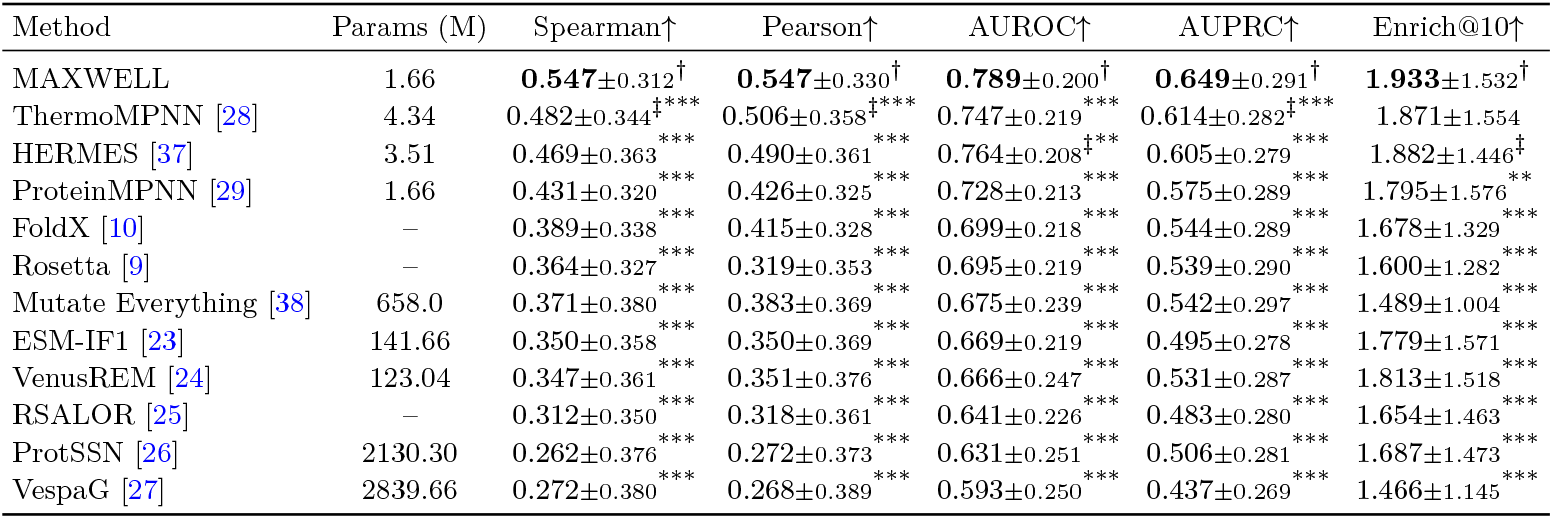
Performance evaluation on curated protein stability landscapes. We benchmarked MAXWELL and the baseline methods on Test12K, a curated dataset of experimentally measured effects of single-point mutations on protein stability. Enrich@10 is the fold-enrichment of true stabilizing mutations among each protein’s top-10 model-ranked candidates, relative to the background stabilizing-mutation rate. Bold values with *†* mark the best method in each column, and *‡* marks the second-best method. Asterisks mark paired Wilcoxon tests against MAXWELL, with ^*\**^*P <* 0.05, ^*\*\**^*P <* 0.01, ^*\*\*\**^*P <* 0.001.

| Method | Params (M) | Spearman↑ | Pearson↑ | AUROC↑ | AUPRC↑ | Enrich@10↑ |
|---|---|---|---|---|---|---|
| MAXWELL | 1.66 | **0.547**<sub>±0.312</sub><sup>†</sup> | **0.547**<sub>±0.330</sub><sup>†</sup> | **0.789**<sub>±0.200</sub><sup>†</sup> | **0.649**<sub>±0.291</sub><sup>†</sup> | **1.933**<sub>±1.532</sub><sup>†</sup> |
| ThermoMPNN [28] | 4.34 | 0.482<sub>±0.344</sub><sup>‡***</sup> | 0.506<sub>±0.358</sub><sup>‡***</sup> | 0.747<sub>±0.219</sub><sup>***</sup> | 0.614<sub>±0.282</sub><sup>‡***</sup> | 1.871<sub>±1.554</sub> |
| HERMES [37] | 3.51 | 0.469<sub>±0.363</sub><sup>***</sup> | 0.490<sub>±0.361</sub><sup>***</sup> | 0.764<sub>±0.208</sub><sup>‡**</sup> | 0.605<sub>±0.279</sub><sup>***</sup> | 1.882<sub>±1.446</sub><sup>‡</sup> |
| ProteinMPNN [29] | 1.66 | 0.431<sub>±0.320</sub><sup>***</sup> | 0.426<sub>±0.325</sub><sup>***</sup> | 0.728<sub>±0.213</sub><sup>***</sup> | 0.575<sub>±0.289</sub><sup>***</sup> | 1.795<sub>±1.576</sub><sup>**</sup> |
| FoldX [10] | – | 0.389<sub>±0.338</sub><sup>***</sup> | 0.415<sub>±0.328</sub><sup>***</sup> | 0.699<sub>±0.218</sub><sup>***</sup> | 0.544<sub>±0.289</sub><sup>***</sup> | 1.678<sub>±1.329</sub><sup>***</sup> |
| Rosetta [9] | – | 0.364<sub>±0.327</sub><sup>***</sup> | 0.319<sub>±0.353</sub><sup>***</sup> | 0.695<sub>±0.219</sub><sup>***</sup> | 0.539<sub>±0.290</sub><sup>***</sup> | 1.600<sub>±1.282</sub><sup>***</sup> |
| Mutate Everything [38] | 658.0 | 0.371<sub>±0.380</sub><sup>***</sup> | 0.383<sub>±0.369</sub><sup>***</sup> | 0.675<sub>±0.239</sub><sup>***</sup> | 0.542<sub>±0.297</sub><sup>***</sup> | 1.489<sub>±1.004</sub><sup>***</sup> |
| ESM-IF1 [23] | 141.66 | 0.350<sub>±0.358</sub><sup>***</sup> | 0.350<sub>±0.369</sub><sup>***</sup> | 0.669<sub>±0.219</sub><sup>***</sup> | 0.495<sub>±0.278</sub><sup>***</sup> | 1.779<sub>±1.571</sub><sup>***</sup> |
| VenusREM [24] | 123.04 | 0.347<sub>±0.361</sub><sup>***</sup> | 0.351<sub>±0.376</sub><sup>***</sup> | 0.666<sub>±0.247</sub><sup>***</sup> | 0.531<sub>±0.287</sub><sup>***</sup> | 1.813<sub>±1.518</sub><sup>***</sup> |
| RSALOR [25] | – | 0.312<sub>±0.350</sub><sup>***</sup> | 0.318<sub>±0.361</sub><sup>***</sup> | 0.641<sub>±0.226</sub><sup>***</sup> | 0.483<sub>±0.280</sub><sup>***</sup> | 1.654<sub>±1.463</sub><sup>***</sup> |
| ProtSSN [26] | 2130.30 | 0.262<sub>±0.376</sub><sup>***</sup> | 0.272<sub>±0.373</sub><sup>***</sup> | 0.631<sub>±0.251</sub><sup>***</sup> | 0.506<sub>±0.281</sub><sup>***</sup> | 1.687<sub>±1.473</sub><sup>***</sup> |
| VespaG [27] | 2839.66 | 0.272<sub>±0.380</sub><sup>***</sup> | 0.268<sub>±0.389</sub><sup>***</sup> | 0.593<sub>±0.250</sub><sup>***</sup> | 0.437<sub>±0.269</sub><sup>***</sup> | 1.466<sub>±1.145</sub><sup>***</sup> |

To assess the contribution of post-training, we compared MAXWELL directly with zero-shot ProteinMPNN using the same backbone. Consistent with this distinction, MAXWELL’s calibrated landscape improved average Spearman correlation from 0.431 for zero-shot ProteinMPNN to 0.547, and improved average AUROC from 0.728 to 0.789, on the curated benchmark (Fig. 2a,b; Supplementary Tables S10 and S11). The cumulative Spearman distribution shifted to higher values after calibration, showing that the gain was broadly distributed across proteins (Fig. 2c; Supplementary Table S12).

**Fig. 2.**
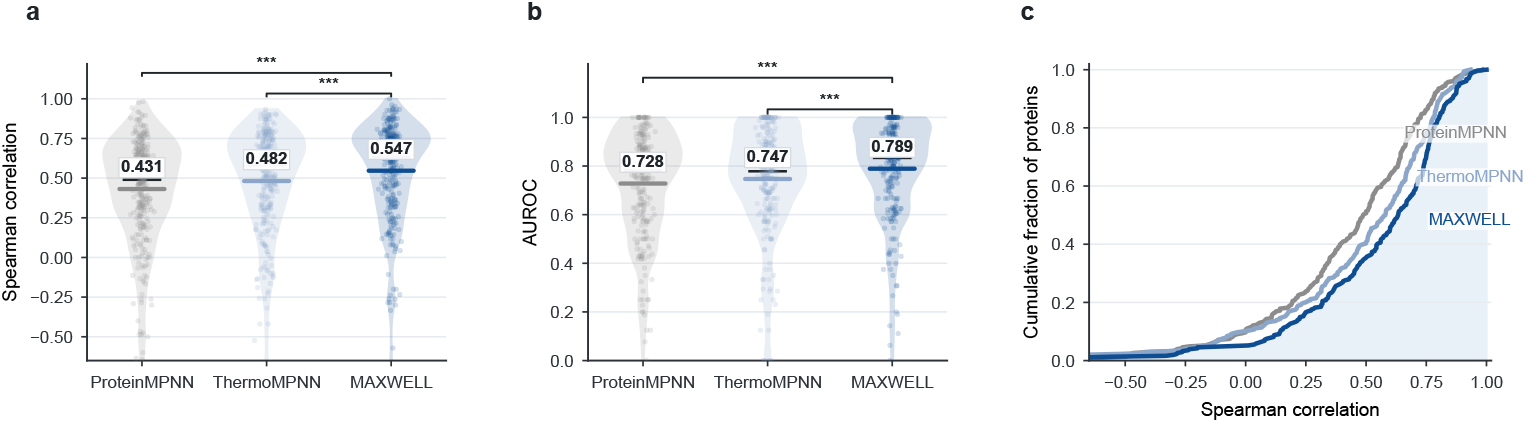
Performance comparison of ProteinMPNN before and after distribution calibration for predicting mutation effects on protein stability. **a**, Comparison of per-protein Spearman correlations for ProteinMPNN, ThermoMPNN, and calibrated MAXWELL across 308 test proteins. Violin plots show the distribution across proteins; dots represent individual proteins, thin black bars indicate medians, and colored horizontal ticks with labels indicate mean values. **b**, Comparison of stabilizing-mutation AUROC for ProteinMPNN, ThermoMPNN, and calibrated MAXWELL on proteins containing both stabilizing and non-stabilizing mutations, using *−*ΔΔ*G ≥* 0.3 kcal mol^*−*1^ as the threshold for stabilizing mutations. **c**, Cumulative distributions of per-protein Spearman correlations for ProteinMPNN, ThermoMPNN, and calibrated MAXWELL across 308 test proteins. The rightward shift after calibration indicates that sparse measured stability labels redirect the mutation preferences of ProteinMPNN toward experimentally measured effects of mutations on protein stability. Asterisks denote two-sided paired Wilcoxon tests against MAXWELL, following the convention used in table 1: ^*\**^*P <* 0.05, ^*\*\**^*P <* 0.01, and ^*\*\*\**^*P <* 0.001. Detailed numerical summaries are provided in Supplementary Tables S10 (a), S11 (b), and S12 (c); paired-test results are provided in Supplementary Tables S13 (a) and S14 (b).

The improvement was also larger than that achieved by ThermoMPNN, which uses ProteinMPNN only as a fixed feature extractor for a separately trained stability regressor: MAXWELL significantly exceeded ThermoMPNN in Spearman correlation (0.547 versus 0.482) and AUROC (0.789 versus 0.747; two-sided paired Wilcoxon tests, both *P <* 0.001; Supplementary Tables S13 and S14).

These results indicate that the main gain comes from calibrating ProteinMPNN’s native amino-acid distribution with experimental stability data to produce a mutation landscape for stability prediction.

#### Stabilizing-mutation discovery

We also evaluated MAXWELL as a global screening tool that pools and ranks all candidate mutations across the benchmark. Among the 12,102 pooled candidates, 2,085 (17.2%) were experimentally stabilizing. MAXWELL’s globally ranked top 10,top 100 and top 5% candidates were 90.0%, 86.0% and 67.8% stabilizing, respectively, corresponding to a 3.94-fold enrichment over the background stabilizing-mutation rate at the top-5% screening budget (Supplementary Table S3). Calibration also improved this discovery task relative to zero-shot ProteinMPNN: precision among the top 100 globally ranked candidates rose from 70.0% to 86.0%, and enrichment at the top-5% budget rose from 3.25-fold to 3.94-fold (Supplementary Table S4). These discovery-oriented metrics indicate that MAXWELL’s calibrated landscape concentrates true stabilizing mutations near the top of a global candidate ranking, the operating regime relevant to a fixed experimental screening budget.

### 2.3 MAXWELL reshapes ProteinMPNN preferences toward residue and biophysical stability rules

#### Substitution-pattern consistency

Pre-trained protein language models map a wild-type sequence to an amino-acid distribution, and MAXWELL calibrates this distribution on experimental data to produce the performance gains reported above. Throughout this section, the *stability* of a mutation denotes its effect on folding free energy, reported as −ΔΔ*G*, so that larger values indicate more stabilizing substitutions. Because measurements in Test12K come from multiple proteins and experimental sources, model scores and experimental values were converted to within-protein z-scores before pooling across proteins. Thus, the analyses below assess relative stability patterns rather than absolute kcal/mol calibration.

Using ProteinMPNN as a representative backbone, we assessed whether calibration makes the model output distribution reproduce these measured effects. From the curated test set Test12K, we summarized substitution behaviour as 20-by-20 mutation-preference matrices whose rows are the wild-type residue and columns are the mutant residue. Each cell reports the mean within-protein z-scored stability effect for that sub-stitution class. We constructed matched matrices for zero-shot ProteinMPNN scores, experimental −ΔΔ*G* values and MAXWELL-calibrated scores, shown in that order from left to right in Fig. 3a (Supplementary Tables S15, S16 and S17).

**Fig. 3.**
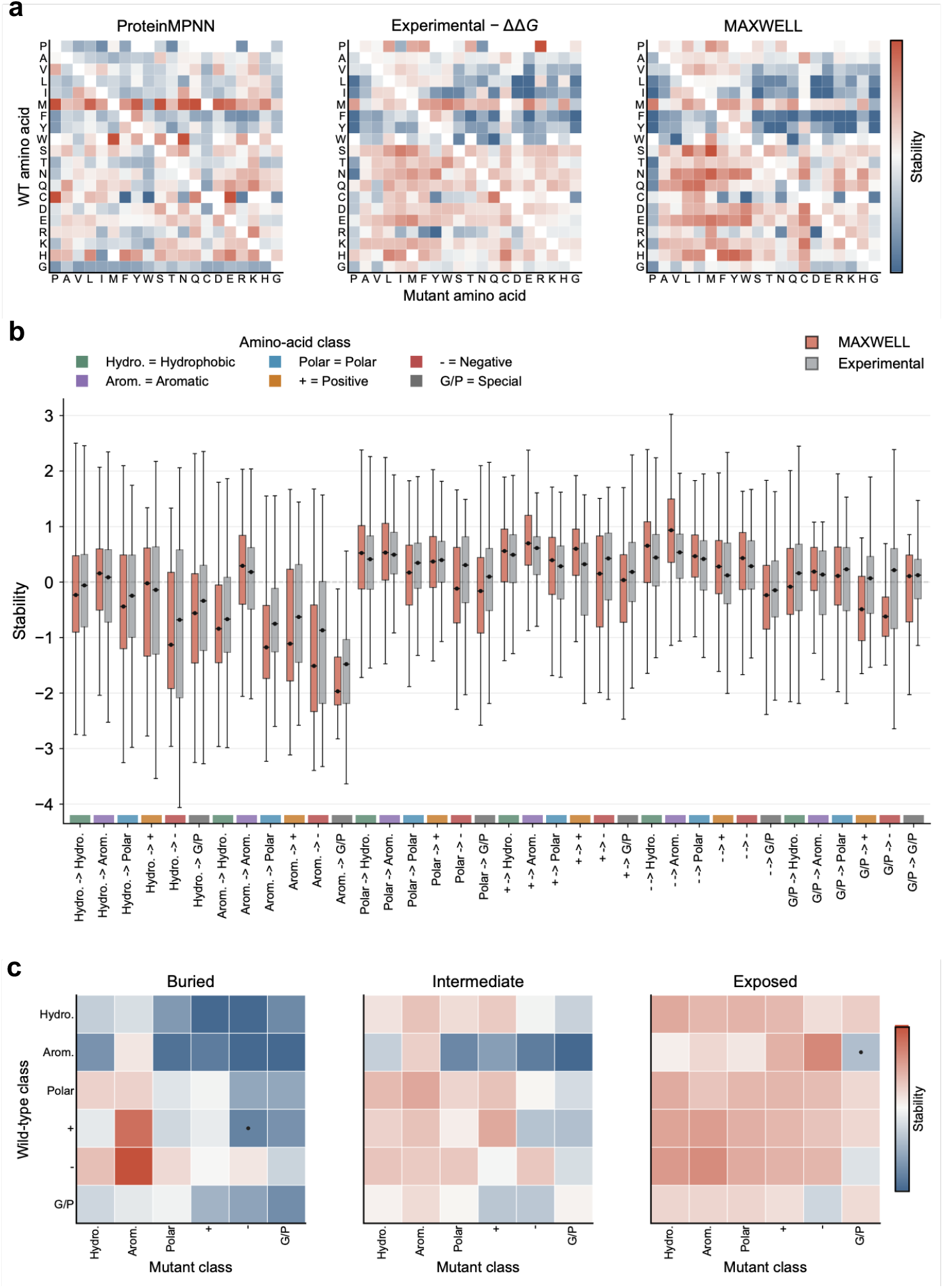
Calibrated MAXWELL distributions match experimentally measured stability patterns. **a**, Substitution preference matrices for ProteinMPNN, experiment and calibrated MAXWELL (wild-type rows, mutant columns). Complete substitution-level values are reported in Supplementary Tables S15, S16 and S17. **b**, Physicochemical class-transition distributions for calibrated MAXWELL versus experiment. **c**, Structure-context stratification across buried (*n* = 5,549), intermediate (*n* = 3,901) and exposed (*n* = 2,491) environments.

Compared with zero-shot ProteinMPNN, the MAXWELL-calibrated matrix was substantially more consistent with the experimental −ΔΔ*G* matrix. As we have shown before, per-protein Spearman correlation rose from 0.431 to 0.547 (Table 1), while the substitution-matrix level Spearman correlation increased from 0.345 to 0.776. To directly examine what calibration changed, we computed a correction matrix defined as the MAXWELL score minus the zero-shot ProteinMPNN score for each wild-type-to-mutant residue pair. We then compared this correction matrix with the experimental residual relative to ProteinMPNN. The two matrices were strongly correlated (Pearson *r* = 0.785; Spearman *ρ* = 0.808), and the same trend was also observed at the individual-mutation level (Pearson *r* = 0.604; Spearman *ρ* = 0.612). Thus, MAXWELL does not merely rescale ProteinMPNN scores; it systematically increases scores for substitution types underestimated by the zero-shot model and decreases scores for substitution types overestimated by the zero-shot model.

The wild-type proline signal illustrates this correction. ProteinMPNN scored sub-stitutions away from a native proline as destabilizing on average (mean within-protein z-score −0.66), whereas MAXWELL scored them as stabilizing on average (+0.25), in agreement with experiment. This is expected, because proline’s cyclic side chain fixes the backbone *ϕ* angle and removes the amide hydrogen-bond donor, destabilizing regular secondary structure; a strained native proline is accommodated only at a conformational cost, so replacing it relieves that cost and stabilizes the fold [39–41].

#### Physicochemical class-transition consistency

We next grouped amino acids into physicochemical classes and tested whether calibration recovered broader stability rules beyond individual residue pairs. Grouping mutations by the class of the wild-type and mutant residues (Fig. 3b), calibrated and experimental −ΔΔ*G* values were highly correlated across class transitions (Pearson *r* = 0.951; Spearman *ρ* = 0.922), whereas zero-shot ProteinMPNN showed weaker agreement (Pearson *r* = 0.424; Spearman *ρ* = 0.526). MAXWELL and experiment had matching signs for 31 of the 36 class transitions. The most destabilizing transitions were consistent with established biophysical expectations—aromatic-to-Gly/Pro, aromatic-to-charged, aromatic-to-polar and hydrophobic-to-charged—matching the expected loss of hydrophobic burial and aromatic interactions, or the introduction of charged and polar groups left uncompensated by the local electrostatic environment [42–47]. This class-level agreement provides pattern-level consistency evidence that calibration steers the pretrained distribution toward measured effects.

#### Structure-dependent mutation preferences

The analyses above show that MAXWELL calibration recovers residue-level and physicochemical stability patterns in the held-out benchmark. Finally, we examined whether the calibrated stability patterns depended on local structural environment. Residues were stratified by relative solvent accessibility into buried, intermediate and exposed environments, yielding 5,549 buried, 3,901 intermediate and 2,491 exposed substitutions in the main analysis set. For each RSA stratum, MAXWELL scores were averaged by wild-type and mutant physicochemical class to produce the structure-context heatmaps shown in Fig. 3c. The calibrated distribution shifted systematically. In buried positions, MAXWELL assigned stronger destabilizing scores to disruptive class transitions, especially substitutions from hydrophobic or aromatic residues to charged, polar or Gly/Pro residues. In contrast, exposed positions showed a broader tolerance for polar and charged substitutions, consistent with the weaker packing constraints and greater solvent buffering on protein surfaces. This pattern agrees with the classical expectation that buried protein cores are more sensitive to packing disruption and unsatisfied polar or charged groups, whereas solvent-exposed regions can accommodate such substitutions more readily [48, 49].

To quantify whether these structure-dependent MAXWELL patterns were aligned with experimental measurements, we compared MAXWELL and experimental class-transition means separately within each RSA stratum. MAXWELL retained strong agreement with experiment across all three environments, with Pearson/Spearman correlations of 0.904/0.910 in buried positions, 0.904/0.774 in intermediate positions and 0.855/0.723 in exposed positions. By comparison, zero-shot ProteinMPNN showed weaker agreement in the same environments, with Pearson/Spearman correlations of 0.601/0.588, 0.209/0.327 and 0.413/0.384, respectively.

These results show that MAXWELL effectively calibrates the probability distribution produced by the protein language model into a mutation landscape whose structure tracks mutation stability effects and varies coherently with the local structural environment, consistent with the experimentally measured stability patterns.

### 2.3 MAXWELL adapts to diverse protein language model families

#### Testing calibration across model families

MAXWELL is a model-agnostic framework that can be used to calibrate diverse protein language models; its only required interface is that a pretrained model exposes softmax-normalized amino-acid probabilities, or the corresponding log-probabilities, over the wild-type sequence. We therefore tested whether the same distribution-to-landscape calibration could be applied across four representative sources of protein-model distributions: ESM-2 [50] as a masked language model, ProGen2 [51] as an autoregressive sequence model, ProteinMPNN [29] as an autoregressive inverse-folding model and MIF-ST [52] as a masked inverse-folding model. For sequence-only models, the distribution was conditioned on the wild-type sequence; for inverse-folding models, it was additionally conditioned on the protein structure. In each case, the uncalibrated distribution provided a zero-shot mutation score, whereas MAXWELL used sparse measured stability labels to calibrate the same distribution for stability prediction. All configurations were trained on the same mega-scale cDNA-display stability source [30] and evaluated on the same curated benchmark. Model-specific batch sizes were adjusted to accommodate differences in memory requirements. We assessed both zero-shot and post-trained models using mean per-protein Spearman correlation and AUROC for stabilizing-mutation classification, following the evaluation protocol used in Table 1.

#### Calibration consistently improved every family

Across all four model families, calibration consistently improved both mutation-stability ranking and stabilizing-mutation classification (Table 2). For ESM-2, MAXWELL increased Spearman correlation by 0.194 and AUROC by 0.104, corresponding to relative gains of 84.3% and 17.2%, respectively. For ProGen2, MAXWELL increased Spearman correlation by 0.225 and AUROC by 0.075, corresponding to a Spearman sign reversal from −0.038 to 0.187 and an AUROC relative gain of 14.5%. For MIF-ST, MAXWELL increased Spearman correlation by 0.140 and AUROC by 0.091, corresponding to relative gains of 42.4% and 13.9%. For ProteinMPNN, MAXWELL increased Spearman correlation by 0.116 and AUROC by 0.061, corresponding to relative gains of 26.9% and 8.4%, while also giving the best absolute performance among the tested configurations. ProGen2’s calibrated Spearman correlation nonetheless remained the lowest among the four families (0.187), consistent with the broader observation that autoregressive sequence-likelihood models tend to underperform masked and structure-conditioned models at zero-shot protein fitness prediction [22].

**Table 2.** Pre- and post-calibration performance of MAXWELL applied to diverse protein language models. For each protein language model, Spearman correlation and AUROC were evaluated on Test12K before calibration (zero-shot) and after calibration by MAXWELL, together with the relative improvement (%). Asterisks mark paired Wilcoxon tests between calibrated and zero-shot scores within each model family, with ^*\*\*\**^*P <* 0.001.

| Model | Type | Params (M) | Spearman |  |  | AUROC |  |  |
|---|---|---|---|---|---|---|---|---|
|  |  |  | Zero-shot | MAXWELL | Improvement (%) | Zero-shot | MAXWELL | Improvement (%) |
| ESM-2<sup>1</sup> [50] | Masked language model | 650.0 | 0.230 | 0.424<sup>***</sup> | 84.3% | 0.604 | 0.708<sup>***</sup> | 17.2% |
| ProGen2<sup>2</sup> [51] | Autoregressive sequence model | 151.0 | -0.038 | 0.187<sup>***</sup> | - | 0.518 | 0.593<sup>***</sup> | 14.5% |
| ProteinMPNN<sup>3</sup> [29] | Autoregressive inverse-folding model | 1.66 | 0.431 | 0.547<sup>***</sup> | 26.9% | 0.728 | 0.789<sup>***</sup> | 8.4% |
| MIF-ST<sup>4</sup> [52] | Masked inverse-folding model | 643.4 | 0.330 | 0.470<sup>***</sup> | 42.4% | 0.653 | 0.744<sup>***</sup> | 13.9% |

<sup>1</sup>ESM-2: https://huggingface.co/facebook/esm2\_t33\_650M\_UR50D.
<sup>2</sup>ProGen2-small: https://github.com/enijkamp/progen2.
<sup>3</sup>ProteinMPNN v\_48\_020: https://github.com/dauparas/ProteinMPNN/blob/main/vanilla\_model\_weights/v\_48\_020.pt.
<sup>4</sup>MIF-ST: https://github.com/microsoft/protein-sequence-models.

Together, these results show that experimental stability measurements can improve the mutation-stability ranking of diverse protein-model backbones through the same sequence-to-landscape calibration strategy. The magnitude of the improvement nevertheless depends on the information and inductive biases available to the underlying model, with structure-conditioned backbones achieving the strongest final performance in this comparison.

#### Structure context sets the calibration ceiling

Comparing across families further showed that the two structure-conditioned models, ProteinMPNN and MIF-ST, reached higher calibrated Spearman correlations (0.547 and 0.470) than the two sequence-only models, ESM-2 and ProGen2 (0.424 and 0.187), even though ProteinMPNN required roughly two orders of magnitude fewer inference-time parameters than MIF-ST’s effective parameter budget (1.66M versus 643.4M; Table 2).

This ordering indicates that structural context sets the practical ceiling on how much a calibrated landscape can recover, echoing the parameter-efficiency argument for MAXWELL’s main ProteinMPNN instance (Table 1). Because the ProteinMPNN-based instance achieved the highest calibrated Spearman correlation and AUROC, we selected it as the release model and used it for the main benchmark, interpretation, computational-cost analysis and DhaA experimental validation.

### 2.5 A single forward pass generates a complete single-mutation stability landscape

MAXWELL uses the complete single-mutation landscape as its computational unit of prediction. For a protein of length *L*, one forward pass produces an *L* × 20 score matrix containing all 19*L* non-wild-type single substitutions. To quantify the efficiency of this formulation, we benchmarked MAXWELL against eight GPU-based baselines on seven nested truncations of one Test12K protein, spanning 64–957 residues, using a single NVIDIA RTX PRO 5000 72GB Blackwell GPU (Fig. 4a–c; Methods).

**Fig. 4.**
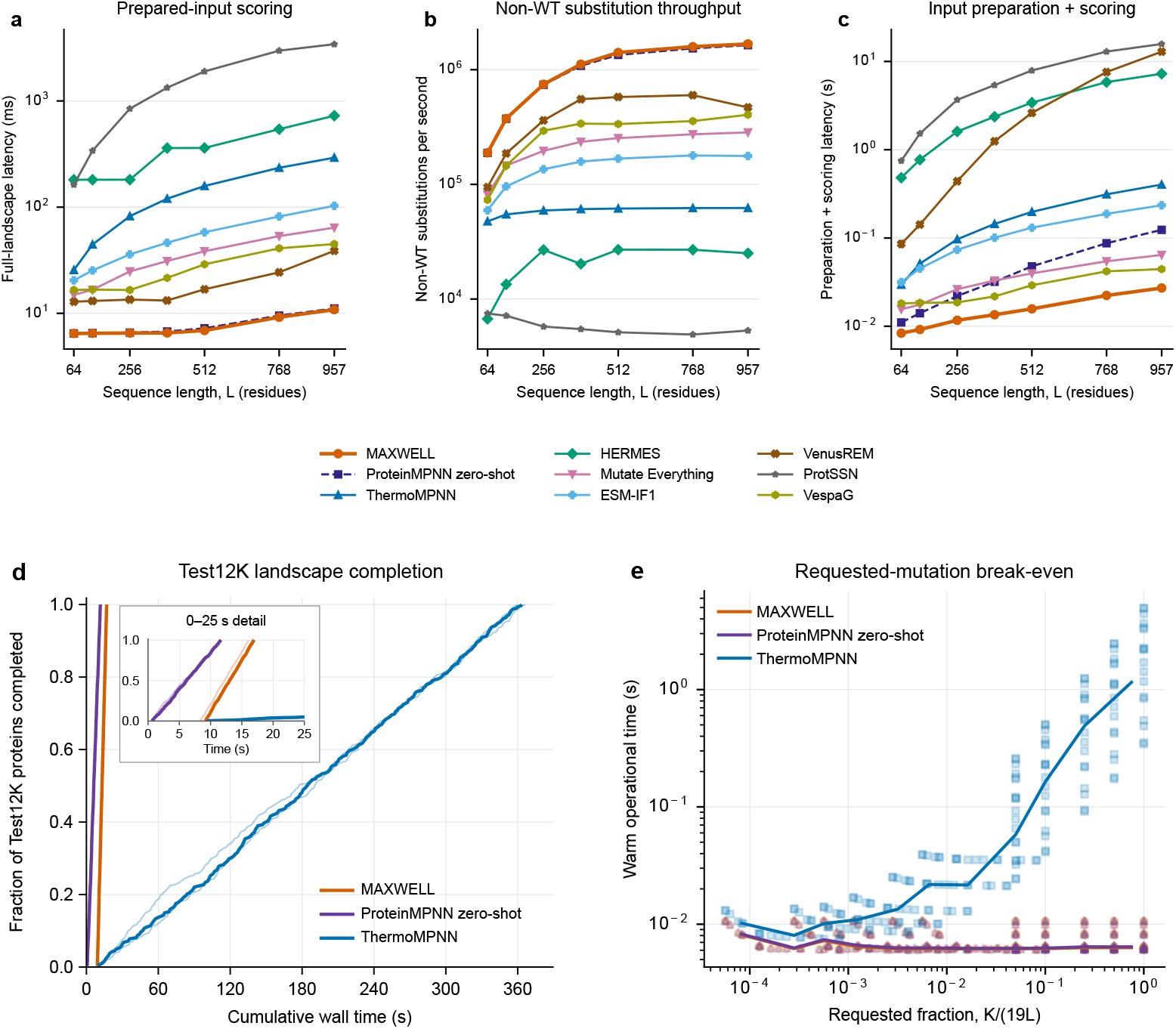
Computational efficiency of complete-landscape scoring and candidate requests. **a**, Prepared-input full-landscape scoring latency. **b**, Effective non-wild-type substitution throughput, calculated as 19*L* divided by the mean latency in a. **c**, Input preparation and scoring latency, starting from supplied sequences, structures and alignments as applicable; model loading, structure prediction and alignment search are excluded. Panels a–c compare MAXWELL with eight baselines on seven nested truncations of one protein. Points show means; error bars in a and c show sample s.d. across at least 30 and three repetitions, respectively. **d**, Cumulative completion of full landscapes for all 308 Test12K proteins, including model loading and output writing. Thin curves show three runs and thick curves show pointwise medians. The inset expands the first 25 s without changing the timing data. **e**, Model-loaded request latency versus the requested fraction *K/*(19*L*), where *K* is the number of requested substitutions. Points show per-protein, per-request-size medians across 18 proteins; curves show binned medians. Panels d and e compare MAXWELL, zero-shot ProteinMPNN and ThermoMPNN. All benchmarks used NVIDIA RTX PRO 5000 72GB Blackwell GPUs. Timing boundaries and model-specific implementations, including the different ThermoMPNN scoring interfaces in a–c and d–e, are specified in Supplementary Methods.

Across the tested lengths, MAXWELL achieved the lowest measured prepared-input scoring latency while remaining close to zero-shot ProteinMPNN (Fig. 4a). At *L* = 957, scoring all 18,183 non-wild-type substitutions required 10.84 ms for MAXWELL and 11.11 ms for zero-shot ProteinMPNN, corresponding to effective throughputs of 1.68×10^6^ and 1.64×10^6^ substitutions per second, respectively (Fig. 4b). At this length, the prepared-input latencies of the other seven baselines were 3.58–315.79 times that of MAXWELL, including 5.91-fold for Mutate Everything, 9.49-fold for ESM-IF1, 26.95-fold for ThermoMPNN and 66.93-fold for HERMES. The low computational cost was retained when model-specific input preparation was included: preparation and scoring at *L* = 957 together required 0.0271 s for MAXWELL, compared with 0.0444 s for VespaG, the fastest baseline in this timing scope (Fig. 4c). These measurements start from supplied sequences, structures and alignments as applicable, and exclude model loading, structure prediction and alignment search.

We next evaluated dataset-level landscape completion for MAXWELL, zero-shot ProteinMPNN and ThermoMPNN. Across all 308 Test12K proteins, each method produced 1,222,574 non-wild-type substitution scores (Fig. 4d). Including one-time model loading, per-protein input preparation, scoring and output writing, median completion times across three runs were 16.98 s, 11.67 s and 363.34 s, respectively. For MAXWELL and zero-shot ProteinMPNN, median processing times excluding model loading were 9.03 s and 11.45 s, respectively, showing that model loading can change their ordering in a short dataset-level task. This operational comparison used ThermoMPNN’s mutation-list interface, whereas panels a–c reused its residue-wise amino-acid outputs; timing ratios therefore remain specific to each implementation and workload.

Finally, a candidate-request benchmark across 18 proteins showed that MAXWELL’s request latency remained approximately constant as the number of requested substitutions increased (Fig. 4e), consistent with generating a complete landscape and extracting the requested entries. Together, these results show that calibration preserves the low landscape-scoring cost of the ProteinMPNN backbone and supports efficient single-site candidate prioritization under the evaluated input and hardware conditions.

### 2.6 MAXWELL transfers to melting-temperature mutation landscapes

To assess transfer across experimental readouts, we evaluated MAXWELL and 11 baselines on the complete FireProt2 Δ*T*_*m*_ benchmark, comprising 2,838 single substitutions across 86 proteins. All methods were evaluated using fixed scoring protocols and existing model checkpoints where applicable. No Δ*T*_*m*_ labels were used for model fitting, selection, or score calibration in this evaluation. Metrics were calculated within each protein and summarized across proteins, with stabilizing mutations defined as Δ*T*_*m*_ ≥ 1 °C.

Table 3 summarizes performance on this benchmark. MAXWELL achieved the highest average Spearman correlation (0.507), AUROC (0.737), and Enrich@10 (2.103). Its Pearson correlation was also numerically highest (0.485 versus 0.484 for ThermoMPNN), while its AUPRC was close to ThermoMPNN (0.578 versus 0.579), with HERMES achieving the highest AUPRC (0.597).

**Table 3.** Performance evaluation on melting-temperature mutation landscapes. We benchmarked MAXWELL and 11 baseline methods on the complete FireProt2 Δ*T*_*m*_ dataset, comprising 2,838 mutations across 86 proteins. Values are per-protein means *±* sample standard deviations across eligible proteins. Stabilizing mutations were defined as Δ*T*_*m*_ *≥* 1 ^*°*^C. Enrich@10 measures fold-enrichment among the top-10 ranked candidates relative to the background stabilizing-mutation rate. Bold means with *†* mark the highest mean in each column, and *‡* marks the second-highest mean.

| Method | Spearman↑ | Pearson↑ | AUROC↑ | AUPRC↑ | Enrich@10↑ |
|---|---|---|---|---|---|
| MAXWELL | **0.507**±0.359<sup>†</sup> | **0.485**±0.403<sup>†</sup> | **0.737**±0.218<sup>†</sup> | 0.578±0.287 | **2.103**±1.598<sup>†</sup> |
| ThermoMPNN | 0.467±0.372<sup>‡</sup> | 0.484±0.371<sup>‡</sup> | 0.733±0.206<sup>‡</sup> | 0.579±0.288<sup>‡</sup> | 1.988±1.153 |
| HERMES | 0.431±0.371 | 0.439±0.394 | 0.716±0.234 | **0.597**±0.295<sup>†</sup> | 2.044±1.428<sup>‡</sup> |
| Rosetta | 0.390±0.377 | 0.370±0.415 | 0.703±0.227 | 0.577±0.306 | 1.630±1.105 |
| Mutate Everything | 0.337±0.361 | 0.352±0.371 | 0.648±0.260 | 0.535±0.300 | 1.528±1.115 |
| FoldX | 0.337±0.340 | 0.336±0.366 | 0.630±0.234 | 0.509±0.299 | 1.606±1.198 |
| ProteinMPNN (zero-shot) | 0.361±0.385 | 0.339±0.398 | 0.687±0.238 | 0.572±0.285 | 1.971±1.322 |
| ProtSSN (zero-shot) | 0.281±0.386 | 0.280±0.394 | 0.643±0.231 | 0.547±0.299 | 1.612±1.106 |
| VenusREM (zero-shot) | 0.340±0.369 | 0.339±0.411 | 0.651±0.242 | 0.541±0.312 | 1.562±1.147 |
| RSALOR (zero-shot) | 0.300±0.359 | 0.324±0.389 | 0.602±0.214 | 0.480±0.282 | 1.397±1.084 |
| VespaG (zero-shot) | 0.268±0.365 | 0.274±0.398 | 0.595±0.249 | 0.477±0.292 | 1.412±1.080 |
| ESM-IF1 (zero-shot) | 0.287±0.358 | 0.289±0.397 | 0.625±0.234 | 0.518±0.275 | 1.830±1.674 |

To isolate the contribution of calibration, we compared MAXWELL with zero-shot ProteinMPNN within each of the 86 proteins. Calibration increased mean Spearman correlation from 0.361 to 0.507, with a mean paired improvement of 0.146 (95% protein-bootstrap confidence interval, 0.076–0.219; Fig. 5). AUROC also increased from 0.687 to 0.737. This within-protein comparison supports transfer of the calibration gain to the melting-temperature readout, rather than an improvement arising from differences in the proteins evaluated.

**Fig. 5.**
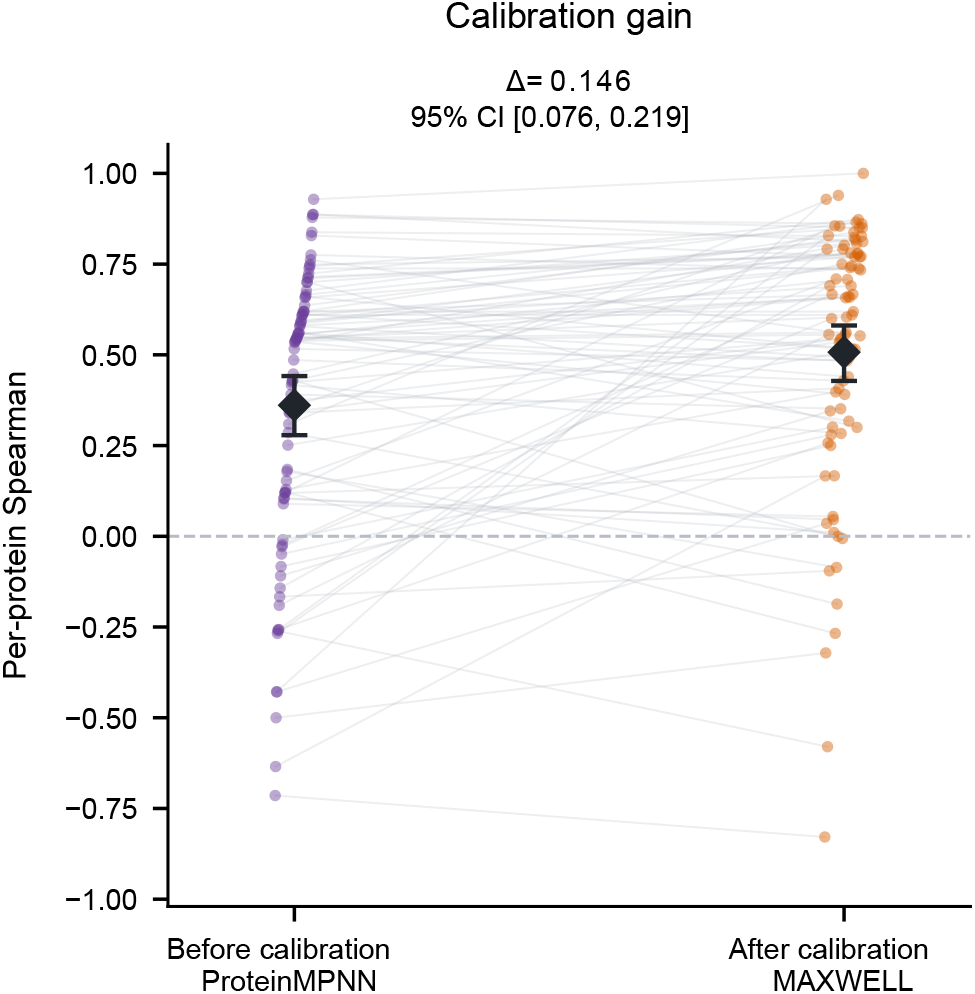
Stability calibration improves mutation ranking on melting-temperature land-scapes. Paired per-protein Spearman correlations for zero-shot ProteinMPNN and calibrated MAXWELL on 2,838 measured single substitutions across 86 proteins. Lines connect the same protein; diamonds and error bars indicate unweighted means and 95% protein-bootstrap confidence intervals. The annotated mean difference and its interval were obtained by paired resampling of proteins. No Δ*T*_*m*_ labels were used for fitting, model selection, or score calibration in this evaluation.

Sensitivity analyses using stabilizing-mutation thresholds of 0.5, 1, and 2 °C yielded macro AUROC values of 0.750, 0.737, and 0.748, respectively (Supplementary Table S24), indicating that the classification performance was not restricted to a single threshold. Together, these results support the use of the calibrated landscape for prioritizing mutations by thermal-stability effects without Δ*T*_*m*_-specific fitting, while not establishing prediction of absolute Δ*T*_*m*_ values.

### 2.7 Experimental evaluation of MAXWELL-prioritized DhaA mutants

Stability constrains whether an engineered protein can be produced, stored and deployed under real operating conditions [1–3], so experimentally validated stability gains carry practical significance beyond computational benchmarks. To test whether a MAXWELL-calibrated mutation landscape translates into such gains, we applied MAXWELL to DhaA, a bacterial haloalkane dehalogenase that also serves as the enzyme scaffold for HaloTag protein-labeling technology [36, 53], as a single-mutation wet-lab validation target. Starting from the wild-type DhaA structure (PDB ID: 4HZG), MAXWELL generated a position-by-amino-acid landscape of predicted stability effects and ranked candidate substitutions by predicted stability. We selected the top ten highest-ranked single-site variants, constructed and purified the corresponding proteins, and measured their melting temperatures against the wild-type baseline; wild-type DhaA had a mean *T*_*m*_ of 54.92 °C across three measurements (Supplementary Table S5).

Seven of the ten tested variants showed higher mean *T*_*m*_ values than wild type, with four variants reaching statistical significance in a two-sided Welch’s t-test based on three replicate measurements per protein (Fig. 6b; Supplementary Table S6). The strongest stabilizing variant was G171W, which increased *T*_*m*_ by 4.91 °C, from 54.92 to 59.83 °C. Two E20 substitutions, E20Y and E20F, produced more moderate increases of 1.95 and 1.74 °C, respectively, and K236F produced a smaller gain of 0.79 °C. Other substitutions at K236 showed small, non-significant mean increases, whereas T148W was approximately neutral and K236V and H84F were destabilizing under this assay. Relative-activity measurements further revealed a trade-off between thermal improvement and catalytic activity. G171W retained 44.2% activity relative to wild type, whereas E20Y and E20F retained 93.0% and 85.7% activity, respectively, while still significantly increasing *T*_*m*_ (Supplementary Table S6).

**Fig. 6.**
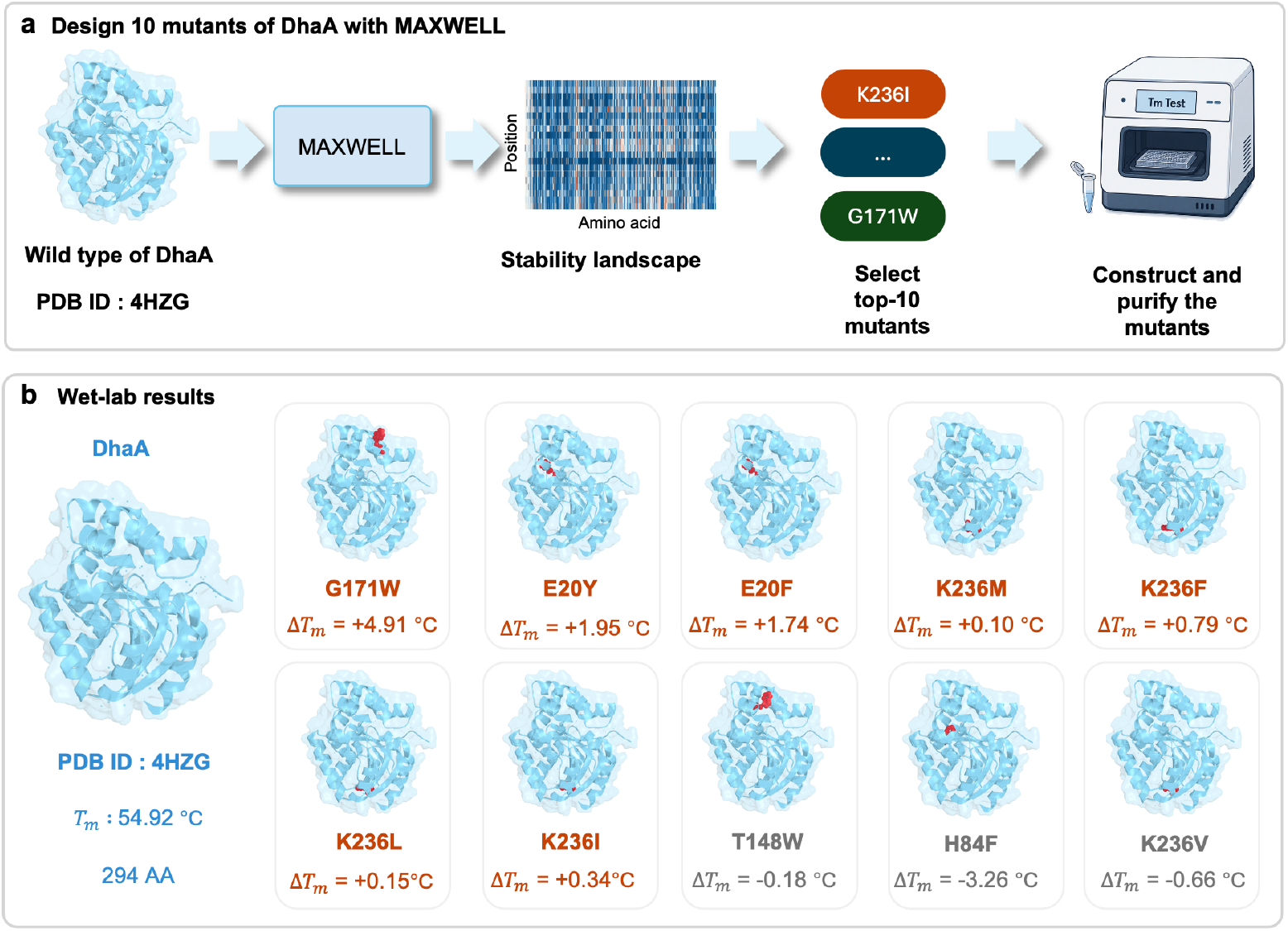
Wet-lab design and validation of MAXWELL-prioritized DhaA variants. **a**, MAXWELL ranks single-mutation stability effects from wild-type DhaA (PDB ID: 4HZG) and selects ten variants for *T*_*m*_ measurement (Supplementary Table S5). **b**, Measured *T*_*m*_ for wild-type DhaA and mean Δ*T*_*m*_ values of the ten designed variants relative to wild type (Supplementary Table S6).

These results show that the MAXWELL-predicted DhaA landscape can nominate experimentally testable variants with increased mean thermal stability. In particular, G171W provided the largest observed increase in *T*_*m*_, whereas E20Y and E20F showed a more favorable balance between increased *T*_*m*_ and retained activity. Together with the computational benchmark and biophysical analyses above, this wet-lab outcome closes the loop from calibrated prediction to experimental validation, supporting MAXWELL as a practical tool for concentrating limited experimental resources on a small, enriched candidate set before broader engineering campaigns.

## 3 Discussion

MAXWELL reframes mutation-stability prediction as a calibration problem rather than a representation-transfer problem. Most supervised stability predictors treat pretrained protein language models as fixed feature extractors and learn a separate scoring head on top of their embeddings [28]. By contrast, MAXWELL supervises the model’s native position-by-amino-acid output directly, converting it into a wild-type-normalized mutation landscape whose entries provide stability-oriented scores for single-site variants. This design choice enables MAXWELL to leverage sequence and structural information learned during pretraining more directly than representation-transfer approaches (Table 1). Rather than adding a separately trained scoring network on top of frozen embeddings, MAXWELL supervises ProteinMPNN’s native position-by-amino-acid output and calibrates it with sparse stability labels, redirecting the pretrained distribution toward measured thermostability.

The benchmark and biophysical analyses together suggest that calibration works by steering an existing distribution, not by learning stability from scratch. Across four protein language model families, calibration consistently improved ranking and stabilizing-mutation classification relative to each model’s zero-shot score (Table 2), and models with stronger zero-shot performance yielded stronger calibrated land-scapes. Structure-conditioned backbones reached the highest calibrated performance, consistent with structural information providing useful signal for mutation-stability prediction. The substitution-pattern, class-transition and solvent-exposure analyses (Fig. 3) further indicate that the calibrated landscape is consistent with established biophysical mutation trends at the distribution level, which supports interpreting MAXWELL outputs as stability-prioritized design suggestions rather than opaque regression scores.

The DhaA experiment provides an initial experimental assessment of whether MAXWELL-predicted variants can exhibit improved thermal properties. Seven of ten model-prioritized single-site variants increased *T*_*m*_ relative to wild type, including G171W with a 4.91 °C gain (Fig. 6b). This outcome suggests that a calibrated land-scape can concentrate limited wet-lab effort on a small, high-yield candidate set before activity, expression and manufacturability assays. Because the most thermostable variant is not always the most useful one, MAXWELL is best viewed as a stability-prioritization tool that narrows the search space rather than as a final design selector. Variants that show reproducible improvements after independent experimental replication could provide a seed set for downstream combinatorial design. Individually validated substitutions can be combined with specialized multi-point methods when higher-order variants are the target.

These screening benefits extend beyond a single dehalogenase scaffold. Enzyme engineering often requires stability across wide operating ranges [1, 2], and biologics development requires expression, storage and manufacturability guarantees [3, 5, 6]. A calibration-ranked candidate set could therefore reduce experimental burden in both settings. The sequence-to-landscape formulation also lowers computational cost for saturation-mutagenesis planning, because one forward pass yields the full position-by-amino-acid landscape rather than one score per substitution (Fig. 4; Supplementary Tables S18, S19 and S20).

The current framework is scoped to single-point mutation landscapes and does not model epistasis among simultaneous substitutions. Specialized combinatorial-mutation methods therefore remain important when multi-site epistasis is the primary design objective [54–56]. Extending calibration from single substitutions to controlled multi-mutation settings, where epistatic interactions are learned explicitly, is a natural next step. More broadly, the calibration principle may apply wherever pretrained protein models expose amino-acid probability distributions and sparse experimental labels can redirect those outputs toward a downstream fitness objective.

## 4 Methods

### 4.1 Designing Stable Mutations via Protein Stability Prediction

Identifying stabilizing mutations is a central objective in protein engineering. Because experimental validation of a large number of variants is resource-consuming, computational stability prediction can be used to select potential candidates for subsequent validation. Let **A** = (*a*_1_, …, *a*_*n*_) denote a wild-type protein sequence of length *n*, and let *A* denote the standard amino acid alphabet. We represent the predicted effects of all allowable single-point mutations by a mutation stability landscape **L** ∈ ℝ^*n*×|*A*|^, where each row corresponds to a sequence position, each column corresponds to a candidate amino acid, and the entry *L*_*i,a*_ quantifies the predicted stability change associated with substituting the wild-type residue *a*_*i*_ at position *i* with amino acid *a*. The wild-type substitution may be excluded or assigned a neutral value. Given such a landscape, candidate mutations can be directly compared according to their predicted stability effects. The full mutation space can then be ranked, and the mutations predicted to be most stabilizing can be selected for experimental evaluation. Our goal is therefore to learn a function that maps the wild-type sequence **A**, together with structural information **S** when available, to a predicted mutation stability landscape: 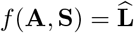. Given this landscape, mutations can be compared according to their stability effects, allowing the full mutation space to be ranked and the most stabilizing candidates to be selected for experimental testing.

### 4.2 Protein language models enable zero-shot protein mutation-induced stability change prediction

MAXWELL starts from a pretrained protein language model that exposes amino-acid probabilities or logits aligned to protein positions. We view such a model as a parameterized function *f*_*θ*_ that maps a wild-type protein sequence **A** = (*a*_1_, …, *a*_*n*_) to a position-by-amino-acid distribution matrix

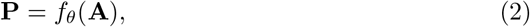

where **P** ∈ ℝ^*n*×|*A*|^ and **P**_*ij*_ is the probability or normalized score of amino acid *j* at position *i*. For structure-conditioned models, the same interface includes a structural input **S**,

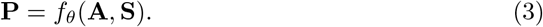

This notation covers sequence-only protein language models and inverse-folding models; **S** is omitted when the underlying model does not use structure.

Protein language models can be used for mutation scoring because their training distributions are dominated by naturally occurring proteins shaped by evolutionary selection. For a mutation at position *i* from the wild-type amino acid *a*_*i*_ to amino acid *j*, the standard zero-shot score compares the mutant and wild-type log-probabilities:

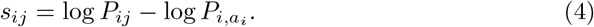

This log-probability-ratio form follows the standard zero-shot mutation-scoring interface used for protein language models and inverse-folding models [21, 23, 57]. The score is not necessarily a direct stability value for every model, because it can also reflect activity, expression, structural compatibility or other evolutionary constraints. For inverse-folding models, however, this log-likelihood ratio has a direct free-energy interpretation related to ΔΔ*G* [57].

### 4.3 Stability landscape construction

For a wild-type protein of length *n*, we define its single-mutation ΔΔ*G* landscape as a real-valued matrix **D** ∈ ℝ^*n*×|*A*|^. Entry *D*_*ij*_ denotes the measured ΔΔ*G* value produced by replacing the wild-type residue at position *i* with amino acid *j*. To make the direction of the learning target consistent with the MAXWELL score, we define the stability-oriented target landscape **Y** = −**D**, such that **Y**_**ij**_ *>* 0 indicates stabilization and larger **Y**_**ij**_ indicates a more stabilizing mutation. Because experimental assays usually measure only a small fraction of all possible substitutions, this landscape is sparse. We record the observed entries with a binary mask **M** ∈ {0, 1}^*n*×|*A*|^, where *M*_*ij*_ = 1 if *Y*_*ij*_ is experimentally available and *M*_*ij*_ = 0 otherwise.

### 4.4 Matrix-wise landscape learning

MAXWELL reformulates the zero-shot mutation score in equation (4) as a differentiable matrix operation over the full position-by-amino-acid landscape. Given **A** and,when available, **S**, the pretrained model first produces **P** through equation (2) or equation (3). We then convert the model output to a log-probability or log-softmax-normalized score matrix

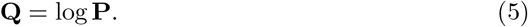

Let **X** ∈ {0, 1}^*n*×|*A*|^ be the one-hot encoding of the wild-type sequence. For each position *i*, only the entry corresponding to the wild-type amino acid *a*_*i*_ is set to one. The Hadamard product between the log-probability matrix and the one-hot matrix extracts the wild-type log-probability at each position. The resulting wild-type log-probability vector is

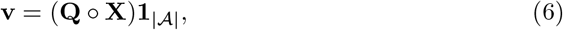

where ° denotes element-wise multiplication and **1**_|*A*|_ is an all-one vector.

We replicate **v** across amino-acid identities and subtract it from **Q** to obtain the predicted mutation landscape:

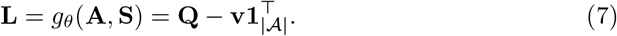

Equivalently,

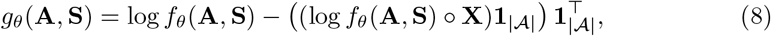

with the structural argument omitted for sequence-only models. Each entry *L*_*ij*_ is the log-probability difference between amino acid *j* and the wild-type amino acid at position *i*, so equation (7) is the matrix form of equation (4). Thus, a single model call produces the full single-mutation landscape. Because *g*_*θ*_ is differentiable whenever *f*_*θ*_ is differentiable, sparse ΔΔ*G* labels can update the pretrained model through backpropagation while retaining the native protein-model distribution interface.

### 4.5 Pearson landscape objective

The training objective compares the predicted landscape with the measured ΔΔ*G* landscape only at observed entries. For each protein landscape, we use a masked Pearson correlation loss:

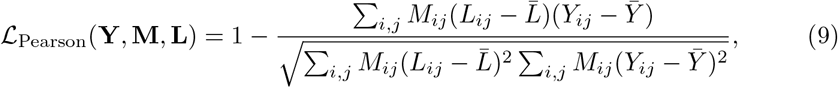

where 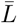 and 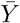 are the mean predicted and measured stability scores over entries with *M*_*ij*_ = 1.

Across a training set *D* of sparse protein landscapes, the optimized objective is

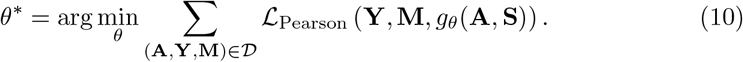

This Pearson-only objective is the sole training loss and aligns within-protein mutation preferences directly. It keeps unmeasured landscape entries available for prediction while restricting the training signal to experimentally observed mutations. We also evaluated a listwise ranking alternative under the same training setup; it did not improve validation or benchmark performance relative to the Pearson-only objective (Supplementary Table S1). The full training loop and prediction procedure are described in Supplementary Methods, Sequence-to-landscape learning algorithm (Supplementary Section 1.1).

### 4.6 Training dataset and benchmark setup

#### Test set

The test set aggregates mutation ΔΔ*G* measurements currently available from multiple published sources, including p53, myoglobin and Ssym [31], S669 [32], S8754 and M1261 [20], vb1432 [33], FireProtDB [34] and ThermoMutDB [35]. When duplicate measurements existed for the same mutation, we retained the ΔΔ*G* value with the largest absolute magnitude. This choice was intended to reduce neutral-label bias and prevent predictions from collapsing toward zero. After deduplication, the test set contained 12,443 mutations across 308 proteins, forming 308 sparse single-mutation ΔΔ*G* landscapes. The main benchmark comparison used a filtered subset of 12,102 mutations for which comparable predictions were available across evaluated methods. We refer to this benchmark as Test12K. Records were organized by wild-type protein to enable per-protein prediction and evaluation, consistent with benchmarks such as ProteinGym [22]. Only proteins with at least five measured mutations were retained so that per-protein analyses remained meaningful.

#### Training set

The training set was derived from cDNA272K, a mega-scale cDNA-display stability dataset containing 272K protein mutation sequences [30]. This resource is widely used for protein stability prediction [58]. To limit data leakage, we removed from the training set all sequences sharing at least 30% sequence identity with any Test12K sequence using MMseqs2 [59]. We chose the 30% threshold because it lies within the “twilight zone” in which 95% of protein pairs have distinct structural folds [60]. The resulting filtered training set, Train226K, contained 226K mutation sequences with less than 30% sequence identity to any Test12K sequence. All datasets were stored in a protein-specific format to facilitate model training and evaluation. Sequences requiring structural input for structure-conditioned models were folded with ColabFold 2.5 [61].

To further verify separation between the training and test sets, we performed pairwise global sequence alignment with the Needleman-Wunsch algorithm [62]. The resulting similarity distribution had a mean of 6.17%, and more than 88% of test-training pairs showed less than 10% sequence identity. This low overlap indicates that sequence-based deduplication effectively limits leakage and supports generalization to unseen sequences.

#### 4.7 Training protocol and hyperparameter search

MAXWELL models were trained with the Adam optimizer [63]. All hyperparameters were selected on Train226K without using Test12K. The training loss was the masked Pearson objective defined in equation (10).

To prevent test-set leakage, we performed five-fold cross-validation at the protein-landscape level on Train226K. Test12K proteins were excluded from this procedure and reserved for final evaluation only. Each wild-type protein and all of its measured mutations were assigned to a single fold, so no mutation landscape was split across internal training and validation sets. The 255 training protein landscapes were partitioned into five disjoint folds, denoted *T*_1_, …, *T*_5_. For each fold index *i*, the validation set was *T*_*i*_ and the internal training set was *D*_train_ \ *T*_*i*_, comprising the remaining four folds. For each candidate hyperparameter setting, we trained on the internal training set, evaluated on the corresponding validation fold and repeated this procedure for all five folds. We recorded the best mean per-protein Spearman correlation on each validation fold and averaged these five values; the setting with the highest mean validation Spearman correlation was selected. During each fold, training was stopped early if validation performance did not improve for five consecutive epochs.

For the main ProteinMPNN-based MAXWELL model, we searched batch size and learning rate over batch sizes {1, 2, 4, 8, 16, 32} and learning rates {1 × 10^−5^, 5 ×10^−5^, 1 × 10^−4^, 5 × 10^−4^, 1 × 10^−3^}. The highest validation Spearman correlation was achieved at batch size 16 and learning rate 5×10^−4^. Based on this cross-validation, the final ProteinMPNN-based model used these settings. After hyperparameter selection, a single model was trained on the complete Train226K dataset for the selected number of epochs and evaluated once on Test12K. The full grid-search results are reported in Supplementary Table S2.

#### 4.8 Evaluation metrics and baselines

We compared MAXWELL against representative baselines on Test12K using a filtered benchmark table of 12,102 mutations across 308 proteins (section 4.6). All methods were evaluated on this shared table under a common protocol. Predictions were aligned to the measured ΔΔ*G* landscapes defined in section 4.3. Model scores were oriented so that higher values indicated stronger predicted stabilization. The same metric implementation was applied to every method. For released checkpoints or author-provided prediction files, raw outputs were recomputed on the filtered table rather than importing literature summary values. To keep supervised comparisons fair, neural predictors were restricted where possible to models trained from the same stability-data source and evaluated with fixed weights on the test proteins.

The compared methods spanned three classes. Physics-based energy models comprised FoldX [10] and Rosetta Cartesian mutation scoring implemented with PyRosetta [9]. The Zero-shot baselines comprised ProteinMPNN [29], ESM-IF1 [23], VenusREM [24], RSALOR [25], ProtSSN [26] and VespaG [27]. Supervised neural predictors comprised ThermoMPNN [28], HERMES [37] and Mutate Everything [38]. Methods excluded from the main comparison and literature-only reference values are listed in Supplementary Information. ESM-IF1 scores substitutions from structure-conditioned amino-acid probabilities. VenusREM integrates a structure-aware protein language model with evolutionary information retrieved from homologous sequences. RSALOR combines position-specific evolutionary log-odds with relative solvent accessibility. ProtSSN combines protein language model representations with an equivariant graph encoder, whereas VespaG uses an ESM-2-based prediction head trained on GEMME-derived pseudo-labels. Method-specific implementations and scoring configurations are described in Supplementary Methods.

Let *P* denote the set of test proteins. For each protein *p* ∈ *P*, let *n*_*p*_ be the number of measured mutations with predicted scores *ŝ*_*p,i*_ and measured −ΔΔ*G* values *y*_*p,i*_. Stabilizing mutations were defined as *y*_*p,i*_ ≥ 0.3 kcal/mol and were encoded by indicators *z*_*p,i*_ ∈ {0, 1}.

The primary benchmark metric was mean Spearman correlation averaged across all test proteins,

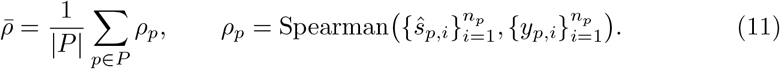

This averaging prevents proteins with many measured mutations from dominating the aggregate score. Although training optimized the differentiable masked Pearson objective in equation (10), Spearman correlation was used for benchmark reporting and hyperparameter selection. Mutation prioritization depends on within-protein rank order rather than absolute scale. Spearman correlation is not differentiable and therefore cannot serve directly as a training loss.

Pearson correlation was computed in the same way by replacing *ρ*_*p*_ with the within-protein Pearson correlation *r*_*p*_,

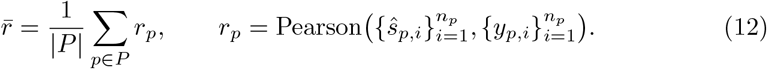

For each protein with both stabilizing and non-stabilizing mutations, we computed

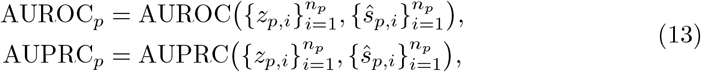

and reported the mean AUROC and AUPRC across eligible proteins. For Enrichment@10, measured mutations within each protein were ranked by *ŝ*_*p,i*_ and

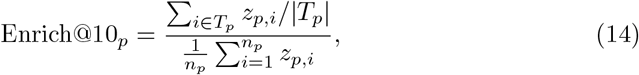

where *T*_*p*_ contains the ten highest-scoring measured mutations in protein *p*. The reported Enrichment@10 is the mean of Enrich@10_*p*_ averaged across all test proteins with at least ten measured mutations. Table 1 reports 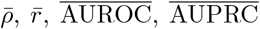 and mean Enrichment@10 on the filtered benchmark table. Full definitions of additional secondary and ranking metrics are provided in Supplementary Methods, Evaluation metrics (Supplementary Section 1.4). Length-stratified baseline analyses are reported in Supplementary Table S8.

#### Evaluation on melting-temperature changes

We evaluated MAXWELL and eleven baselines on the same 2,838 measured single-point mutations from 86 proteins in the FireProt2 Δ*T*_*m*_ dataset. Experimental Δ*T*_*m*_ values and prediction scores were oriented so that higher values indicated greater stabilization. Metrics were calculated within each protein and summarized by their unweighted mean and sample standard deviation (denominator *n* − 1). Spearman and Pearson correlations were defined for 86 proteins for each method except RSALOR, for which 85 proteins were included because one protein had constant predictions. Stabilizing mutations were defined as Δ*T*_*m*_ ≥ 1 °C. AUROC and AUPRC were averaged across the 66 proteins containing both stabilizing and non-stabilizing mutations; AUPRC was computed as average precision using average_precision_score. For Enrich@10, measured mutations were ranked within each protein by descending prediction score, and the stabilizing-mutation fraction among the top 10 was divided by that protein’s overall stabilizing-mutation fraction. This metric included the 49 proteins with at least 10 measured candidates and at least one stabilizing mutation. Proteins with an undefined metric were excluded only from that metric’s summary. Sensitivity analyses repeated the same procedure with thresholds of 0.5 and 2 °C. No Δ*T*_*m*_ labels were used to fit, select, or calibrate models during this evaluation. Model scores were assessed for association with and ranking of experimental Δ*T*_*m*_ values; absolute Δ*T*_*m*_ prediction was not evaluated. For the before–after calibration comparison, Spearman correlations were paired within proteins. Confidence intervals for the unweighted mean correlations and the mean paired difference were obtained from 10,000 bootstrap resamples of the 86 proteins, using the 2.5th and 97.5th percentiles; each paired resample retained both predictions for the same protein.

### 4.9 Biophysical consistency analysis

To test whether the calibrated MAXWELL landscape reproduces established sequence- and structure-dependent stability trends, we analyzed ProteinMPNN-based MAXWELL scores on Test12K, retaining single-point substitutions after excluding self-mutations and mutations with a measured ΔΔ*G* of exactly zero. Amino acids were grouped into six physicochemical classes (hydrophobic: A, V, I, L, M; aromatic: F, Y, W; polar: S, T, N, Q, C; positively charged: K, R, H; negatively charged: D, E; and glycine/proline: G and P, grouped in a separate class because of their distinct effects on backbone conformation), and MAXWELL scores, zero-shot Protein-MPNN scores and experimental −ΔΔ*G* values were each converted to within-protein z-scores before pooling across proteins. Structural context was defined by relative solvent accessibility (RSA), computed with the Shrake–Rupley rolling-ball algorithm in Biopython on each folded or experimentally determined structure [64] and normalized by the corresponding maximum residue accessible surface area [65]; residues were stratified as buried (RSA ≤ 0.20), intermediate (0.20 *<* RSA ≤ 0.50) or exposed (RSA *>* 0.50). For substitution-pattern analysis, pooled mutations were grouped by wild-type and mutant amino acid to form matched 20-by-20 matrices reporting the mean within-protein z-score in each cell. For class-transition analysis, mutations were summarized into the 36 wild-type-class to mutant-class transitions; transition-level mean z-scores were compared between calibrated MAXWELL and experiment using Pearson and Spearman correlation, with sign agreement counted at the transition level. For structure-context analysis, calibrated values were averaged within each RSA stratum by wild-type and mutant physicochemical class to form matched six-by-six class-transition heatmaps for buried, intermediate and exposed environments. Additional filtering rules, sample counts and plotting thresholds are provided in Supplementary Methods, Biophysical consistency analyses (Supplementary Section 1.8); complete substitution-level matrix values underlying Fig. 3a are reported in Supplementary Tables S15, S16 and S17.

### 4.10 DhaA experimental validation

#### Mutant selection procedure

We used MAXWELL to prioritize single-point mutations in DhaA dehalogenase for wet-lab validation. The wild-type DhaA structure (PDB entry 4HZG) was used as the structural input for the ProteinMPNN-based MAXWELL instance. MAXWELL generated a full position-by-amino-acid landscape of predicted stability effects for all possible single amino-acid substitutions. Candidate mutations were ranked by the calibrated landscape entries *L*_*ij*_ in equation (7). We selected the ten highest-scoring single-site variants without per-position deduplication, yielding G171W, E20Y, E20F, K236M, K236F, K236L, K236I, T148W, H84F and K236V. The complete ranked list with predicted scores is provided in Supplementary Table S5. Candidate selection was determined solely by this model ranking before wet-lab measurements, and thermal-stability and activity assays served only for post hoc validation.

#### Plasmid construction

Codon-optimized coding sequences for wild-type DhaA and the selected variants were synthesized and cloned into the pET-28a (+) expression vector between the NdeI and XhoI restriction sites, placing each construct in frame with an N-terminal hexahistidine tag for protein purification. All expression constructs were confirmed by Sanger sequencing before transformation.

#### Protein expression

Confirmed constructs were transformed into *Escherichia coli* BL21(DE3) cells. Cultures were grown in LB medium supplemented with kanamycin at 50 µg ml^−1^ to an optical density at 600 nm of 0.8. Protein expression was induced with 0.5 mM isopropyl *β*-D-1-thiogalactopyranoside, and cultures were incubated at 16 °C for 20 h. Cells were harvested by centrifugation at 3345 × g for 30 min.

#### Protein purification

Cell pellets were resuspended in lysis buffer (25 mM Tris-HCl, 500 mM NaCl, pH 8.0) and disrupted by ultrasonication (Scientz, China). The lysates were centrifuged at 17,418 × g for 30 min at 4 °C, and the supernatants were loaded onto Ni-NTA columns (Ni NTA Beads 6FF, SA005500; Smart Lifesciences, China) pre-equilibrated with lysis buffer (25 mM Tris-HCl, 500 mM NaCl, pH 8.0). Bound proteins were washed with buffer containing 50 mM imidazole and eluted with buffer containing 250 mM imidazole. Eluted proteins were concentrated and buffer-exchanged into assay buffer (50 mM HEPES, 150 mM NaCl, pH 7.5), using Amicon Ultra centrifugal filters (10 kDa MWCO; Millipore) through repeated dilution and centrifugation cycles at 3345 × g for 25 min at 4 °C. Protein concentration was determined from absorbance at 280 nm using a NanoDrop 2000c spectrophotometer. Purity of representative variants was assessed by SDS–PAGE (Supplementary Fig. S1).

#### Melting-temperature assessment

Melting temperatures were determined by differential scanning fluorimetry (DSF) using the Protein Thermal Shift Dye Kit (Thermo Fisher, USA). SYPRO Orange dye (SUPELCO, USA) was first diluted 50-fold by mixing 1.0 µl dye with 49 µl assay buffer (50 mM HEPES, 150 mM NaCl, pH 7.5). For each reaction, 1 µl of diluted dye was added to 19 µl purified protein at 0.1 mg ml^−1^, giving a final reaction volume of 20 µl. DSF measurements were performed on a LightCycler 480 Instrument II (Roche, USA). Samples were equilibrated at 25 °C and then heated to 99 °C at 0.05 °C s^−1^, followed by a 2 min hold at the final temperature. Melting temperatures were obtained from the fluorescence melting profiles using Protein Thermal Shift software. Each variant was measured in three technical replicates from the same purified preparation, with wild type measured in parallel. For each variant, Δ*T*_*m*_ was calculated as the mean variant *T*_*m*_ minus the mean wild-type *T*_*m*_ from the same experimental batch. Reported values are mean ± s.d.

#### Activity assessment

Dehalogenase activity was measured using a continuous phenol-red pH-indicator assay with 1-iodobutane as substrate. Reactions were performed in 96-well plates at 30 °C in 200 µl containing 1 mM HEPES, pH 8.2, 20 µM phenol red, 0.03 mg ml^−1^ enzyme and 1 mM 1-iodobutane. The reaction was initiated by adding 1-iodobutane from a 100 mM stock prepared in methanol. Absorbance at 558 nm was recorded every 30 s for 30 min. Initial rates were calculated from the initial linear decrease in absorbance, typically over the first 2–5 min, after subtracting substrate-only blanks. Relative activity was defined as the blank-corrected initial rate of each variant divided by that of wild-type DhaA measured in the same experiment. Each variant was assayed in three technical replicates.

#### Statistical reporting

Summary statistics were computed from three technical replicates per variant for both *T*_*m*_ and relative activity. Replicate-level measurements are provided in dhaa_mutant_replicates.csv, and summary values are provided in dhaa_mutant_results.csv. Positive hit rate and stability-activity comparisons are reported descriptively.

## Data availability

The curated Test12K benchmark used for model evaluation, the Train226K training split derived from cDNA272K with leakage filtering, and the DhaA wet-lab measurements reported in this study are available in the project repository at https://github.com/ai4protein/Venus-MAXWELL. Source data underlying the main and supplementary figures are provided with the repository release. Public datasets reused in this study were obtained from the following sources: cDNA272K [30]; p53, myoglobin and Ssym [31]; S669 [32]; S8754 and M1261 [20]; vb1432 [33]; FireProtDB [34]; and ThermoMutDB [35]. Protein structures used for structure-conditioned inference were generated with ColabFold [61].

## Code availability

Custom code for training, evaluation, benchmark construction and figure reproduction is available at https://github.com/ai4protein/Venus-MAXWELL under an open-source licence. The repository includes scripts for reproducing the main bench-mark analyses, supplementary tables and the DhaA validation figures. Pretrained MAXWELL model weights are distributed with the same release.

## Acknowledgements

This work was supported by the National Key Research and Development Program of China (2024YFA0917603); Shanghai Municipal Science and Technology Major Project; the AI for Science Program, Shanghai Municipal Commission of Economy and Informatization (2025-GZL-RGZN-BTBX-02009); the Computational Biology Key Program of Shanghai Science and Technology Commission (23JS1400600); Shanghai Jiao Tong University Scientific and Technological Innovation Funds (21X010200843); Science and Technology Innovation Key R&D Program of Chongqing (CSTB2022TIAD-STX0017, CSTB2024TIAD-STX0032); the Student Innovation Center at Shanghai Jiao Tong University, and Shanghai Artificial Intelligence Laboratory.

## Author contributions

M.L. conceived the study, formulated the sequence-to-landscape one-step computation formula, developed the MAXWELL framework, performed computational analyses and wrote the manuscript. X.C. performed wet-lab experiments and analysed experimental data. M.H. performed computational analyses, prepared figures and contributed to manuscript revision. F.J. and Y.Y. contributed to dataset construction, participated in model implementation and evaluation, and performed biophysical analyses. L.H. and Y.Y. supervised the project, secured funding and revised the manuscript. All authors discussed the results and commented on the manuscript.

## Related publication

An earlier conference version, *Venus-MAXWELL: Efficient Learning of Protein-Mutation Stability Landscapes using Protein Language Models*, was posted in the NeurIPS 2025 main conference track and is available at https://papers.nips.cc/paper_files/paper/2025/hash/c88d0c9bea6230b518ce71268c8e49e0-Abstract-Conference.html. The present manuscript extends that conference paper by improving model performance, providing a more detailed analysis of the model outputs and adding wet-lab experimental validation.

## References

[1] Bell, E.L., Finnigan, W., France, S.P., Green, A.P., Hayes, M.A., Hepworth, L.J., Lovelock, S.L., Niikura, H., Osuna, S., Romero, E., et al.: Biocatalysis. Nature Reviews Methods Primers 1(1), 1–21 (2021)

[2] Sun, Z., Liu, Q., Qu, G., Feng, Y., Reetz, M.T.: Utility of b-factors in protein science: interpreting rigidity, flexibility, and internal motion and engineering thermostability. Chemical reviews 119(3), 1626–1665 (2019)

[3] Ebrahimi, S.B., Samanta, D.: Engineering protein-based therapeutics through structural and chemical design. Nature Communications 14(1), 2411 (2023)

[4] Jiang, F., Bian, J., Liu, H., Li, S., Bai, X., Zheng, L., Jin, S., Liu, Z., Yang, G.-Y., Hong, L.: Creatinase: Using increased entropy to improve the activity and thermostability. The Journal of Physical Chemistry B 127(12), 2671–2682 (2023)

[5] Jay, S.M., Lee, R.T.: Protein engineering for cardiovascular therapeutics: untapped potential for cardiac repair. Circulation research 113(7), 933–943 (2013)

[6] Gebauer, M., Skerra, A.: Engineered protein scaffolds as next-generation therapeutics. Annual review of pharmacology and toxicology 60(1), 391–415 (2020)

[7] Yang, K.K., Wu, Z., Arnold, F.H.: Machine-learning-guided directed evolution for protein engineering. Nature methods 16(8), 687–694 (2019)

[8] Hie, B.L., Yang, K.K.: Adaptive machine learning for protein engineering. Current opinion in structural biology 72, 145–152 (2022)

[9] Alford, R.F., Leaver-Fay, A., Jeliazkov, J.R., O’Meara, M.J., DiMaio, F.P., Park, H., Shapovalov, M.V., Renfrew, P.D., Mulligan, V.K., Kappel, K., et al.: The rosetta all-atom energy function for macromolecular modeling and design. Journal of chemical theory and computation 13(6), 3031–3048 (2017)

[10] Schymkowitz, J., Borg, J., Stricher, F., Nys, R., Rousseau, F., Serrano, L.: The foldx web server: an online force field. Nucleic acids research 33(suppl_2), 382–388 (2005)

[11] Yin, S., Ding, F., Dokholyan, N.V.: Eris: an automated estimator of protein stability. Nature methods 4(6), 466–467 (2007)

[12] Dehouck, Y., Kwasigroch, J.M., Gilis, D., Rooman, M.: Popmusic 2.1: a web server for the estimation of protein stability changes upon mutation and sequence optimality. BMC Bioinformatics 12, 151 (2011) 10.1186/1471-2105-12-151

[13] Pires, D.E., Ascher, D.B., Blundell, T.L.: mcsm: predicting the effects of mutations in proteins using graph-based signatures. Bioinformatics 30(3), 335–342 (2014)

[14] Sanavia, T., Birolo, G., Montanucci, L., Turina, P., Capriotti, E., Fariselli, P.: Limitations and challenges in protein stability prediction upon genome variations: towards future applications in precision medicine. Computational and Structural Biotechnology Journal 18, 1968–1979 (2020) 10.1016/j.csbj.2020.07.011

[15] Cao, H., Wang, J., He, L., Qi, Y., Zhang, J.Z.: Deepddg: predicting the stability change of protein point mutations using neural networks. Journal of chemical information and modeling 59(4), 1508–1514 (2019)

[16] Li, B., Yang, Y.T., Capra, J.A., Gerstein, M.B.: Predicting changes in protein thermodynamic stability upon point mutation with deep 3d convolutional neural networks. PLoS computational biology 16(11), 1008291 (2020)

[17] Wang, S., Tang, H., Shan, P., Wu, Z., Zuo, L.: Pros-gnn: predicting effects of mutations on protein stability using graph neural networks. Computational Biology and Chemistry 107, 107952 (2023)

[18] Zhou, Y., Pan, Q., Pires, D.E., Rodrigues, C.H., Ascher, D.B.: Ddmut: predicting effects of mutations on protein stability using deep learning. Nucleic Acids Research 51(W1), 122–128 (2023)

[19] Li, S.S., Liu, Z.M., Li, J., Ma, Y.B., Dong, Z.Y., Hou, J.W., Shen, F.J., Wang, W.B., Li, Q.M., Su, J.G.: Prediction of mutation-induced protein stability changes based on the geometric representations learned by a self-supervised method. BMC Bioinformatics 25, 269 (2024) 10.1186/s12859-024-05876-6

[20] Xu, Y., Liu, D., Gong, H.: Improving the prediction of protein stability changes upon mutations by geometric learning and a pre-training strategy. Nature Computational Science, 1–11 (2024)

[21] Meier, J., Rao, R., Verkuil, R., Liu, J., Sercu, T., Rives, A.: Language models enable zero-shot prediction of the effects of mutations on protein function. Advances in neural information processing systems 34, 29287–29303 (2021)

[22] Notin, P., Kollasch, A., Ritter, D., Van Niekerk, L., Paul, S., Spinner, H., Rollins, N., Shaw, A., Orenbuch, R., Weitzman, R., et al.: Proteingym: Large-scale bench-marks for protein fitness prediction and design. Advances in Neural Information Processing Systems 36, 64331–64379 (2023)

[23] Hsu, C., Verkuil, R., Liu, J., Lin, Z., Hie, B., Sercu, T., Lerer, A., Rives, A.: Learning inverse folding from millions of predicted structures. In: International Conference on Machine Learning, pp. 8946–8970 (2022). PMLR

[24] Tan, Y., Wang, R., Wu, B., Hong, L., Zhou, B.: From high-throughput evaluation to wet-lab studies: advancing mutation effect prediction with a retrieval-enhanced model. Bioinformatics 41(Supplement 1), 401–409 (2025) 10.1093/bioinformatics/btaf189

[25] Tsishyn, M., Hermans, P., Rooman, M., Pucci, F.: Residue conservation and sol-vent accessibility are (almost) all you need for predicting mutational effects in proteins. Bioinformatics 41(6), 322 (2025) 10.1093/bioinformatics/btaf322

[26] Tan, Y., Zhou, B., Zheng, L., Fan, G., Hong, L.: Semantical and geometrical protein encoding toward enhanced bioactivity and thermostability. eLife 13, 98033 (2025) 10.7554/eLife.98033

[27] Marquet, C., Schlensok, J., Abakarova, M., Rost, B., Laine, E.: Expert-guided protein language models enable accurate and blazingly fast fitness prediction. Bioinformatics 40(11), 621 (2024) 10.1093/bioinformatics/btae621

[28] Dieckhaus, H., Brocidiacono, M., Randolph, N.Z., Kuhlman, B.: Transfer learning to leverage larger datasets for improved prediction of protein stability changes. Proceedings of the National Academy of Sciences 121(6), 2314853121 (2024)

[29] Dauparas, J., Anishchenko, I., Bennett, N., Bai, H., Ragotte, R.J., Milles, L.F., Wicky, B.I., Courbet, A., Haas, R.J., Bethel, N., et al.: Robust deep learning– based protein sequence design using proteinmpnn. Science 378(6615), 49–56 (2022)

[30] Tsuboyama, K., Dauparas, J., Chen, J., Laine, E., Mohseni Behbahani, Y., Weinstein, J.J., Mangan, N.M., Ovchinnikov, S., Rocklin, G.J.: Mega-scale experimental analysis of protein folding stability in biology and design. Nature 620(7973), 434–444 (2023)

[31] Li, B., Yang, Y.T., Capra, J.A., Gerstein, M.B.: Predicting changes in protein thermodynamic stability upon point mutation with deep 3d convolutional neural networks. PLoS computational biology 16(11), 1008291 (2020)

[32] Pancotti, C., Benevenuta, S., Birolo, G., Alberini, V., Repetto, V., Sanavia, T., Capriotti, E., Fariselli, P.: Predicting protein stability changes upon single-point mutation: a thorough comparison of the available tools on a new dataset. Briefings in Bioinformatics 23(2), 555 (2022)

[33] Yang, Y., Urolagin, S., Niroula, A., Ding, X., Shen, B., Vihinen, M.: Pontstab: protein variant stability predictor. importance of training data quality. International journal of molecular sciences 19(4), 1009 (2018)

[34] Stourac, J., Dubrava, J., Musil, M., Horackova, J., Damborsky, J., Mazurenko, S., Bednar, D.: Fireprotdb: database of manually curated protein stability data. Nucleic acids research 49(D1), 319–324 (2021)

[35] Xavier, J.S., Nguyen, T.-B., Karmarkar, M., Portelli, S., Rezende, P.M., Velloso, J.P., Ascher, D.B., Pires, D.E.: Thermomutdb: a thermodynamic database for missense mutations. Nucleic acids research 49(D1), 475–479 (2021)

[36] Kulakova, A.N., Larkin, M.J., Kulakov, L.A.: The plasmid-located haloalkane dehalogenase gene from rhodococcus rhodochrous ncimb 13064. Microbiology 143(1), 109–115 (1997) 10.1099/00221287-143-1-109

[37] Visani, G.M., Pun, M.N., Galvin, W., Daniel, E., Borisiak, K., Wagura, U., Nourmohammad, A.: Hermes: Holographic equivariant neural network model for mutational effect and stability prediction. arXiv preprint arXiv:2407.06703 (2024)

[38] Ouyang-Zhang, J., Diaz, D.J., Klivans, A.R., Krähenbühl, P.: Predicting a protein’s stability under a million mutations. In: Advances in Neural Information Processing Systems, vol. 36 (2023)

[39] MacArthur, M.W., Thornton, J.M.: Influence of proline residues on protein conformation. Journal of Molecular Biology 218(2), 397–412 (1991) 10.1016/0022-2836(91)90721-H

[40] Li, S.C., Goto, N.K., Williams, K.A., Deber, C.M.: Alpha-helical, but not beta-sheet, propensity of proline is determined by peptide environment. Proceedings of the National Academy of Sciences of the United States of America 93(13), 6676–6681 (1996) 10.1073/pnas.93.13.6676

[41] Pace, C.N., Scholtz, J.M.: A helix propensity scale based on experimental studies of peptides and proteins. Biophysical Journal 75(1), 422–427 (1998) 10.1016/S0006-3495(98)77529-0

[42] Dill, K.A.: Dominant forces in protein folding. Biochemistry 29(31), 7133–7155 (1990) 10.1021/bi00483a001

[43] Kellis, J.T. Jr., Nyberg, K., Sali, D., Fersht, A.R.: Contribution of hydrophobic interactions to protein stability. Nature 333(6175), 784–786 (1988) 10.1038/333784a0

[44] Serrano, L., Bycroft, M., Fersht, A.R.: Aromatic-aromatic interactions and protein stability: investigation by double-mutant cycles. Journal of Molecular Biology 218(2), 465–475 (1991) 10.1016/0022-2836(91)90725-L

[45] Fersht, A.R., Serrano, L.: Principles of protein stability derived from protein engineering experiments. Current Opinion in Structural Biology 3(1), 75–83 (1993) 10.1016/0959-440X(93)90205-Y

[46] Hendsch, Z.S., Tidor, B.: Do salt bridges stabilize proteins? a continuum electrostatic analysis. Protein Science 3(2), 211–226 (1994) 10.1002/pro.5560030206

[47] Kumar, S., Nussinov, R.: Salt bridge stability in monomeric proteins. Journal of Molecular Biology 293(5), 1241–1255 (1999) 10.1006/jmbi.1999.3218

[48] Chothia, C.: The nature of the accessible and buried surfaces in proteins. Journal of Molecular Biology 105(1), 1–12 (1976) 10.1016/0022-2836(76)90191-1

[49] Miller, S., Janin, J., Lesk, A.M., Chothia, C.: Interior and surface of monomeric proteins. Journal of Molecular Biology 196(3), 641–656 (1987) 10.1016/0022-2836(87)90038-6

[50] Lin, Z., Akin, H., Rao, R., Hie, B., Zhu, Z., Lu, W., Smetanin, N., Verkuil, R., Kabeli, O., Shmueli, Y., et al.: Evolutionary-scale prediction of atomic-level protein structure with a language model. Science 379(6637), 1123–1130 (2023)

[51] Nijkamp, E., Ruffolo, J.A., Weinstein, E.N., Naik, N., Madani, A.: Progen2: exploring the boundaries of protein language models. Cell systems 14(11), 968–978 (2023)

[52] Yang, K.K., Zanichelli, N., Yeh, H.: Masked inverse folding with sequence transfer for protein representation learning. Protein Engineering, Design and Selection 36, 015 (2023) 10.1093/protein/gzad015

[53] Los, G.V., Encell, L.P., McDougall, M.G., Hartzell, D.D., Karassina, N., Zimprich, C., Wood, M.G., Learish, R., Ohana, R.F., Urh, M., Simpson, D., Mendez, J., Zimmerman, K., Otto, P., Vidugiris, G., Zhu, J., Darzins, A., Klaubert, D.H., Bulleit, R.F., Wood, K.V.: Halotag: A novel protein labeling technology for cell imaging and protein analysis. ACS Chemical Biology 3(6), 373–382 (2008) 10.1021/cb800025k

[54] Luo, Y., Jiang, G., Yu, T., Liu, Y., Vo, L., Ding, H., Su, Y., Qian, W.W., Zhao, H., Peng, J.: Ecnet is an evolutionary context-integrated deep learning framework for protein engineering. Nature communications 12(1), 5743 (2021)

[55] Notin, P., Weitzman, R., Marks, D., Gal, Y.: Proteinnpt: Improving protein property prediction and design with non-parametric transformers. Advances in Neural Information Processing Systems 36, 33529–33563 (2023)

[56] Jiang, F., Li, M., Dong, J., Yu, Y., Sun, X., Wu, B., Huang, J., Kang, L., Pei, Y., Zhang, L., et al.: A general temperature-guided language model to design proteins of enhanced stability and activity. Science Advances 10(48), 2641 (2024)

[57] Frellsen, J., Kassem, M.M., Bengtsen, T., Olsen, L., Lindorff-Larsen, K., Ferkinghoff-Borg, J., Boomsma, W.: Zero-shot protein stability prediction by inverse folding models: a free energy interpretation. arXiv preprint arXiv:2506.05596 (2025)

[58] Diaz, D.J., Gong, C., Ouyang-Zhang, J., Loy, J.M., Wells, J., Yang, D., Ellington, A.D., Dimakis, A.G., Klivans, A.R.: Stability oracle: a structure-based graph-transformer framework for identifying stabilizing mutations. Nature Communications 15(1), 6170 (2024)

[59] Steinegger, M., Söding, J.: Mmseqs2 enables sensitive protein sequence searching for the analysis of massive data sets. Nature biotechnology 35(11), 1026–1028 (2017)

[60] Rost, B.: Twilight zone of protein sequence alignments. Protein engineering 12(2), 85–94 (1999)

[61] Mirdita, M., Schütze, K., Moriwaki, Y., Heo, L., Ovchinnikov, S., Steinegger, M.: Colabfold: making protein folding accessible to all. Nature methods 19(6), 679–682 (2022)

[62] Likic, V.: The needleman-wunsch algorithm for sequence alignment. Lecture given at the 7th Melbourne Bioinformatics Course, Bi021 Molecular Science and Biotechnology Institute, University of Melbourne, 1–46 (2008)

[63] Kingma, D.P.: Adam: A method for stochastic optimization. arXiv preprint arXiv:1412.6980 (2014)

[64] Shrake, A., Rupley, J.A.: Environment and exposure to solvent of protein atoms. lysozyme and insulin. Journal of Molecular Biology 79(2), 351–371 (1973) 10.1016/0022-2836(73)90011-9

[65] Tien, M.Z., Meyer, A.G., Sydykova, D.K., Spielman, S.J., Wilke, C.O.: Maximum allowed solvent accessibilites of residues in proteins. PloS one 8(11), 80635 (2013)

