## Supplementary Information for "MAXWELL: Calibrating the probabilistic outputs of protein language models to the mutation-induced stability change landscape"

#### 1 Supplementary Methods

##### 1.1 Sequence-to-landscape learning algorithm

MAXWELL represents a wild-type protein as a sparse single-mutation landscape rather than as a collection of independent mutant sequences. For each protein $i$, the training tuple is $(\mathbf{A}_i, \mathbf{Y}_i, \mathbf{M}_i)$, where $\mathbf{A}_i$ is the wild-type sequence, $\mathbf{Y}_i$ is the measured stability landscape and $\mathbf{M}_i$ is the observed-entry mask. The pretrained protein model $f_\theta$ produces an amino-acid probability matrix and MAXWELL converts this matrix into a wild-type-normalized log-probability-ratio landscape by subtracting the wild-type residue log-probability from all amino-acid identities at the same position. The prediction and training procedure is:

1. Tokenize the wild-type protein sequence into a one-hot sequence matrix $\mathbf{X}_i$.
2. Compute the log-probability matrix $\mathbf{Q}_i = \log f_\theta(\mathbf{A}_i)$, or $\mathbf{Q}_i = \log f_\theta(\mathbf{A}_i, \mathbf{S}_i)$ when a structure-conditioned model is used.

3. Extract the wild-type log-probability vector $\mathbf{v}_i = (\mathbf{Q}_i \circ \mathbf{X}_i)\mathbf{1}_{|\mathcal{A}|}$.
4. Construct the predicted landscape $\mathbf{L}_i = \mathbf{Q}_i - \mathbf{v}_i \mathbf{1}_{|\mathcal{A}|}^{\top}$.
5. Select observed entries using $\mathbf{M}_i$.
6. Optimize the Pearson correlation objective over the observed entries.

This procedure keeps the full unmeasured landscape available for inference while preventing unmeasured entries from contributing artificial supervision. For prediction, a single forward pass through the pretrained model produces the complete position-by-amino-acid landscape, after which candidate mutations are ranked by the corresponding landscape entries.

#### 1.2 Pearson-only masked objective

The training objective was designed to optimize agreement between predicted and measured mutation preferences within each observed protein landscape. For observed entries, the current MAXWELL implementation uses one minus Pearson correlation between predicted and measured stability values. No MSE term, loss-weighting coefficient or separate calibration head is used. Thus, the predicted log-probability-ratio landscape itself is the optimized score surface, and sparse measured entries provide the only supervised signal. We also evaluated a listwise ranking objective for the ProteinMPNN-based landscape model under the same anchor setting of batch size 16 and learning rate $1 \times 10^{-4}$. The listwise objective did not improve validation Spearman or benchmark metrics relative to the Pearson-only objective. The Pearson-only model reached a validation Spearman correlation of 0.5298, mean per-protein Spearman 0.509 and AUPRC 0.629 in this ablation sweep, whereas the listwise objective reached a validation Spearman correlation of 0.5196, mean per-protein Spearman 0.499 and AUPRC 0.621. These results support retaining the simpler Pearson-only masked objective for the main ProteinMPNN-based model.

**Table S1 ProteinMPNN objective ablation.** Metrics follow the Figure 2 and Table 1 evaluation convention. Stabilizing mutations were defined as $-\Delta\Delta G \geq 0.3$. Correlation, classification and top-ranked recovery metrics were computed per protein and then averaged across proteins.

| Objective | Val. Spearman | Spearman | Pearson | AUROC | AUPRC | F1 | NDCG@10 | Recall@10 | Precision@10 |
|---|---|---|---|---|---|---|---|---|---|
| Pearson-only | 0.5298 | 0.509 | 0.521 | 0.771 | 0.629 | 0.675 | 0.698 | 0.768 | 0.319 |
| Listwise ranking | 0.5196 | 0.499 | 0.511 | 0.762 | 0.621 | 0.670 | 0.693 | 0.767 | 0.318 |

#### 1.3 Cross-validation and hyperparameter selection

Hyperparameters were selected without test-set leakage by evaluating validation performance on training-set proteins only. For each candidate hyperparameter setting, the training proteins were partitioned into five folds. Each fold was used once as the validation fold while the remaining four folds were used for internal training. The selected setting maximized the mean validation Spearman correlation across validation proteins. After selection, a final model was trained on the full training dataset for the selected number of epochs and evaluated once on the held-out Test12K benchmark. For the current ProteinMPNN Pearson-only search, we evaluated batch sizes $\{1, 2, 4, 8, 16, 32\}$ and learning rates $\{1 \times 10^{-5}, 5 \times 10^{-5}, 1 \times 10^{-4}, 5 \times 10^{-4}, 1 \times 10^{-3}\}$. The highest validation Spearman in this grid was observed at batch size 16 and learning rate $5 \times 10^{-4}$, with Spearman 0.547 (Supplementary Table S2).

**Table S2 ProteinMPNN-based MAXWELL batch-size and learning-rate grid search.** Spearman is the mean validation Spearman correlation averaged across five cross-validation folds on Train226K and was used for hyperparameter selection. The selected configuration is in **bold**.

| Batch size | Learning rate | Spearman |
|---|---|---|
| 1 | $1 \times 10^{-5}$ | 0.537 |
| 1 | $5 \times 10^{-5}$ | 0.542 |
| 1 | $1 \times 10^{-4}$ | 0.539 |
| 1 | $5 \times 10^{-4}$ | 0.524 |
| 1 | $1 \times 10^{-3}$ | 0.515 |
| 2 | $1 \times 10^{-5}$ | 0.535 |
| 2 | $5 \times 10^{-5}$ | 0.541 |
| 2 | $1 \times 10^{-4}$ | 0.543 |
| 2 | $5 \times 10^{-4}$ | 0.535 |
| 2 | $1 \times 10^{-3}$ | 0.523 |
| 4 | $1 \times 10^{-5}$ | 0.534 |
| 4 | $5 \times 10^{-5}$ | 0.540 |
| 4 | $1 \times 10^{-4}$ | 0.543 |
| 4 | $5 \times 10^{-4}$ | 0.544 |
| 4 | $1 \times 10^{-3}$ | 0.529 |
| 8 | $1 \times 10^{-5}$ | 0.524 |
| 8 | $5 \times 10^{-5}$ | 0.540 |
| 8 | $1 \times 10^{-4}$ | 0.545 |
| 8 | $5 \times 10^{-4}$ | 0.545 |
| 8 | $1 \times 10^{-3}$ | 0.539 |
| 16 | $1 \times 10^{-5}$ | 0.521 |
| 16 | $5 \times 10^{-5}$ | 0.535 |
| 16 | $1 \times 10^{-4}$ | 0.543 |
| **16** | $5 \times 10^{-4}$ | **0.547** |
| 16 | $1 \times 10^{-3}$ | 0.535 |
| 32 | $1 \times 10^{-5}$ | 0.502 |
| 32 | $5 \times 10^{-5}$ | 0.534 |
| 32 | $1 \times 10^{-4}$ | 0.540 |
| 32 | $5 \times 10^{-4}$ | 0.542 |
| 32 | $1 \times 10^{-3}$ | 0.538 |

#### 1.4 Evaluation metrics

Measured $\Delta\Delta G$ values were converted to an internal stability target $-\Delta\Delta G$, so larger values indicate greater stabilization. Stabilizing mutations were defined as

$-\Delta\Delta G \geq 0.3$ unless otherwise stated. Higher model scores were oriented to indicate stronger predicted stabilization before computing ranking and classification metrics. The primary regression metric was mean per-protein Spearman correlation. For each protein, Spearman correlation was computed between predicted and measured $-\Delta\Delta G$ values across the measured mutations available for that protein. The per-protein Spearman values were then averaged across proteins. This per-protein averaging prevents proteins with many measured mutations from dominating the benchmark. Pearson correlation was computed in the same per-protein manner and measures linear agreement between predicted scores and measured stability values. Kendall correlation was computed as Kendall's $\tau_b$ and measures pairwise rank concordance while accounting for ties. AUROC was computed for stabilizing-mutation classification and measures the probability that a randomly chosen stabilizing mutation receives a higher score than a randomly chosen non-stabilizing mutation. AUPRC was computed as average precision and emphasizes enrichment of stabilizing mutations under class imbalance. F1, Matthews correlation coefficient and balanced accuracy were computed by scanning score thresholds within each protein and retaining the best value for each metric. NDCG@10 evaluates whether stabilizing mutations are concentrated near the top of the ranked candidate list. Recall@10 is the fraction of all stabilizing mutations in a protein recovered among the top 10 ranked mutations. Precision@10 is the fraction of the top 10 ranked mutations that are stabilizing. Enrichment@10 is Precision@10 divided by the background stabilizing-mutation fraction for the same protein. Proteins without the positive or negative class required for a classification metric were excluded from the average for that metric.

##### 1.5 Baselines

The current main Table 1 benchmark comparison includes MAXWELL, ThermoMPNN, HERMES, Mutate Everything, FoldX, Rosetta, ProteinMPNN zero-shot scoring, ESM-IF1, VenusREM, RSALOR, ProtSSN and VespaG. ThermoMPNN is a structure-informed neural stability predictor derived from the ProteinMPNN modeling family and was included as a representative learned structure-based baseline [1]. HERMES is an equivariant neural network for mutational effect and stability prediction and was included as a representative geometric deep-learning baseline [2]. ProteinMPNN zero-shot scoring uses the native structure-conditioned inverse-folding log-probability-ratio signal from ProteinMPNN without supervised stability calibration [3]. FoldX is an empirical force-field method for estimating mutation-induced stability changes from protein structures [4]. Rosetta denotes a Rosetta all-atom Cartesian energy protocol implemented through PyRosetta Cartesian for mutation-effect scoring [5]. Mutate Everything is a large-scale neural model developed for predicting protein stability under many mutations and was included as an additional learned baseline [6]. ESM-IF1 is a structure-conditioned inverse-folding model for predicting amino-acid sequences from protein backbones and was included as a representative zero-shot inverse-folding baseline [7]. VenusREM is a retrieval-enhanced protein language model integrating sequence, structural and evolutionary information and was included as a representative retrieval-based zero-shot baseline [8]. RSALOR is an unsupervised mutation-effect predictor combining evolutionary log-odds with relative solvent accessibility and was included as a representative interpretable evolutionary–structural baseline [9]. ProtSSN is a sequence–structure model combining protein language model representations with equivariant graph neural networks and was included as a representative joint sequence–structure zero-shot baseline [10]. VespaG is an expert-guided mutation-effect predictor using protein language model embeddings and evolutionary pseudo-labels and was included as a representative sequence-based zero-shot baseline [11]. ProteinMPNN transfer, ABYSSAL, ESMTherm and other non-Cartesian Rosetta protocol values were not included in the main Table 1 comparison. These methods were excluded either because they were not part of the final matched main-baseline set or because the current manuscript emphasizes the concise comparison against representative physics-based, inverse-folding, geometric and supervised neural baselines.

#### 1.6 Computational environment

The current MAXWELL benchmark generation and model-evaluation runs were performed on an Ubuntu 22.04 command-line Linux instance with x86_64 architecture. The GPU was an NVIDIA GeForce RTX 4090 with 24,564 MiB memory. The NVIDIA driver version was 550.142 and the reported CUDA version was 12.4. The CPU was an AMD EPYC 7402 24-Core Processor exposed as 10 logical CPUs with one thread per core. The system had 52 GiB RAM and no configured swap. Benchmark generation used PyTorch 2.4.1 with TorchVision 0.19.1 and TorchAudio 2.4.1. The hardware and software environment is summarized in Supplementary Table S9.

#### 1.7 Dataset construction and leakage control

Mutation records were grouped by wild-type protein so that each model input corresponded to a sparse landscape. Training proteins with at least 30% sequence identity to test proteins were removed using MMseqs2 [12]. Needleman-Wunsch global alignment was used as an additional train-test separation check [13, 14]. The current filtered benchmark table contains 12,102 mutation records across 308 proteins, including 2,085 stabilizing mutations under the $-\Delta\Delta G \geq 0.3$ definition.

#### 1.8 Biophysical consistency analyses

The main biophysical analysis compares three matched mutation-preference views: ProteinMPNN zero-shot scores, experimental stability effects and calibrated MAXWELL scores. For Figure 3a, the analyzed set included 11,941 non-neutral single-point mutations from 308 proteins. The three plotted signals were ProteinMPNN zero-shot scores, experimental Stability defined as $-\Delta\Delta G$ and calibrated MAXWELL scores. Each signal was first z-scored within each protein, so proteins with larger raw score ranges did not dominate the global substitution pattern. Entries were then grouped by wild-type amino acid and mutant amino acid to form matched 20-by-20 substitution matrices. Each matrix cell reports the mean normalized Stability value for one wild-type-to-mutant substitution type. Entries with $\Delta\Delta G = 0$ were excluded when estimating experimental substitution patterns. Proline was placed first in both matrix axes because it has a distinctive role in protein stability and provides a sensitive visual check of the learned pattern. Physicochemical grouping included Hydrophobic, Aromatic, Polar,

Positive, Negative and Special amino-acid classes, where Special denotes glycine/proline. For Figure 3b, mutation records were grouped into the 36 wild-type-class to mutant-class transitions. For each transition, the calibrated MAXWELL Stability distribution and the experimental Stability distribution were compared. The transition-level mean values gave Pearson $r = 0.951$ and Spearman $\rho = 0.922$ between calibrated and experimental Stability. The sign of the class-level effect matched for 31 of 36 transitions. For Figure 3c, structural-context grouping used relative solvent accessibility classes, with Buried defined as RSA $\leq 0.20$, Intermediate as $0.20 <$ RSA $\leq 0.50$ and Exposed as RSA $> 0.50$. Within each solvent-exposure class, calibrated MAXWELL values were averaged by wild-type amino-acid class and mutant amino-acid class. This produced three matched 6-by-6 class-transition heatmaps for buried, intermediate and exposed residue environments. Cells with fewer than eight observed mutations were marked with a small black dot in the plotted panel. Complete substitution-level values are reported in Supplementary Tables S15, S16 and S17.

#### 1.9 Speed measurements

For the nine-method matched-length benchmark, all methods used the same physical NVIDIA RTX PRO 5000 72GB Blackwell GPU, eight allocated CPU cores, NVIDIA driver 595.84 and PyTorch 2.12.1+cu132 with CUDA runtime 13.2 (Supplementary Table S22). These software and allocation details describe panels a–c, rather than all experiments in the study. The three-method operational benchmarks in panels d and e were conducted separately on the same GPU model; they are not treated as additional repeats of panels a–c.

Prepared-input scoring began after input construction and device transfer but included learned sequence encoders required for scoring. Sequence embeddings and predictions were not reused across timed requests. HERMES included all ten supervised members, with site batches of at most 256. ProtSSN included all nine GNN ensemble members and recomputed the shared ESM2-650M embedding once per request. Mutate Everything used its full pretrained ESM2-650M encoder and single-substitution decoder; the double-substitution decoder was outside the requested workload. VespaG retained its FP16 ESM2-3B encoder and FP32 head; other methods used FP32. Gradients and TF32 were disabled. Prepared-input timings used ten warm-ups followed by at least 30 repetitions and five cumulative seconds of synchronized wall-clock measurement. CUDA-event timings were retained as diagnostics but were not used for the plotted throughput.

Input-preparation-plus-scoring measurements reconstructed model inputs from the supplied files for each of three repetitions, with model weights resident. This included PDB parsing, graph or structural-token construction, HERMES PyRosetta/Zernike preprocessing, MSA-frequency calculation, tokenization and device transfer when required by the model. Model loading, MSA search and structure prediction were excluded. Operating-system file caching was not disabled, so this scope is not a cold-machine run from FASTA alone. Prepared-input and preparation-plus- scoring timings were measured separately; subtracting their means does not provide an independently measured preprocessing time.

In panels a–c, ThermoMPNN computed ProteinMPNN features once per request and evaluated its native residue-wise prediction head $L$ times, reusing the amino-acid outputs at each position across the 19 substitutions. Scores were checked against its mutation-list interface on eight substitutions at every tested length, with absolute and relative tolerances of $5 \times 10^{-4}$. Panels d and e instead used the mutation-list scoring interface. No timing ratio was pooled across these implementations.

For panel d, each of three runs processed the same 308 proteins in a fixed order. The cumulative timer comprised one-time model loading and the sum of per-protein input preparation, scoring, score extraction and CSV-writing times. It did not include scheduler waiting, software startup before model loading or structure prediction. All nine model–run combinations completed 308 proteins and returned 1,222,574 non-wild-type scores. The main plot and inset use the same cumulative seconds; thin curves retain individual runs and thick curves summarize pointwise medians at each completed-protein index.

For panel e, 18 proteins were selected across length strata, comprising 15 Test12K proteins and three longer supplemental inputs. Lengths ranged from 64 to 936 residues. Each protein had eleven nested request sizes ending at $K = 19L$, and identical candidates were used for all methods. Model weights and input features were cached, but output predictions were recomputed. Timed requests included extraction and return of the requested scores to the CPU. Ten warm-ups preceded at least ten repeats and ten cumulative measured seconds per configuration. Points show per-protein, per-request-size medians. Curves use eleven logarithmically spaced request-fraction bins and plot the median request fraction against the median of the per-protein timing summaries in each bin. The first measured crossover is the smallest tested $K$ at which median MAXWELL latency does not exceed the comparator's; this is specific to the measured grid and does not establish an architecture-wide lower bound on advantageous request sizes.

#### 1.10 DhaA wet-lab validation

The DhaA wet-lab protocol is described in the main-text Methods, DhaA experimental validation. Wild-type and variant proteins were expressed from pET constructs with an N-terminal hexahistidine tag in BL21(DE3) cells, purified by nickel-nitrilotriacetic acid affinity chromatography, and verified by SDS–PAGE for representative variants (Supplementary Fig. S1). Thermal stability was measured by SYPRO Orange fluorescence thermal-shift differential scanning fluorimetry on a real-time PCR thermal cycler, and dehalogenase activity was measured with 1,2-dibromoethane and a pH-sensitive indicator at 620 nm. Each variant was measured in three technical replicates from the same purified preparation for both assays. Summary measurements and the model ranking of the ten tested variants are reported in Supplementary Tables S6 and S5.

### 2 Supplementary Results

#### 2.1 Current benchmark summary

The current filtered ProteinMPNN-landscape benchmark contains 12,102 mutations from 308 proteins. Among these records, 2,085 mutations are stabilizing under the $-\Delta\Delta G \geq 0.3$ definition. ProteinMPNN-landscape achieved a mean per-protein Spearman correlation of 0.54660. It also achieved a global Spearman correlation of 0.64806 in the current filtered benchmark table. For per-protein stabilizing-mutation discovery, ProteinMPNN-landscape achieved an AUPRC of 0.64928. The corresponding per-protein AUROC was 0.78892. Precision was 0.900 among the top 10 globally ranked mutations, 0.860 among the top 100 mutations and 0.678 at the top 5% global screening budget. The top 5% enrichment was 3.94-fold over the background stabilizing-mutation rate.

**Table S3 Current filtered ProteinMPNN-landscape benchmark summary.**

| Metric | Value |
|---|---|
| Mutation records | 12,102 |
| Proteins | 308 |
| Stabilizing mutations, $-\Delta\Delta G \geq 0.3$ | 2,085 |
| Mean per-protein Spearman | 0.54660 |
| Mean per-protein Pearson | 0.54745 |
| Global Spearman | 0.64806 |
| Mean per-protein AUPRC | 0.64928 |
| Mean per-protein AUROC | 0.78892 |
| Mean per-protein Recall@10 | 0.78195 |
| Mean per-protein Enrichment@10 | 1.93307 |
| Precision at top 10 globally | 0.900 |
| Precision at top 100 globally | 0.860 |
| Precision at top 5% globally | 0.678 |
| Enrichment at top 5% globally | 3.94 |

#### 2.2 ProteinMPNN-landscape calibration improves stabilizing-mutation discovery over ProteinMPNN zero-shot scoring

ProteinMPNN zero-shot scores provided a strong structure-conditioned amino-acid prior, but this prior was not directly calibrated for experimental thermostability. On the same filtered benchmark table, ProteinMPNN zero-shot scoring reached a mean per-protein AUPRC of 0.57473 and AUROC of 0.72795 for stabilizing-mutation discovery. After landscape calibration, ProteinMPNN-landscape improved AUPRC to 0.64928 and AUROC to 0.78892. Precision at the top 100 globally ranked candidates increased from 0.700 to 0.860. Precision at the top 5% global screening budget increased from 0.559 to 0.678, and enrichment increased from 3.25-fold to 3.94-fold.

**Table S4 ProteinMPNN zero-shot versus calibrated ProteinMPNN-landscape discovery metrics.** Stabilizing mutations are defined as $-\Delta\Delta G \geq 0.3$.

| Metric | ProteinMPNN zero-shot | ProteinMPNN-landscape |
|---|---|---|
| Mean per-protein AUPRC | 0.57473 | 0.64928 |
| Mean per-protein AUROC | 0.72795 | 0.78892 |
| Precision at top 10 globally | 0.800 | 0.900 |
| Precision at top 100 globally | 0.700 | 0.860 |
| Precision at top 5% globally | 0.559 | 0.678 |
| Enrichment at top 5% globally | 3.25 | 3.94 |

#### 2.3 Matched landscape scoring and operational efficiency

Prepared-input latency, effective throughput and input-preparation-plus-scoring latency for the nine GPU methods are reported in Supplementary Tables S18, S19 and S20. The $L = 957$ comparison is summarized in Supplementary Table S21. Throughput counts all $19L$ non-wild-type substitutions, excluding wild-type entries from methods that emit an $L \times 20$ matrix.

Dataset-level totals for the three-method operational benchmark are reported in Supplementary Table S23. All models completed all 308 proteins in all three runs. The median total for MAXWELL was 16.98 s, compared with 11.67 s for zero-shot ProteinMPNN and 363.34 s for ThermoMPNN. The shorter ProteinMPNN total reflected model-loading overhead: median processing times excluding loading were 9.03 s for MAXWELL and 11.45 s for zero-shot ProteinMPNN. The ThermoMPNN operational totals use its mutation-list interface and should not be interpreted as the total runtime of its residue-wise-output-reuse implementation.

In the candidate-request scan, the first measured MAXWELL crossover against the tested ThermoMPNN interface occurred at $K = 1$ for all 18 proteins. Against zero-shot ProteinMPNN, it occurred at $K = 1$ for 17 proteins and $K = 10$ for one 64-residue protein. These are descriptive grid-specific crossings, not evidence that one method is significantly faster at every request size. The two landscape-producing methods showed similar, approximately request-size-independent latency.

#### 2.4 DhaA validation measurements

The DhaA validation table contains 11 variants including wild type. G171W produced the largest measured $T_m$ increase, with $T_m = 59.83 \pm 0.20$°C and $\Delta T_m = 4.91$°C. E20Y and E20F increased $T_m$ by 1.95 °C and 1.74 °C, respectively. Unlike G171W, which retained 44.2% relative activity, E20Y and E20F retained 93.0% and 85.7% relative activity.

#### 2.5 Detailed benchmark tests and length-stratified baseline distributions

The concise main-text benchmark table reports only the mean ± s.d. summary values and significance markers for the main comparison. Supplementary Table S7 reports

**Table S5 MAXWELL ranking of the ten tested DhaA variants.** Variants were ranked by the calibrated landscape entry $L_{ij}$ defined in the main-text Methods without per-position deduplication. Measured $T_m$, $\Delta T_m$ and relative activity are shown for post hoc comparison.

| Rank | Variant | $L_{ij}$ | $\Delta T_m$ (°C) | Relative activity (%) | $T_m$ (°C) |
|---|---|---|---|---|---|
| 1 | K236I | 4.35 | 0.34 | 76.9 | $55.26 \pm 0.41$ |
| 2 | E20Y | 3.61 | 1.95 | 93.0 | $56.87 \pm 0.12$ |
| 3 | K236V | 3.43 | −0.66 | 74.0 | $54.26 \pm 0.15$ |
| 4 | K236M | 3.33 | 0.10 | 57.0 | $55.02 \pm 0.19$ |
| 5 | K236L | 3.23 | 0.15 | 74.0 | $55.07 \pm 0.21$ |
| 6 | T148W | 3.07 | −0.18 | 57.4 | $54.74 \pm 0.07$ |
| 7 | H84F | 3.02 | −3.26 | 78.0 | $51.66 \pm 0.17$ |
| 8 | K236F | 2.99 | 0.79 | 27.3 | $55.71 \pm 0.13$ |
| 9 | G171W | 2.97 | 4.91 | 44.2 | $59.83 \pm 0.20$ |
| 10 | E20F | 2.91 | 1.74 | 85.7 | $56.66 \pm 0.13$ |

**Table S6 DhaA thermal-stability and activity measurements.** Values are mean ± s.d.; $n = 3$ technical replicates for $T_m$ and relative activity.

| Variant | $T_m$ (°C) | $\Delta T_m$ (°C) | Relative activity (%) | $n$ |
|---|---|---|---|---|
| WT | $54.92 \pm 0.10$ | 0.00 | $100.0 \pm 4.7$ | 3 |
| G171W | $59.83 \pm 0.20$ | 4.91 | $44.2 \pm 3.6$ | 3 |
| E20Y | $56.87 \pm 0.12$ | 1.95 | $93.0 \pm 0.9$ | 3 |
| E20F | $56.66 \pm 0.13$ | 1.74 | $85.7 \pm 3.3$ | 3 |
| H84F | $51.66 \pm 0.17$ | -3.26 | $78.0 \pm 2.8$ | 3 |
| K236V | $54.26 \pm 0.15$ | -0.66 | $74.0 \pm 5.1$ | 3 |
| K236L | $55.07 \pm 0.21$ | 0.15 | $74.0 \pm 8.5$ | 3 |
| K236I | $55.26 \pm 0.41$ | 0.34 | $76.9 \pm 1.3$ | 3 |
| T148W | $54.74 \pm 0.07$ | -0.18 | $57.4 \pm 4.8$ | 3 |
| K236M | $55.02 \pm 0.19$ | 0.10 | $57.0 \pm 16.1$ | 3 |
| K236F | $55.71 \pm 0.13$ | 0.79 | $27.3 \pm 2.3$ | 3 |

the corresponding paired-test statistics, including raw Wilcoxon $P$ values and Holm-adjusted values. Supplementary Table S8 summarizes the same baseline set after stratifying proteins by sequence length.

#### 2.6 Detailed numerical values for Figure 2

Supplementary Tables S10, S11 and S12 report the numerical summaries underlying Figure 2a–c. Panels a and c use the same per-protein Spearman values from 308 test proteins, whereas panel b uses the 247 proteins with both stabilizing and non-stabilizing mutations for AUROC calculation. For panel c, the reported threshold fractions summarize the cumulative distribution rather than an additional hypothesis test.

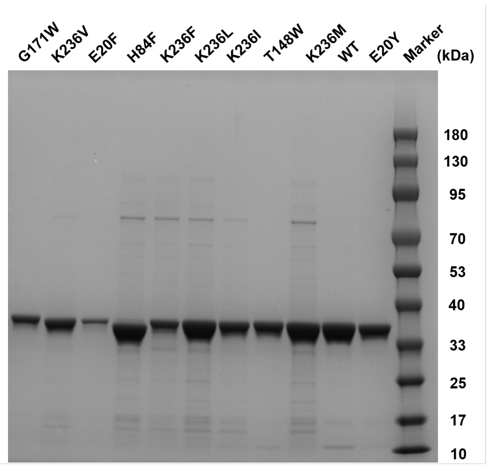


**Fig. S1 Supplementary Figure 1 — SDS–PAGE of representative purified DhaA variants.** Affinity-purified wild-type DhaA and selected variants were resolved by SDS–PAGE alongside a molecular-weight marker. Representative lanes include wild type, G171W, E20Y and E20F.

Supplementary Tables S13 and S14 report the paired tests corresponding to the significance markers in Figure 2a and 2b. Figure 2c is shown as a cumulative distribution only and does not include a separate hypothesis test.

**Table S7 Detailed paired tests underlying the main benchmark comparison.** Metric rows report two-sided paired Wilcoxon signed-rank tests comparing per-protein metrics between MAXWELL and each baseline. Holm-adjusted values are shown both within each metric and across all metric-baseline tests.

| Baseline | Metric or test | $n$ | MAXWELL mean | Baseline mean | Mean diff. | Median diff. | Win rate | $P$ / adjusted $P$ |
|---|---|---|---|---|---|---|---|---|
| ThermoMPNN | Spearman | 308 | 0.547 | 0.482 | 0.065 | 0.049 | 64.3% | $3.98 \times 10^{-10}$ / $1.19 \times 10^{-9}$ / $8.35 \times 10^{-9}$ |
| HERMES | Spearman | 308 | 0.547 | 0.469 | 0.077 | 0.050 | 63.6% | $2.07 \times 10^{-8}$ / $4.15 \times 10^{-8}$ / $3.53 \times 10^{-7}$ |
| ProteinMPNN | Spearman | 308 | 0.547 | 0.431 | 0.115 | 0.094 | 74.0% | $3.42 \times 10^{-19}$ / $1.37 \times 10^{-18}$ / $9.91 \times 10^{-18}$ |
| FoldX | Spearman | 308 | 0.547 | 0.389 | 0.157 | 0.135 | 77.3% | $1.18 \times 10^{-22}$ / $7.09 \times 10^{-22}$ / $3.90 \times 10^{-21}$ |
| Rosetta | Spearman | 308 | 0.547 | 0.364 | 0.182 | 0.167 | 78.2% | $8.69 \times 10^{-25}$ / $6.08 \times 10^{-24}$ / $2.96 \times 10^{-23}$ |
| Mutate Everything | Spearman | 308 | 0.547 | 0.371 | 0.176 | 0.153 | 74.0% | $3.97 \times 10^{-21}$ / $1.99 \times 10^{-20}$ / $1.27 \times 10^{-19}$ |
| ThermoMPNN | Pearson | 308 | 0.547 | 0.506 | 0.042 | 0.030 | 64.3% | $1.74 \times 10^{-5}$ / $3.48 \times 10^{-5}$ / $2.09 \times 10^{-4}$ |
| HERMES | Pearson | 308 | 0.547 | 0.490 | 0.058 | 0.039 | 63.3% | $3.01 \times 10^{-7}$ / $9.03 \times 10^{-7}$ / $4.82 \times 10^{-6}$ |
| ProteinMPNN | Pearson | 308 | 0.547 | 0.426 | 0.122 | 0.102 | 76.0% | $2.95 \times 10^{-20}$ / $1.48 \times 10^{-19}$ / $8.85 \times 10^{-19}$ |
| FoldX | Pearson | 308 | 0.547 | 0.415 | 0.132 | 0.131 | 73.1% | $2.39 \times 10^{-18}$ / $9.54 \times 10^{-18}$ / $6.68 \times 10^{-17}$ |
| Rosetta | Pearson | 308 | 0.547 | 0.319 | 0.229 | 0.219 | 82.1% | $1.92 \times 10^{-29}$ / $1.34 \times 10^{-28}$ / $6.72 \times 10^{-28}$ |
| Mutate Everything | Pearson | 308 | 0.547 | 0.383 | 0.164 | 0.131 | 73.1% | $1.84 \times 10^{-20}$ / $1.10 \times 10^{-19}$ / $5.71 \times 10^{-19}$ |
| ThermoMPNN | AUROC | 247 | 0.789 | 0.747 | 0.042 | 0.017 | 53.8% | $2.40 \times 10^{-6}$ / $7.19 \times 10^{-6}$ / $3.60 \times 10^{-5}$ |
| HERMES | AUROC | 247 | 0.789 | 0.764 | 0.025 | 0.006 | 51.0% | 0.003 / 0.006 / 0.023 |
| ProteinMPNN | AUROC | 247 | 0.789 | 0.728 | 0.061 | 0.040 | 61.5% | $7.53 \times 10^{-10}$ / $3.01 \times 10^{-9}$ / $1.51 \times 10^{-8}$ |
| FoldX | AUROC | 247 | 0.789 | 0.699 | 0.090 | 0.062 | 65.6% | $6.37 \times 10^{-12}$ / $3.82 \times 10^{-11}$ / $1.59 \times 10^{-10}$ |
| Rosetta | AUROC | 247 | 0.789 | 0.695 | 0.094 | 0.062 | 63.2% | $8.55 \times 10^{-11}$ / $4.28 \times 10^{-10}$ / $1.97 \times 10^{-9}$ |
| Mutate Everything | AUROC | 247 | 0.789 | 0.675 | 0.114 | 0.096 | 69.2% | $3.44 \times 10^{-15}$ / $2.41 \times 10^{-14}$ / $9.30 \times 10^{-14}$ |
| ThermoMPNN | AUPRC | 252 | 0.649 | 0.614 | 0.035 | 0.011 | 51.6% | $4.70 \times 10^{-5}$ / $1.41 \times 10^{-4}$ / $5.17 \times 10^{-4}$ |
| HERMES | AUPRC | 252 | 0.649 | 0.605 | 0.044 | 0.014 | 51.6% | $1.91 \times 10^{-4}$ / $3.82 \times 10^{-4}$ / 0.002 |
| ProteinMPNN | AUPRC | 252 | 0.649 | 0.575 | 0.075 | 0.039 | 60.7% | $1.77 \times 10^{-9}$ / $7.07 \times 10^{-9}$ / $3.36 \times 10^{-8}$ |
| FoldX | AUPRC | 252 | 0.649 | 0.544 | 0.105 | 0.050 | 62.7% | $3.02 \times 10^{-11}$ / $1.81 \times 10^{-10}$ / $7.26 \times 10^{-10}$ |
| Rosetta | AUPRC | 252 | 0.649 | 0.539 | 0.110 | 0.058 | 58.7% | $9.39 \times 10^{-11}$ / $4.69 \times 10^{-10}$ / $2.07 \times 10^{-9}$ |
| Mutate Everything | AUPRC | 252 | 0.649 | 0.542 | 0.107 | 0.105 | 66.3% | $6.43 \times 10^{-13}$ / $4.50 \times 10^{-12}$ / $1.67 \times 10^{-11}$ |
| ThermoMPNN | Enrich@10 | 252 | 1.933 | 1.871 | 0.063 | 0.000 | 17.1% | 0.087 / 0.262 / 0.573 |
| HERMES | Enrich@10 | 252 | 1.933 | 1.882 | 0.051 | 0.000 | 17.5% | 0.138 / 0.276 / 0.690 |
| ProteinMPNN | Enrich@10 | 252 | 1.933 | 1.795 | 0.138 | 0.000 | 20.2% | 0.002 / 0.007 / 0.016 |
| FoldX | Enrich@10 | 252 | 1.933 | 1.678 | 0.255 | 0.000 | 26.6% | $5.42 \times 10^{-6}$ / $2.71 \times 10^{-5}$ / $7.04 \times 10^{-5}$ |
| Rosetta | Enrich@10 | 252 | 1.933 | 1.600 | 0.333 | 0.000 | 27.0% | $3.28 \times 10^{-6}$ / $1.97 \times 10^{-5}$ / $4.59 \times 10^{-5}$ |
| Mutate Everything | Enrich@10 | 252 | 1.933 | 1.489 | 0.444 | 0.000 | 34.1% | $2.64 \times 10^{-9}$ / $1.85 \times 10^{-8}$ / $4.75 \times 10^{-8}$ |
| ESM-IF1 | Spearman | 308 | 0.522 | 0.350 | 0.172 | 0.158 | 76.6% | $6.68 \times 10^{-22}$ / $5.35 \times 10^{-21}$ / $3.21 \times 10^{-20}$ |
| VenusREM | Spearman | 308 | 0.522 | 0.347 | 0.174 | 0.164 | 71.8% | $8.00 \times 10^{-17}$ / $2.40 \times 10^{-16}$ / $2.96 \times 10^{-15}$ |
| RSALOR | Spearman | 307 | 0.523 | 0.312 | 0.211 | 0.183 | 75.9% | $5.65 \times 10^{-20}$ / $2.83 \times 10^{-19}$ / $2.43 \times 10^{-18}$ |
| ProtSSN | Spearman | 308 | 0.522 | 0.262 | 0.260 | 0.214 | 80.5% | $1.25 \times 10^{-25}$ / $1.37 \times 10^{-24}$ / $6.74 \times 10^{-24}$ |
| VespaG | Spearman | 308 | 0.522 | 0.272 | 0.250 | 0.218 | 77.3% | $1.08 \times 10^{-21}$ / $7.58 \times 10^{-21}$ / $5.09 \times 10^{-20}$ |
| ESM-IF1 | Pearson | 308 | 0.525 | 0.350 | 0.176 | 0.165 | 76.0% | $3.13 \times 10^{-23}$ / $2.82 \times 10^{-22}$ / $1.60 \times 10^{-21}$ |
| VenusREM | Pearson | 308 | 0.525 | 0.351 | 0.174 | 0.166 | 72.4% | $1.68 \times 10^{-18}$ / $6.71 \times 10^{-18}$ / $6.71 \times 10^{-17}$ |
| RSALOR | Pearson | 307 | 0.526 | 0.318 | 0.208 | 0.205 | 76.5% | $2.48 \times 10^{-19}$ / $1.24 \times 10^{-18}$ / $1.04 \times 10^{-17}$ |
| ProtSSN | Pearson | 308 | 0.525 | 0.272 | 0.253 | 0.205 | 78.2% | $1.33 \times 10^{-24}$ / $1.33 \times 10^{-23}$ / $6.93 \times 10^{-23}$ |
| VespaG | Pearson | 308 | 0.525 | 0.268 | 0.257 | 0.226 | 76.9% | $1.06 \times 10^{-22}$ / $8.46 \times 10^{-22}$ / $5.29 \times 10^{-21}$ |
| ESM-IF1 | AUROC | 247 | 0.770 | 0.669 | 0.100 | 0.094 | 64.8% | $1.96 \times 10^{-12}$ / $1.57 \times 10^{-11}$ / $6.09 \times 10^{-11}$ |
| VenusREM | AUROC | 247 | 0.770 | 0.666 | 0.104 | 0.083 | 65.6% | $8.04 \times 10^{-10}$ / $3.01 \times 10^{-9}$ / $1.69 \times 10^{-8}$ |
| RSALOR | AUROC | 247 | 0.770 | 0.641 | 0.128 | 0.110 | 67.6% | $5.81 \times 10^{-11}$ / $3.48 \times 10^{-10}$ / $1.51 \times 10^{-9}$ |
| ProtSSN | AUROC | 247 | 0.770 | 0.631 | 0.139 | 0.088 | 64.8% | $3.39 \times 10^{-13}$ / $3.05 \times 10^{-12}$ / $1.15 \times 10^{-11}$ |
| VespaG | AUROC | 247 | 0.770 | 0.593 | 0.177 | 0.158 | 72.9% | $6.08 \times 10^{-16}$ / $6.69 \times 10^{-15}$ / $2.19 \times 10^{-14}$ |
| ESM-IF1 | AUPRC | 247 | 0.620 | 0.495 | 0.125 | 0.074 | 64.4% | $8.89 \times 10^{-12}$ / $6.53 \times 10^{-11}$ / $2.49 \times 10^{-10}$ |
| VenusREM | AUPRC | 247 | 0.620 | 0.531 | 0.089 | 0.058 | 64.0% | $2.85 \times 10^{-7}$ / $8.56 \times 10^{-7}$ / $4.57 \times 10^{-6}$ |
| RSALOR | AUPRC | 247 | 0.620 | 0.483 | 0.137 | 0.085 | 66.4% | $1.55 \times 10^{-12}$ / $1.40 \times 10^{-11}$ / $4.97 \times 10^{-11}$ |
| ProtSSN | AUPRC | 247 | 0.620 | 0.506 | 0.114 | 0.088 | 66.8% | $8.16 \times 10^{-12}$ / $6.53 \times 10^{-11}$ / $2.37 \times 10^{-10}$ |
| VespaG | AUPRC | 247 | 0.620 | 0.437 | 0.183 | 0.167 | 73.7% | $9.12 \times 10^{-18}$ / $1.00 \times 10^{-16}$ / $3.47 \times 10^{-16}$ |
| ESM-IF1 | Enrich@10 | 195 | 2.080 | 1.779 | 0.301 | 0.000 | 37.9% | $1.56 \times 10^{-4}$ / $7.80 \times 10^{-4}$ / $1.09 \times 10^{-3}$ |
| VenusREM | Enrich@10 | 195 | 2.080 | 1.813 | 0.267 | 0.000 | 39.5% | $3.43 \times 10^{-4}$ / $1.37 \times 10^{-3}$ / $1.72 \times 10^{-3}$ |
| RSALOR | Enrich@10 | 195 | 2.080 | 1.654 | 0.426 | 0.000 | 45.6% | $5.19 \times 10^{-7}$ / $4.67 \times 10^{-6}$ / $7.26 \times 10^{-6}$ |
| ProtSSN | Enrich@10 | 195 | 2.080 | 1.687 | 0.393 | 0.000 | 45.6% | $2.33 \times 10^{-6}$ / $1.87 \times 10^{-5}$ / $3.03 \times 10^{-5}$ |
| VespaG | Enrich@10 | 195 | 2.080 | 1.466 | 0.614 | 0.000 | 50.8% | $2.98 \times 10^{-9}$ / $2.98 \times 10^{-8}$ / $5.37 \times 10^{-8}$ |

**Table S8 Length-stratified baseline coverage and performance.** Sequence lengths were obtained from the FASTA records used for the benchmark. Metrics are mean per-protein values within each sequence-length bin.

| Method | Length bin | Proteins | Mutations | Spearman | Pearson | AUROC | AUPRC | Enrich@10 |
|---|---|---|---|---|---|---|---|---|
| MAXWELL | $< 100$ | 81 | 2,895 | 0.702 | 0.691 | 0.873 | 0.766 | 1.921 |
| MAXWELL | 100–199 | 118 | 4,995 | 0.509 | 0.505 | 0.752 | 0.538 | 2.116 |
| MAXWELL | 200–399 | 72 | 2,400 | 0.443 | 0.443 | 0.737 | 0.654 | 1.487 |
| MAXWELL | $\geq 400$ | 37 | 1,812 | 0.527 | 0.574 | 0.795 | 0.741 | 2.165 |
| ThermoMPNN | $< 100$ | 81 | 2,895 | 0.656 | 0.668 | 0.819 | 0.699 | 1.817 |
| ThermoMPNN | 100–199 | 118 | 4,995 | 0.445 | 0.456 | 0.738 | 0.538 | 2.067 |
| ThermoMPNN | 200–399 | 72 | 2,400 | 0.342 | 0.388 | 0.660 | 0.617 | 1.557 |
| ThermoMPNN | $\geq 400$ | 37 | 1,812 | 0.487 | 0.539 | 0.746 | 0.662 | 1.901 |
| HERMES | $< 100$ | 81 | 2,895 | 0.663 | 0.667 | 0.839 | 0.688 | 1.928 |
| HERMES | 100–199 | 118 | 4,995 | 0.444 | 0.442 | 0.739 | 0.518 | 2.041 |
| HERMES | 200–399 | 72 | 2,400 | 0.335 | 0.374 | 0.703 | 0.625 | 1.453 |
| HERMES | $\geq 400$ | 37 | 1,812 | 0.385 | 0.479 | 0.765 | 0.666 | 2.017 |
| ProteinMPNN | $< 100$ | 81 | 2,895 | 0.536 | 0.509 | 0.778 | 0.634 | 1.735 |
| ProteinMPNN | 100–199 | 118 | 4,995 | 0.427 | 0.420 | 0.695 | 0.490 | 1.915 |
| ProteinMPNN | 200–399 | 72 | 2,400 | 0.329 | 0.348 | 0.699 | 0.613 | 1.569 |
| ProteinMPNN | $\geq 400$ | 37 | 1,812 | 0.414 | 0.417 | 0.767 | 0.652 | 1.952 |
| FoldX | $< 100$ | 81 | 2,895 | 0.505 | 0.536 | 0.744 | 0.622 | 1.706 |
| FoldX | 100–199 | 118 | 4,995 | 0.414 | 0.395 | 0.701 | 0.475 | 1.786 |
| FoldX | 200–399 | 72 | 2,400 | 0.239 | 0.331 | 0.619 | 0.565 | 1.476 |
| FoldX | $\geq 400$ | 37 | 1,812 | 0.352 | 0.376 | 0.720 | 0.549 | 1.600 |
| Rosetta | $< 100$ | 81 | 2,895 | 0.415 | 0.369 | 0.687 | 0.563 | 1.398 |
| Rosetta | 100–199 | 118 | 4,995 | 0.369 | 0.291 | 0.692 | 0.485 | 1.799 |
| Rosetta | 200–399 | 72 | 2,400 | 0.326 | 0.294 | 0.681 | 0.585 | 1.336 |
| Rosetta | $\geq 400$ | 37 | 1,812 | 0.308 | 0.348 | 0.755 | 0.585 | 1.922 |
| Mutate Everything | $< 100$ | 81 | 2,895 | 0.531 | 0.529 | 0.752 | 0.629 | 1.483 |
| Mutate Everything | 100–199 | 118 | 4,995 | 0.417 | 0.418 | 0.684 | 0.502 | 1.666 |
| Mutate Everything | 200–399 | 72 | 2,400 | 0.221 | 0.250 | 0.587 | 0.526 | 1.102 |
| Mutate Everything | $\geq 400$ | 37 | 1,812 | 0.169 | 0.214 | 0.592 | 0.493 | 1.613 |
| ESM-IF1 | $< 100$ | 81 | 2,895 | 0.480 | 0.443 | 0.715 | 0.561 | 1.768 |
| ESM-IF1 | 100–199 | 118 | 4,995 | 0.383 | 0.379 | 0.662 | 0.430 | 1.799 |
| ESM-IF1 | 200–399 | 72 | 2,400 | 0.187 | 0.224 | 0.589 | 0.478 | 1.503 |
| ESM-IF1 | $\geq 400$ | 37 | 1,812 | 0.274 | 0.297 | 0.725 | 0.595 | 2.248 |
| VenusREM | $< 100$ | 81 | 2,895 | 0.441 | 0.444 | 0.734 | 0.590 | 1.742 |
| VenusREM | 100–199 | 118 | 4,995 | 0.421 | 0.396 | 0.688 | 0.513 | 2.041 |
| VenusREM | 200–399 | 72 | 2,400 | 0.191 | 0.243 | 0.549 | 0.489 | 1.292 |
| VenusREM | $\geq 400$ | 37 | 1,812 | 0.209 | 0.216 | 0.612 | 0.509 | 1.985 |
| RSALOR | $< 100$ | 81 | 2,895 | 0.347 | 0.350 | 0.658 | 0.517 | 1.436 |
| RSALOR | 100–199 | 118 | 4,995 | 0.400 | 0.386 | 0.673 | 0.437 | 1.858 |
| RSALOR | 200–399 | 72 | 2,400 | 0.186 | 0.223 | 0.575 | 0.527 | 1.358 |
| RSALOR | $\geq 400$ | 37 | 1,812 | 0.200 | 0.215 | 0.600 | 0.480 | 1.935 |
| ProtSSN | $< 100$ | 81 | 2,895 | 0.303 | 0.280 | 0.664 | 0.536 | 1.440 |
| ProtSSN | 100–199 | 118 | 4,995 | 0.360 | 0.364 | 0.667 | 0.478 | 1.893 |
| ProtSSN | 200–399 | 72 | 2,400 | 0.121 | 0.179 | 0.524 | 0.492 | 1.411 |
| ProtSSN | $\geq 400$ | 37 | 1,812 | 0.130 | 0.141 | 0.612 | 0.555 | 2.004 |
| VespaG | $< 100$ | 81 | 2,895 | 0.305 | 0.287 | 0.587 | 0.425 | 1.226 |
| VespaG | 100–199 | 118 | 4,995 | 0.343 | 0.330 | 0.634 | 0.420 | 1.637 |
| VespaG | 200–399 | 72 | 2,400 | 0.139 | 0.164 | 0.499 | 0.448 | 1.103 |
| VespaG | $\geq 400$ | 37 | 1,812 | 0.232 | 0.236 | 0.637 | 0.514 | 2.079 |

**Table S9 Computational environment for current MAXWELL benchmark generation.**

| Component | Value |
|---|---|
| Operating system | Ubuntu 22.04 command-line Linux instance |
| Architecture | x86_64 |
| GPU | NVIDIA GeForce RTX 4090 |
| GPU memory | 24,564 MiB |
| NVIDIA driver | 550.142 |
| CUDA version | 12.4 |
| CPU | AMD EPYC 7402 24-Core Processor |
| Logical CPUs | 10 |
| Threads per core | 1 |
| System memory | 52 GiB |
| Swap | 0 B |
| PyTorch | 2.4.1 |
| TorchVision | 0.19.1 |
| TorchAudio | 2.4.1 |

**Table S10 Numerical summaries underlying Figure 2a.** Per-protein Spearman correlations are summarized across 308 test proteins for ProteinMPNN, ThermoMPNN and calibrated MAXWELL.

| Method | $n$ | Mean | s.d. | Median | IQR |
|---|---|---|---|---|---|
| ProteinMPNN | 308 | 0.431 | 0.320 | 0.489 | 0.269–0.666 |
| ThermoMPNN | 308 | 0.482 | 0.344 | 0.581 | 0.317–0.733 |
| MAXWELL | 308 | 0.547 | 0.312 | 0.633 | 0.371–0.762 |

**Table S11 Numerical summaries underlying Figure 2b.** Stabilizing-mutation AUROC values are summarized across 247 proteins with both positive and negative stabilizing-mutation labels under the $-\Delta\Delta G \geq 0.3$ threshold.

| Method | $n$ | Mean | s.d. | Median | IQR |
|---|---|---|---|---|---|
| ProteinMPNN | 247 | 0.728 | 0.213 | 0.781 | 0.595–0.880 |
| ThermoMPNN | 247 | 0.747 | 0.219 | 0.778 | 0.646–0.899 |
| MAXWELL | 247 | 0.789 | 0.200 | 0.833 | 0.678–0.941 |

**Table S12 Numerical summaries underlying Figure 2c.** Cumulative distribution of per-protein Spearman correlations across 308 test proteins. Threshold fractions report the fraction of proteins with Spearman greater than or equal to the indicated value.

| Method | $n$ | Mean | s.d. | Median | Spearman $\geq 0.4$ | Spearman $\geq 0.6$ |
|---|---|---|---|---|---|---|
| ProteinMPNN | 308 | 0.431 | 0.320 | 0.489 | 59.7% | 36.4% |
| ThermoMPNN | 308 | 0.482 | 0.344 | 0.581 | 68.2% | 47.4% |
| MAXWELL | 308 | 0.547 | 0.312 | 0.633 | 73.4% | 54.5% |

**Table S13 Paired tests underlying Figure 2a significance markers.** Values report two-sided paired Wilcoxon signed-rank tests comparing MAXWELL with the indicated comparator across 308 matched proteins. Asterisks follow the main-text convention: $^{*}P < 0.05$, $^{**}P < 0.01$ and $^{***}P < 0.001$.

| Metric | Comparison | $n$ | MAXWELL mean | Comparator mean | Raw $P$ value |
|---|---|---|---|---|---|
| Spearman | MAXWELL versus ProteinMPNN | 308 | 0.547 | 0.431 | $3.42 \times 10^{-19}$ *** |
| Spearman | MAXWELL versus ThermoMPNN | 308 | 0.547 | 0.482 | $3.98 \times 10^{-10}$ *** |

**Table S14 Paired tests underlying Figure 2b significance markers.** Values report two-sided paired Wilcoxon signed-rank tests comparing MAXWELL with the indicated comparator across 247 proteins with both stabilizing and non-stabilizing mutations. Asterisks follow the main-text convention: $^{*}P < 0.05$, $^{**}P < 0.01$ and $^{***}P < 0.001$.

| Metric | Comparison | $n$ | MAXWELL mean | Comparator mean | Raw $P$ value |
|---|---|---|---|---|---|
| AUROC | MAXWELL versus ProteinMPNN | 247 | 0.789 | 0.728 | $7.53 \times 10^{-10}$ *** |
| AUROC | MAXWELL versus ThermoMPNN | 247 | 0.789 | 0.747 | $2.40 \times 10^{-6}$ *** |

#### 2.7 Detailed numerical values for Figure 3a

Supplementary Tables S15, S16 and S17 report the complete substitution-level values underlying the three Figure 3a amino-acid substitution preference matrices. For each wild-type amino acid to mutant amino acid pair, the tables report the number of analyzed mutations and the mean within-protein z-score.

**Table S15**: **Numerical values for Figure 3a (ProteinMPNN zero-shot substitution-pattern matrix.)** Values report the mean within-protein z-score for each wild-type amino acid to mutant amino acid substitution type. The analysis includes non-neutral single-point mutations from the curated test set. Rows with $n = 0$ correspond to substitution cells without analyzed mutations after filtering and are shown as dashes.

| WT | Mutant | $n$ | Mean z-score |
|---|---|---|---|
| A | A | 0 | – |
| A | C | 16 | -0.382 |
| A | D | 22 | 0.171 |
| A | E | 32 | -0.136 |
| A | F | 35 | -0.540 |
| A | G | 260 | -0.093 |
| A | H | 22 | -0.525 |
| A | I | 38 | -0.555 |
| A | K | 29 | 0.378 |
| A | L | 57 | -0.440 |
| A | M | 25 | -0.764 |
| A | N | 19 | 0.082 |
| A | P | 40 | 0.012 |
| A | Q | 17 | 0.055 |
| A | R | 20 | -0.066 |
| A | S | 65 | 0.419 |
| A | T | 47 | -0.261 |
| A | V | 105 | -0.219 |
| A | W | 10 | -0.551 |
| A | Y | 25 | -0.453 |
| C | A | 57 | 0.435 |
| C | C | 0 | – |
| C | D | 0 | – |
| C | E | 3 | 1.589 |
| C | F | 1 | -1.671 |
| C | G | 6 | -1.171 |
| C | H | 0 | – |
| C | I | 4 | 0.475 |
| C | K | 0 | – |
| C | L | 2 | 0.569 |
| C | M | 2 | 0.077 |
| C | N | 1 | -0.981 |
| C | P | 1 | 1.950 |
| C | Q | 1 | 1.071 |
| C | R | 2 | -1.314 |
| C | S | 46 | 0.054 |
| C | T | 15 | 0.441 |
| C | V | 23 | 0.010 |
| C | W | 1 | -1.726 |
| C | Y | 3 | -0.193 |
| D | A | 197 | 0.118 |

Continued on next page

**Table S15: Continued.**

| WT | Mutant | $n$ | Mean z-score |
|---|---|---|---|
| D | C | 6 | -0.930 |
| D | D | 0 | – |
| D | E | 49 | 0.557 |
| D | F | 34 | -0.072 |
| D | G | 59 | 0.097 |
| D | H | 38 | -0.019 |
| D | I | 21 | -0.359 |
| D | K | 58 | 0.200 |
| D | L | 24 | 0.034 |
| D | M | 18 | -0.505 |
| D | N | 107 | 0.583 |
| D | P | 25 | -0.243 |
| D | Q | 22 | -0.040 |
| D | R | 28 | -0.122 |
| D | S | 30 | 0.737 |
| D | T | 18 | 0.006 |
| D | V | 19 | -0.441 |
| D | W | 13 | 0.187 |
| D | Y | 30 | 0.099 |
| E | A | 227 | 0.369 |
| E | C | 13 | -0.068 |
| E | D | 50 | 0.678 |
| E | E | 0 | – |
| E | F | 41 | 0.024 |
| E | G | 93 | -0.333 |
| E | H | 27 | -0.301 |
| E | I | 24 | 0.117 |
| E | K | 112 | 0.344 |
| E | L | 32 | 0.383 |
| E | M | 18 | 0.123 |
| E | N | 23 | 0.258 |
| E | P | 19 | -0.282 |
| E | Q | 118 | 0.427 |
| E | R | 21 | -0.124 |
| E | S | 28 | 0.176 |
| E | T | 31 | 0.517 |
| E | V | 39 | 0.238 |
| E | W | 18 | 0.042 |
| E | Y | 26 | -0.002 |
| F | A | 153 | -0.370 |
| F | C | 8 | -0.924 |
| F | D | 5 | -0.289 |
| F | E | 10 | -1.105 |
| F | F | 0 | – |
| F | G | 12 | -0.686 |
| F | H | 8 | -0.274 |
| F | I | 21 | -0.736 |
| F | K | 6 | -1.027 |
| F | L | 82 | -0.090 |
| F | M | 9 | -0.422 |
| F | N | 7 | -0.295 |
| F | P | 4 | -1.039 |
| F | Q | 4 | -0.889 |
| F | R | 3 | -1.161 |
| F | S | 25 | -0.719 |
| F | T | 7 | -0.605 |

Continued on next page

**Table S15**: **Continued.**

| WT | Mutant | $n$ | Mean z-score |
|---|---|---|---|
| F | V | 28 | -0.479 |
| F | W | 44 | -0.305 |
| F | Y | 34 | 0.831 |
| G | A | 219 | -0.517 |
| G | C | 15 | -0.877 |
| G | D | 42 | -0.866 |
| G | E | 19 | -0.678 |
| G | F | 19 | -0.953 |
| G | G | 0 | – |
| G | H | 15 | -0.877 |
| G | I | 8 | -0.904 |
| G | K | 15 | -0.874 |
| G | L | 16 | -0.908 |
| G | M | 8 | -0.903 |
| G | N | 19 | -0.381 |
| G | P | 10 | -1.081 |
| G | Q | 18 | -0.534 |
| G | R | 26 | -0.721 |
| G | S | 47 | -0.379 |
| G | T | 16 | -0.656 |
| G | V | 67 | -1.012 |
| G | W | 9 | -0.694 |
| G | Y | 16 | -0.913 |
| H | A | 76 | 0.563 |
| H | C | 1 | -0.084 |
| H | D | 6 | 0.755 |
| H | E | 15 | 1.128 |
| H | F | 9 | 1.070 |
| H | G | 32 | 0.384 |
| H | H | 0 | – |
| H | I | 1 | -1.092 |
| H | K | 5 | 0.890 |
| H | L | 7 | 1.015 |
| H | M | 0 | – |
| H | N | 15 | 0.529 |
| H | P | 3 | 0.631 |
| H | Q | 33 | 0.563 |
| H | R | 12 | 0.429 |
| H | S | 5 | 0.714 |
| H | T | 7 | 0.416 |
| H | V | 5 | -0.259 |
| H | W | 5 | -0.162 |
| H | Y | 13 | 0.823 |
| I | A | 282 | -0.582 |
| I | C | 7 | -0.519 |
| I | D | 12 | -0.662 |
| I | E | 14 | -0.293 |
| I | F | 24 | -0.772 |
| I | G | 42 | -0.672 |
| I | H | 2 | 0.325 |
| I | I | 0 | – |
| I | K | 7 | 0.467 |
| I | L | 59 | 0.290 |
| I | M | 47 | -0.345 |
| I | N | 4 | 0.760 |
| I | P | 3 | -0.419 |

Continued on next page

**Table S15**: **Continued.**

| WT | Mutant | $n$ | Mean z-score |
|---|---|---|---|
| I | Q | 5 | 0.079 |
| I | R | 5 | 0.080 |
| I | S | 15 | -0.432 |
| I | T | 41 | -0.168 |
| I | V | 238 | 0.698 |
| I | W | 7 | -0.384 |
| I | Y | 12 | -0.586 |
| K | A | 275 | 0.352 |
| K | C | 6 | -0.422 |
| K | D | 22 | 0.138 |
| K | E | 78 | 0.447 |
| K | F | 55 | -0.255 |
| K | G | 128 | -0.145 |
| K | H | 27 | -0.250 |
| K | I | 22 | -0.437 |
| K | K | 0 | – |
| K | L | 17 | -0.266 |
| K | M | 40 | -0.308 |
| K | N | 33 | 0.178 |
| K | P | 14 | -0.639 |
| K | Q | 81 | 0.132 |
| K | R | 41 | 0.477 |
| K | S | 20 | 0.447 |
| K | T | 23 | 0.522 |
| K | V | 24 | -0.060 |
| K | W | 4 | -0.903 |
| K | Y | 19 | -0.610 |
| L | A | 404 | -0.692 |
| L | C | 15 | -0.555 |
| L | D | 7 | -0.376 |
| L | E | 26 | -0.490 |
| L | F | 27 | -0.519 |
| L | G | 46 | -0.597 |
| L | H | 8 | 0.283 |
| L | I | 66 | -0.432 |
| L | K | 8 | 0.157 |
| L | L | 0 | – |
| L | M | 26 | -0.344 |
| L | N | 15 | -0.900 |
| L | P | 21 | -0.475 |
| L | Q | 11 | 0.048 |
| L | R | 19 | 0.237 |
| L | S | 13 | -0.346 |
| L | T | 25 | -0.195 |
| L | V | 112 | -0.383 |
| L | W | 10 | -0.513 |
| L | Y | 11 | 0.023 |
| M | A | 84 | 0.491 |
| M | C | 0 | – |
| M | D | 7 | 1.400 |
| M | E | 7 | 1.260 |
| M | F | 13 | 0.906 |
| M | G | 16 | 0.325 |
| M | H | 3 | 1.084 |
| M | I | 27 | 0.816 |
| M | K | 12 | 0.608 |

Continued on next page

**Table S15**: **Continued.**

| WT | Mutant | $n$ | Mean z-score |
|---|---|---|---|
| M | L | 34 | 1.389 |
| M | M | 0 | – |
| M | N | 3 | 1.445 |
| M | P | 5 | 2.185 |
| M | Q | 4 | 1.556 |
| M | R | 8 | 0.754 |
| M | S | 2 | 1.972 |
| M | T | 8 | 0.546 |
| M | V | 11 | 0.613 |
| M | W | 2 | -0.810 |
| M | Y | 5 | 1.432 |
| N | A | 132 | -0.075 |
| N | C | 6 | -0.703 |
| N | D | 57 | 0.179 |
| N | E | 15 | 0.262 |
| N | F | 12 | -0.091 |
| N | G | 54 | 0.220 |
| N | H | 16 | 0.337 |
| N | I | 15 | 0.168 |
| N | K | 24 | 0.623 |
| N | L | 13 | 0.075 |
| N | M | 15 | 0.239 |
| N | N | 0 | – |
| N | P | 8 | -0.578 |
| N | Q | 11 | 0.312 |
| N | R | 12 | 0.622 |
| N | S | 25 | -0.133 |
| N | T | 14 | -0.088 |
| N | V | 14 | 0.244 |
| N | W | 6 | -0.764 |
| N | Y | 13 | 0.217 |
| P | A | 149 | -0.345 |
| P | C | 0 | – |
| P | D | 0 | – |
| P | E | 0 | – |
| P | F | 6 | -0.912 |
| P | G | 56 | -0.532 |
| P | H | 0 | – |
| P | I | 1 | -1.181 |
| P | K | 0 | – |
| P | L | 11 | -0.701 |
| P | M | 1 | 0.055 |
| P | N | 2 | -0.052 |
| P | P | 0 | – |
| P | Q | 0 | – |
| P | R | 1 | -1.124 |
| P | S | 21 | -0.398 |
| P | T | 3 | -1.199 |
| P | V | 7 | -0.840 |
| P | W | 0 | – |
| P | Y | 0 | – |
| Q | A | 105 | 0.807 |
| Q | C | 3 | -0.955 |
| Q | D | 6 | 0.472 |
| Q | E | 23 | 0.867 |
| Q | F | 9 | 0.358 |

Continued on next page

**Table S15**: **Continued.**

| WT | Mutant | $n$ | Mean z-score |
|---|---|---|---|
| Q | G | 58 | 0.270 |
| Q | H | 9 | 0.422 |
| Q | I | 8 | 0.442 |
| Q | K | 12 | 1.047 |
| Q | L | 22 | 0.039 |
| Q | M | 5 | 0.233 |
| Q | N | 16 | -0.261 |
| Q | P | 11 | -0.165 |
| Q | Q | 0 | – |
| Q | R | 9 | 0.621 |
| Q | S | 10 | 0.181 |
| Q | T | 4 | 0.169 |
| Q | V | 8 | -0.114 |
| Q | W | 1 | -0.623 |
| Q | Y | 7 | -0.431 |
| R | A | 163 | 0.231 |
| R | C | 8 | -0.078 |
| R | D | 3 | -0.389 |
| R | E | 21 | 0.437 |
| R | F | 5 | -0.428 |
| R | G | 49 | 0.104 |
| R | H | 16 | 0.214 |
| R | I | 2 | -0.725 |
| R | K | 29 | 1.020 |
| R | L | 19 | 0.211 |
| R | M | 9 | 0.228 |
| R | N | 3 | -0.760 |
| R | P | 6 | 0.198 |
| R | Q | 27 | 0.628 |
| R | R | 0 | – |
| R | S | 9 | -0.285 |
| R | T | 4 | -0.052 |
| R | V | 5 | 0.193 |
| R | W | 8 | -0.445 |
| R | Y | 2 | -0.692 |
| S | A | 217 | 0.522 |
| S | C | 9 | -0.516 |
| S | D | 20 | 0.404 |
| S | E | 12 | 0.321 |
| S | F | 14 | 0.107 |
| S | G | 64 | 0.303 |
| S | H | 8 | -0.131 |
| S | I | 7 | 0.092 |
| S | K | 12 | 0.377 |
| S | L | 9 | 0.538 |
| S | M | 2 | 0.447 |
| S | N | 12 | 0.814 |
| S | P | 13 | 0.180 |
| S | Q | 5 | -0.064 |
| S | R | 13 | -0.014 |
| S | S | 0 | – |
| S | T | 22 | 0.523 |
| S | V | 17 | -0.283 |
| S | W | 7 | -0.652 |
| S | Y | 7 | -0.202 |
| T | A | 227 | -0.119 |

Continued on next page

**Table S15: Continued.**

| WT | Mutant | $n$ | Mean z-score |
|---|---|---|---|
| T | C | 26 | -0.349 |
| T | D | 44 | -0.322 |
| T | E | 43 | -0.092 |
| T | F | 28 | -0.640 |
| T | G | 84 | -0.538 |
| T | H | 37 | -0.640 |
| T | I | 60 | -0.030 |
| T | K | 26 | 0.620 |
| T | L | 32 | -0.462 |
| T | M | 29 | -0.444 |
| T | N | 38 | -0.031 |
| T | P | 29 | -0.765 |
| T | Q | 36 | -0.239 |
| T | R | 30 | -0.037 |
| T | S | 104 | 0.304 |
| T | T | 0 | – |
| T | V | 137 | 0.131 |
| T | W | 7 | -0.775 |
| T | Y | 34 | -0.737 |
| V | A | 400 | -0.380 |
| V | C | 23 | -0.793 |
| V | D | 17 | 0.621 |
| V | E | 26 | -0.157 |
| V | F | 32 | -0.373 |
| V | G | 86 | -0.387 |
| V | H | 19 | -0.542 |
| V | I | 121 | 0.403 |
| V | K | 14 | 0.539 |
| V | L | 84 | 0.094 |
| V | M | 39 | -0.056 |
| V | N | 26 | 0.038 |
| V | P | 11 | 0.700 |
| V | Q | 11 | 0.303 |
| V | R | 21 | 0.180 |
| V | S | 51 | -0.437 |
| V | T | 116 | -0.135 |
| V | V | 0 | – |
| V | W | 6 | -0.495 |
| V | Y | 27 | -0.071 |
| W | A | 44 | 0.060 |
| W | C | 3 | -0.542 |
| W | D | 0 | – |
| W | E | 9 | 0.087 |
| W | F | 76 | 0.051 |
| W | G | 0 | – |
| W | H | 16 | 0.438 |
| W | I | 0 | – |
| W | K | 0 | – |
| W | L | 15 | 0.209 |
| W | M | 4 | 1.747 |
| W | N | 3 | 2.081 |
| W | P | 0 | – |
| W | Q | 2 | -0.459 |
| W | R | 8 | 0.772 |
| W | S | 3 | 0.927 |
| W | T | 0 | – |

Continued on next page

**Table S15: Continued.**

| WT | Mutant | $n$ | Mean z-score |
|---|---|---|---|
| W | V | 2 | -0.515 |
| W | W | 0 | – |
| W | Y | 30 | 0.696 |
| Y | A | 134 | -0.360 |
| Y | C | 17 | -0.578 |
| Y | D | 20 | -0.148 |
| Y | E | 17 | -0.379 |
| Y | F | 188 | 0.453 |
| Y | G | 44 | -0.614 |
| Y | H | 16 | 0.163 |
| Y | I | 10 | -0.254 |
| Y | K | 10 | -0.512 |
| Y | L | 43 | -0.055 |
| Y | M | 8 | -0.700 |
| Y | N | 15 | -0.427 |
| Y | P | 8 | -1.014 |
| Y | Q | 13 | 0.102 |
| Y | R | 9 | -0.834 |
| Y | S | 22 | -0.230 |
| Y | T | 17 | -1.036 |
| Y | V | 14 | -0.237 |
| Y | W | 28 | 0.146 |
| Y | Y | 0 | – |

**Table S16: Numerical values for Figure 3a (Experimental $-\Delta\Delta G$ substitution-pattern matrix.)** Values report the mean within-protein z-score for each wild-type amino acid to mutant amino acid substitution type. The analysis includes non-neutral single-point mutations from the curated test set. Rows with $n = 0$ correspond to substitution cells without analyzed mutations after filtering and are shown as dashes.

| WT | Mutant | $n$ | Mean z-score |
|---|---|---|---|
| A | A | 0 | – |
| A | C | 16 | 0.419 |
| A | D | 22 | -0.002 |
| A | E | 32 | -0.652 |
| A | F | 35 | 0.029 |
| A | G | 260 | -0.156 |
| A | H | 22 | -0.363 |
| A | I | 38 | 0.145 |
| A | K | 29 | -0.160 |
| A | L | 57 | -0.044 |
| A | M | 25 | 0.101 |
| A | N | 19 | -0.126 |
| A | P | 40 | -0.300 |
| A | Q | 17 | 0.095 |
| A | R | 20 | -0.162 |
| A | S | 65 | 0.179 |
| A | T | 47 | -0.151 |
| A | V | 105 | 0.044 |
| A | W | 10 | -0.057 |
| A | Y | 25 | -0.283 |
| C | A | 57 | -0.117 |

Continued on next page

**Table S16**: **Continued.**

| WT | Mutant | $n$ | Mean z-score |
|---|---|---|---|
| C | C | 0 | – |
| C | D | 0 | – |
| C | E | 3 | 0.006 |
| C | F | 1 | 0.519 |
| C | G | 6 | -1.101 |
| C | H | 0 | – |
| C | I | 4 | 0.537 |
| C | K | 0 | – |
| C | L | 2 | 0.386 |
| C | M | 2 | 0.064 |
| C | N | 1 | 0.420 |
| C | P | 1 | -0.514 |
| C | Q | 1 | 1.171 |
| C | R | 2 | 0.123 |
| C | S | 46 | -0.380 |
| C | T | 15 | -0.265 |
| C | V | 23 | -0.585 |
| C | W | 1 | -0.410 |
| C | Y | 3 | -1.221 |
| D | A | 197 | 0.281 |
| D | C | 6 | 0.466 |
| D | D | 0 | – |
| D | E | 49 | 0.424 |
| D | F | 34 | 0.302 |
| D | G | 59 | 0.036 |
| D | H | 38 | 0.087 |
| D | I | 21 | 0.508 |
| D | K | 58 | 0.236 |
| D | L | 24 | 0.398 |
| D | M | 18 | 0.293 |
| D | N | 107 | 0.231 |
| D | P | 25 | -0.532 |
| D | Q | 22 | 0.226 |
| D | R | 28 | 0.325 |
| D | S | 30 | 0.496 |
| D | T | 18 | 0.234 |
| D | V | 19 | -0.087 |
| D | W | 13 | 0.527 |
| D | Y | 30 | 0.493 |
| E | A | 227 | 0.345 |
| E | C | 13 | 0.673 |
| E | D | 50 | 0.130 |
| E | E | 0 | – |
| E | F | 41 | 0.497 |
| E | G | 93 | -0.171 |
| E | H | 27 | -0.043 |
| E | I | 24 | 0.897 |
| E | K | 112 | 0.081 |
| E | L | 32 | 0.655 |
| E | M | 18 | 1.002 |
| E | N | 23 | 0.336 |
| E | P | 19 | -0.086 |
| E | Q | 118 | 0.309 |
| E | R | 21 | 0.127 |
| E | S | 28 | 0.063 |
| E | T | 31 | 0.355 |

Continued on next page

**Table S16**: **Continued.**

| WT | Mutant | $n$ | Mean z-score |
|---|---|---|---|
| E | V | 39 | 0.503 |
| E | W | 18 | 0.423 |
| E | Y | 26 | 0.530 |
| F | A | 153 | -0.823 |
| F | C | 8 | 0.059 |
| F | D | 5 | -0.677 |
| F | E | 10 | -1.543 |
| F | F | 0 | – |
| F | G | 12 | -1.719 |
| F | H | 8 | -1.174 |
| F | I | 21 | -0.231 |
| F | K | 6 | -0.984 |
| F | L | 82 | -0.426 |
| F | M | 9 | -0.473 |
| F | N | 7 | -0.827 |
| F | P | 4 | -1.484 |
| F | Q | 4 | -1.434 |
| F | R | 3 | -0.671 |
| F | S | 25 | -0.938 |
| F | T | 7 | -1.475 |
| F | V | 28 | -0.960 |
| F | W | 44 | 0.186 |
| F | Y | 34 | 0.587 |
| G | A | 219 | 0.058 |
| G | C | 15 | 0.463 |
| G | D | 42 | 0.010 |
| G | E | 19 | -0.289 |
| G | F | 19 | -0.131 |
| G | G | 0 | – |
| G | H | 15 | -0.173 |
| G | I | 8 | 0.274 |
| G | K | 15 | -0.129 |
| G | L | 16 | 0.070 |
| G | M | 8 | 0.360 |
| G | N | 19 | -0.197 |
| G | P | 10 | -0.679 |
| G | Q | 18 | 0.067 |
| G | R | 26 | -0.083 |
| G | S | 47 | 0.098 |
| G | T | 16 | -0.066 |
| G | V | 67 | -0.670 |
| G | W | 9 | -0.305 |
| G | Y | 16 | -0.187 |
| H | A | 76 | 0.147 |
| H | C | 1 | 1.093 |
| H | D | 6 | 0.245 |
| H | E | 15 | 0.520 |
| H | F | 9 | 0.687 |
| H | G | 32 | 0.098 |
| H | H | 0 | – |
| H | I | 1 | 0.909 |
| H | K | 5 | 0.332 |
| H | L | 7 | 1.074 |
| H | M | 0 | – |
| H | N | 15 | -0.134 |
| H | P | 3 | 0.395 |

Continued on next page

**Table S16**: **Continued.**

| WT | Mutant | $n$ | Mean z-score |
|---|---|---|---|
| H | Q | 33 | 0.295 |
| H | R | 12 | -0.013 |
| H | S | 5 | -0.094 |
| H | T | 7 | 0.030 |
| H | V | 5 | -0.029 |
| H | W | 5 | -0.151 |
| H | Y | 13 | 0.405 |
| I | A | 282 | -0.650 |
| I | C | 7 | 0.387 |
| I | D | 12 | -1.494 |
| I | E | 14 | -1.581 |
| I | F | 24 | -0.422 |
| I | G | 42 | -1.177 |
| I | H | 2 | -0.673 |
| I | I | 0 | – |
| I | K | 7 | -0.952 |
| I | L | 59 | 0.273 |
| I | M | 47 | 0.002 |
| I | N | 4 | 0.256 |
| I | P | 3 | -1.258 |
| I | Q | 5 | -1.260 |
| I | R | 5 | -1.057 |
| I | S | 15 | -1.240 |
| I | T | 41 | -0.625 |
| I | V | 238 | 0.296 |
| I | W | 7 | -1.270 |
| I | Y | 12 | -1.049 |
| K | A | 275 | 0.534 |
| K | C | 6 | 0.638 |
| K | D | 22 | -0.423 |
| K | E | 78 | 0.234 |
| K | F | 55 | 0.567 |
| K | G | 128 | 0.231 |
| K | H | 27 | 0.167 |
| K | I | 22 | 0.535 |
| K | K | 0 | – |
| K | L | 17 | 0.504 |
| K | M | 40 | 0.745 |
| K | N | 33 | 0.210 |
| K | P | 14 | -0.394 |
| K | Q | 81 | 0.205 |
| K | R | 41 | 0.260 |
| K | S | 20 | 0.323 |
| K | T | 23 | 0.088 |
| K | V | 24 | 0.509 |
| K | W | 4 | 0.054 |
| K | Y | 19 | 0.527 |
| L | A | 404 | -0.736 |
| L | C | 15 | -0.286 |
| L | D | 7 | -1.662 |
| L | E | 26 | -1.528 |
| L | F | 27 | 0.201 |
| L | G | 46 | -1.399 |
| L | H | 8 | 0.135 |
| L | I | 66 | 0.108 |
| L | K | 8 | -1.011 |

Continued on next page

**Table S16**: **Continued.**

| WT | Mutant | $n$ | Mean z-score |
|---|---|---|---|
| L | L | 0 | – |
| L | M | 26 | 0.277 |
| L | N | 15 | -1.084 |
| L | P | 21 | -1.469 |
| L | Q | 11 | -0.267 |
| L | R | 19 | 0.292 |
| L | S | 13 | -0.771 |
| L | T | 25 | -0.780 |
| L | V | 112 | -0.143 |
| L | W | 10 | 0.151 |
| L | Y | 11 | 0.098 |
| M | A | 84 | -0.227 |
| M | C | 0 | – |
| M | D | 7 | 0.177 |
| M | E | 7 | 1.099 |
| M | F | 13 | 0.742 |
| M | G | 16 | -0.510 |
| M | H | 3 | 0.819 |
| M | I | 27 | 0.317 |
| M | K | 12 | -0.457 |
| M | L | 34 | 0.476 |
| M | M | 0 | – |
| M | N | 3 | 0.593 |
| M | P | 5 | 0.839 |
| M | Q | 4 | 0.615 |
| M | R | 8 | 0.512 |
| M | S | 2 | 0.870 |
| M | T | 8 | 0.441 |
| M | V | 11 | 0.131 |
| M | W | 2 | 1.104 |
| M | Y | 5 | 0.876 |
| N | A | 132 | 0.099 |
| N | C | 6 | 0.315 |
| N | D | 57 | 0.097 |
| N | E | 15 | 0.017 |
| N | F | 12 | 0.640 |
| N | G | 54 | 0.236 |
| N | H | 16 | 0.425 |
| N | I | 15 | 0.681 |
| N | K | 24 | 0.246 |
| N | L | 13 | 0.562 |
| N | M | 15 | 0.894 |
| N | N | 0 | – |
| N | P | 8 | -1.357 |
| N | Q | 11 | 0.540 |
| N | R | 12 | 0.434 |
| N | S | 25 | 0.145 |
| N | T | 14 | 0.431 |
| N | V | 14 | 0.412 |
| N | W | 6 | -0.082 |
| N | Y | 13 | 1.025 |
| P | A | 149 | 0.100 |
| P | C | 0 | – |
| P | D | 0 | – |
| P | E | 0 | – |
| P | F | 6 | 0.807 |

Continued on next page

**Table S16: Continued.**

| WT | Mutant | $n$ | Mean z-score |
|---|---|---|---|
| P | G | 56 | 0.185 |
| P | H | 0 | – |
| P | I | 1 | -0.782 |
| P | K | 0 | – |
| P | L | 11 | 0.817 |
| P | M | 1 | 0.721 |
| P | N | 2 | -1.081 |
| P | P | 0 | – |
| P | Q | 0 | – |
| P | R | 1 | 1.893 |
| P | S | 21 | -0.100 |
| P | T | 3 | 0.712 |
| P | V | 7 | -0.019 |
| P | W | 0 | – |
| P | Y | 0 | – |
| Q | A | 105 | 0.506 |
| Q | C | 3 | 0.632 |
| Q | D | 6 | 0.182 |
| Q | E | 23 | 0.479 |
| Q | F | 9 | 0.478 |
| Q | G | 58 | 0.150 |
| Q | H | 9 | 0.441 |
| Q | I | 8 | 0.417 |
| Q | K | 12 | 0.601 |
| Q | L | 22 | 0.753 |
| Q | M | 5 | 0.616 |
| Q | N | 16 | -0.145 |
| Q | P | 11 | 0.094 |
| Q | Q | 0 | – |
| Q | R | 9 | 0.068 |
| Q | S | 10 | 0.286 |
| Q | T | 4 | 0.553 |
| Q | V | 8 | -0.566 |
| Q | W | 1 | 0.480 |
| Q | Y | 7 | 0.334 |
| R | A | 163 | -0.007 |
| R | C | 8 | -0.145 |
| R | D | 3 | -0.602 |
| R | E | 21 | 0.215 |
| R | F | 5 | -1.174 |
| R | G | 49 | -0.091 |
| R | H | 16 | -0.130 |
| R | I | 2 | -0.770 |
| R | K | 29 | -0.130 |
| R | L | 19 | 0.137 |
| R | M | 9 | -0.235 |
| R | N | 3 | -0.471 |
| R | P | 6 | -0.464 |
| R | Q | 27 | 0.152 |
| R | R | 0 | – |
| R | S | 9 | -0.722 |
| R | T | 4 | -0.639 |
| R | V | 5 | 0.393 |
| R | W | 8 | -0.056 |
| R | Y | 2 | -1.748 |
| S | A | 217 | 0.455 |

Continued on next page

**Table S16**: **Continued.**

| WT | Mutant | $n$ | Mean z-score |
|---|---|---|---|
| S | C | 9 | 0.538 |
| S | D | 20 | 0.118 |
| S | E | 12 | 0.452 |
| S | F | 14 | 0.488 |
| S | G | 64 | 0.426 |
| S | H | 8 | 0.329 |
| S | I | 7 | 1.139 |
| S | K | 12 | 0.555 |
| S | L | 9 | 0.800 |
| S | M | 2 | 0.956 |
| S | N | 12 | 0.283 |
| S | P | 13 | 0.113 |
| S | Q | 5 | -0.229 |
| S | R | 13 | 0.486 |
| S | S | 0 | – |
| S | T | 22 | 0.548 |
| S | V | 17 | 0.446 |
| S | W | 7 | 0.461 |
| S | Y | 7 | -0.257 |
| T | A | 227 | 0.071 |
| T | C | 26 | 0.539 |
| T | D | 44 | 0.008 |
| T | E | 43 | 0.190 |
| T | F | 28 | 0.537 |
| T | G | 84 | -0.167 |
| T | H | 37 | -0.067 |
| T | I | 60 | 0.654 |
| T | K | 26 | 0.359 |
| T | L | 32 | 0.355 |
| T | M | 29 | 0.344 |
| T | N | 38 | 0.274 |
| T | P | 29 | -0.933 |
| T | Q | 36 | -0.044 |
| T | R | 30 | 0.444 |
| T | S | 104 | 0.403 |
| T | T | 0 | – |
| T | V | 137 | 0.430 |
| T | W | 7 | 0.920 |
| T | Y | 34 | 0.477 |
| V | A | 400 | -0.262 |
| V | C | 23 | -0.038 |
| V | D | 17 | -0.012 |
| V | E | 26 | -1.281 |
| V | F | 32 | -0.112 |
| V | G | 86 | -0.821 |
| V | H | 19 | -0.655 |
| V | I | 121 | 0.372 |
| V | K | 14 | -0.768 |
| V | L | 84 | 0.110 |
| V | M | 39 | 0.246 |
| V | N | 26 | -0.576 |
| V | P | 11 | -0.229 |
| V | Q | 11 | -0.707 |
| V | R | 21 | -0.305 |
| V | S | 51 | -0.799 |
| V | T | 116 | -0.304 |

Continued on next page

**Table S16**: **Continued.**

| WT | Mutant | $n$ | Mean z-score |
|---|---|---|---|
| V | V | 0 | – |
| V | W | 6 | 0.162 |
| V | Y | 27 | -0.146 |
| W | A | 44 | -0.500 |
| W | C | 3 | -0.399 |
| W | D | 0 | – |
| W | E | 9 | -0.861 |
| W | F | 76 | -0.077 |
| W | G | 0 | – |
| W | H | 16 | -0.311 |
| W | I | 0 | – |
| W | K | 0 | – |
| W | L | 15 | 0.292 |
| W | M | 4 | -0.448 |
| W | N | 3 | -1.067 |
| W | P | 0 | – |
| W | Q | 2 | -1.772 |
| W | R | 8 | -0.862 |
| W | S | 3 | -1.269 |
| W | T | 0 | – |
| W | V | 2 | -0.631 |
| W | W | 0 | – |
| W | Y | 30 | -0.318 |
| Y | A | 134 | -0.662 |
| Y | C | 17 | 0.282 |
| Y | D | 20 | -0.741 |
| Y | E | 17 | -0.996 |
| Y | F | 188 | 0.046 |
| Y | G | 44 | -1.365 |
| Y | H | 16 | -0.485 |
| Y | I | 10 | -0.982 |
| Y | K | 10 | -0.456 |
| Y | L | 43 | -0.383 |
| Y | M | 8 | -0.915 |
| Y | N | 15 | -1.028 |
| Y | P | 8 | -1.479 |
| Y | Q | 13 | -0.565 |
| Y | R | 9 | -0.814 |
| Y | S | 22 | -0.548 |
| Y | T | 17 | -1.020 |
| Y | V | 14 | -0.808 |
| Y | W | 28 | 0.039 |
| Y | Y | 0 | – |

**Table S17**: **Numerical values for Figure 3a (Calibrated MAXWELL substitution-pattern matrix.)** Values report the mean within-protein z-score for each wild-type amino acid to mutant amino acid substitution type. The analysis includes non-neutral single-point mutations from the curated test set. Rows with $n = 0$ correspond to substitution cells without analyzed mutations after filtering and are shown as dashes.

| WT | Mutant | $n$ | Mean z-score |
|---|---|---|---|
| A | A | 0 | – |
| A | C | 16 | 0.786 |
| A | D | 22 | -0.348 |
| A | E | 32 | -0.675 |
| A | F | 35 | 0.261 |
| A | G | 260 | -0.273 |
| A | H | 22 | -0.057 |
| A | I | 38 | 0.257 |
| A | K | 29 | -0.120 |
| A | L | 57 | 0.199 |
| A | M | 25 | 0.112 |
| A | N | 19 | -0.173 |
| A | P | 40 | -0.727 |
| A | Q | 17 | -0.092 |
| A | R | 20 | 0.125 |
| A | S | 65 | 0.158 |
| A | T | 47 | -0.230 |
| A | V | 105 | 0.142 |
| A | W | 10 | 0.152 |
| A | Y | 25 | -0.113 |
| C | A | 57 | 0.046 |
| C | C | 0 | – |
| C | D | 0 | – |
| C | E | 3 | -0.026 |
| C | F | 1 | -1.185 |
| C | G | 6 | -1.341 |
| C | H | 0 | – |
| C | I | 4 | 0.769 |
| C | K | 0 | – |
| C | L | 2 | 0.626 |
| C | M | 2 | 0.036 |
| C | N | 1 | -1.142 |
| C | P | 1 | -0.157 |
| C | Q | 1 | -0.099 |
| C | R | 2 | -1.248 |
| C | S | 46 | -0.448 |
| C | T | 15 | -0.204 |
| C | V | 23 | 0.037 |
| C | W | 1 | -1.627 |
| C | Y | 3 | -0.226 |
| D | A | 197 | 0.363 |
| D | C | 6 | 1.108 |
| D | D | 0 | – |
| D | E | 49 | 0.349 |
| D | F | 34 | 0.541 |
| D | G | 59 | 0.023 |
| D | H | 38 | 0.448 |
| D | I | 21 | 0.657 |
| D | K | 58 | 0.048 |
| D | L | 24 | 0.602 |

Continued on next page

**Table S17**: **Continued.**

| WT | Mutant | $n$ | Mean z-score |
|---|---|---|---|
| D | M | 18 | 0.358 |
| D | N | 107 | 0.393 |
| D | P | 25 | -0.660 |
| D | Q | 22 | 0.107 |
| D | R | 28 | 0.097 |
| D | S | 30 | 0.585 |
| D | T | 18 | 0.254 |
| D | V | 19 | -0.046 |
| D | W | 13 | 1.044 |
| D | Y | 30 | 0.754 |
| E | A | 227 | 0.520 |
| E | C | 13 | 1.050 |
| E | D | 50 | 0.396 |
| E | E | 0 | – |
| E | F | 41 | 1.040 |
| E | G | 93 | -0.267 |
| E | H | 27 | 0.339 |
| E | I | 24 | 0.997 |
| E | K | 112 | 0.178 |
| E | L | 32 | 0.915 |
| E | M | 18 | 1.197 |
| E | N | 23 | 0.075 |
| E | P | 19 | -0.698 |
| E | Q | 118 | 0.505 |
| E | R | 21 | 0.454 |
| E | S | 28 | 0.224 |
| E | T | 31 | 0.467 |
| E | V | 39 | 0.745 |
| E | W | 18 | 1.160 |
| E | Y | 26 | 0.922 |
| F | A | 153 | -1.051 |
| F | C | 8 | -1.045 |
| F | D | 5 | -1.893 |
| F | E | 10 | -2.352 |
| F | F | 0 | – |
| F | G | 12 | -1.918 |
| F | H | 8 | -0.830 |
| F | I | 21 | -0.603 |
| F | K | 6 | -2.203 |
| F | L | 82 | -0.385 |
| F | M | 9 | -0.800 |
| F | N | 7 | -1.406 |
| F | P | 4 | -2.375 |
| F | Q | 4 | -2.064 |
| F | R | 3 | -2.690 |
| F | S | 25 | -1.424 |
| F | T | 7 | -1.302 |
| F | V | 28 | -0.939 |
| F | W | 44 | -0.084 |
| F | Y | 34 | 0.330 |
| G | A | 219 | -0.070 |
| G | C | 15 | 0.642 |
| G | D | 42 | -0.594 |
| G | E | 19 | -0.627 |
| G | F | 19 | -0.090 |
| G | G | 0 | – |

Continued on next page

**Table S17: Continued.**

| WT | Mutant | $n$ | Mean z-score |
|---|---|---|---|
| G | H | 15 | -0.119 |
| G | I | 8 | -0.427 |
| G | K | 15 | -0.760 |
| G | L | 16 | 0.075 |
| G | M | 8 | -0.024 |
| G | N | 19 | -0.204 |
| G | P | 10 | -1.123 |
| G | Q | 18 | -0.219 |
| G | R | 26 | -0.498 |
| G | S | 47 | -0.006 |
| G | T | 16 | 0.024 |
| G | V | 67 | -0.580 |
| G | W | 9 | 0.279 |
| G | Y | 16 | 0.092 |
| H | A | 76 | 0.234 |
| H | C | 1 | 0.766 |
| H | D | 6 | -0.056 |
| H | E | 15 | -0.050 |
| H | F | 9 | 1.288 |
| H | G | 32 | -0.106 |
| H | H | 0 | – |
| H | I | 1 | 0.989 |
| H | K | 5 | -0.064 |
| H | L | 7 | 0.985 |
| H | M | 0 | – |
| H | N | 15 | -0.122 |
| H | P | 3 | -0.130 |
| H | Q | 33 | 0.358 |
| H | R | 12 | 0.173 |
| H | S | 5 | 0.246 |
| H | T | 7 | 0.506 |
| H | V | 5 | 0.495 |
| H | W | 5 | 1.076 |
| H | Y | 13 | 1.354 |
| I | A | 282 | -0.953 |
| I | C | 7 | 0.028 |
| I | D | 12 | -2.211 |
| I | E | 14 | -1.568 |
| I | F | 24 | -0.673 |
| I | G | 42 | -1.522 |
| I | H | 2 | -0.343 |
| I | I | 0 | – |
| I | K | 7 | -0.764 |
| I | L | 59 | 0.331 |
| I | M | 47 | -0.046 |
| I | N | 4 | -0.841 |
| I | P | 3 | -1.817 |
| I | Q | 5 | -1.030 |
| I | R | 5 | -0.846 |
| I | S | 15 | -1.515 |
| I | T | 41 | -0.992 |
| I | V | 238 | 0.423 |
| I | W | 7 | -0.085 |
| I | Y | 12 | -0.856 |
| K | A | 275 | 0.592 |
| K | C | 6 | 1.206 |

Continued on next page

**Table S17**: **Continued.**

| WT | Mutant | $n$ | Mean z-score |
|---|---|---|---|
| K | D | 22 | -0.332 |
| K | E | 78 | 0.165 |
| K | F | 55 | 0.751 |
| K | G | 128 | 0.023 |
| K | H | 27 | 0.416 |
| K | I | 22 | 0.388 |
| K | K | 0 | – |
| K | L | 17 | 0.514 |
| K | M | 40 | 0.891 |
| K | N | 33 | -0.108 |
| K | P | 14 | -0.836 |
| K | Q | 81 | 0.268 |
| K | R | 41 | 0.744 |
| K | S | 20 | 0.258 |
| K | T | 23 | 0.310 |
| K | V | 24 | 0.530 |
| K | W | 4 | 0.441 |
| K | Y | 19 | 0.308 |
| L | A | 404 | -0.952 |
| L | C | 15 | 0.024 |
| L | D | 7 | -1.746 |
| L | E | 26 | -1.436 |
| L | F | 27 | 0.009 |
| L | G | 46 | -1.260 |
| L | H | 8 | -0.254 |
| L | I | 66 | 0.012 |
| L | K | 8 | -1.099 |
| L | L | 0 | – |
| L | M | 26 | 0.025 |
| L | N | 15 | -1.222 |
| L | P | 21 | -1.325 |
| L | Q | 11 | -0.707 |
| L | R | 19 | -0.586 |
| L | S | 13 | -1.171 |
| L | T | 25 | -1.247 |
| L | V | 112 | -0.278 |
| L | W | 10 | -0.323 |
| L | Y | 11 | -0.038 |
| M | A | 84 | -0.107 |
| M | C | 0 | – |
| M | D | 7 | 0.216 |
| M | E | 7 | 0.253 |
| M | F | 13 | 0.800 |
| M | G | 16 | -0.774 |
| M | H | 3 | 0.678 |
| M | I | 27 | 0.551 |
| M | K | 12 | -0.370 |
| M | L | 34 | 0.857 |
| M | M | 0 | – |
| M | N | 3 | 0.467 |
| M | P | 5 | 1.139 |
| M | Q | 4 | 0.870 |
| M | R | 8 | 0.329 |
| M | S | 2 | 0.765 |
| M | T | 8 | 0.137 |
| M | V | 11 | 0.166 |

Continued on next page

**Table S17: Continued.**

| WT | Mutant | $n$ | Mean z-score |
|---|---|---|---|
| M | W | 2 | 0.192 |
| M | Y | 5 | 0.852 |
| N | A | 132 | 0.218 |
| N | C | 6 | 1.023 |
| N | D | 57 | -0.195 |
| N | E | 15 | 0.245 |
| N | F | 12 | 0.894 |
| N | G | 54 | 0.258 |
| N | H | 16 | 0.884 |
| N | I | 15 | 1.110 |
| N | K | 24 | 0.071 |
| N | L | 13 | 0.907 |
| N | M | 15 | 1.407 |
| N | N | 0 | – |
| N | P | 8 | -1.091 |
| N | Q | 11 | 0.345 |
| N | R | 12 | 0.789 |
| N | S | 25 | -0.008 |
| N | T | 14 | 0.137 |
| N | V | 14 | 1.021 |
| N | W | 6 | 1.045 |
| N | Y | 13 | 1.132 |
| P | A | 149 | 0.202 |
| P | C | 0 | – |
| P | D | 0 | – |
| P | E | 0 | – |
| P | F | 6 | 0.632 |
| P | G | 56 | 0.072 |
| P | H | 0 | – |
| P | I | 1 | -0.117 |
| P | K | 0 | – |
| P | L | 11 | 0.521 |
| P | M | 1 | 0.934 |
| P | N | 2 | 0.412 |
| P | P | 0 | – |
| P | Q | 0 | – |
| P | R | 1 | -0.019 |
| P | S | 21 | 0.235 |
| P | T | 3 | -0.122 |
| P | V | 7 | -0.019 |
| P | W | 0 | – |
| P | Y | 0 | – |
| Q | A | 105 | 0.747 |
| Q | C | 3 | -0.536 |
| Q | D | 6 | -0.127 |
| Q | E | 23 | 0.288 |
| Q | F | 9 | 0.677 |
| Q | G | 58 | 0.004 |
| Q | H | 9 | 0.691 |
| Q | I | 8 | 1.271 |
| Q | K | 12 | 0.451 |
| Q | L | 22 | 1.213 |
| Q | M | 5 | 0.670 |
| Q | N | 16 | -0.756 |
| Q | P | 11 | -1.200 |
| Q | Q | 0 | – |

Continued on next page

**Table S17**: **Continued.**

| WT | Mutant | $n$ | Mean z-score |
|---|---|---|---|
| Q | R | 9 | 0.802 |
| Q | S | 10 | 0.177 |
| Q | T | 4 | 0.152 |
| Q | V | 8 | 0.840 |
| Q | W | 1 | 0.047 |
| Q | Y | 7 | 0.145 |
| R | A | 163 | 0.138 |
| R | C | 8 | 0.584 |
| R | D | 3 | -1.598 |
| R | E | 21 | 0.040 |
| R | F | 5 | -0.224 |
| R | G | 49 | -0.233 |
| R | H | 16 | 0.299 |
| R | I | 2 | -0.356 |
| R | K | 29 | 0.508 |
| R | L | 19 | 0.132 |
| R | M | 9 | 0.858 |
| R | N | 3 | -1.215 |
| R | P | 6 | -0.269 |
| R | Q | 27 | 0.480 |
| R | R | 0 | – |
| R | S | 9 | -0.356 |
| R | T | 4 | 0.007 |
| R | V | 5 | 0.304 |
| R | W | 8 | 0.112 |
| R | Y | 2 | -0.464 |
| S | A | 217 | 0.582 |
| S | C | 9 | 0.372 |
| S | D | 20 | -0.047 |
| S | E | 12 | 0.242 |
| S | F | 14 | 0.534 |
| S | G | 64 | 0.263 |
| S | H | 8 | 0.421 |
| S | I | 7 | 0.786 |
| S | K | 12 | 0.299 |
| S | L | 9 | 1.289 |
| S | M | 2 | 1.578 |
| S | N | 12 | 0.462 |
| S | P | 13 | -0.136 |
| S | Q | 5 | -0.543 |
| S | R | 13 | 0.147 |
| S | S | 0 | – |
| S | T | 22 | 0.681 |
| S | V | 17 | 0.244 |
| S | W | 7 | 0.524 |
| S | Y | 7 | 0.144 |
| T | A | 227 | 0.154 |
| T | C | 26 | 0.752 |
| T | D | 44 | -0.464 |
| T | E | 43 | -0.266 |
| T | F | 28 | 0.472 |
| T | G | 84 | -0.541 |
| T | H | 37 | 0.085 |
| T | I | 60 | 0.584 |
| T | K | 26 | 0.208 |
| T | L | 32 | 0.237 |

Continued on next page

**Table S17: Continued.**

| WT | Mutant | $n$ | Mean z-score |
|---|---|---|---|
| T | M | 29 | 0.354 |
| T | N | 38 | 0.054 |
| T | P | 29 | -1.386 |
| T | Q | 36 | -0.071 |
| T | R | 30 | 0.406 |
| T | S | 104 | 0.237 |
| T | T | 0 | – |
| T | V | 137 | 0.353 |
| T | W | 7 | 0.240 |
| T | Y | 34 | 0.342 |
| V | A | 400 | -0.396 |
| V | C | 23 | -0.014 |
| V | D | 17 | -0.144 |
| V | E | 26 | -1.031 |
| V | F | 32 | 0.207 |
| V | G | 86 | -0.840 |
| V | H | 19 | -0.753 |
| V | I | 121 | 0.695 |
| V | K | 14 | -0.640 |
| V | L | 84 | 0.468 |
| V | M | 39 | 0.392 |
| V | N | 26 | -0.500 |
| V | P | 11 | -0.398 |
| V | Q | 11 | -0.538 |
| V | R | 21 | -0.429 |
| V | S | 51 | -0.913 |
| V | T | 116 | -0.392 |
| V | V | 0 | – |
| V | W | 6 | -0.266 |
| V | Y | 27 | 0.350 |
| W | A | 44 | -0.674 |
| W | C | 3 | -0.514 |
| W | D | 0 | – |
| W | E | 9 | -0.674 |
| W | F | 76 | -0.201 |
| W | G | 0 | – |
| W | H | 16 | -0.102 |
| W | I | 0 | – |
| W | K | 0 | – |
| W | L | 15 | -0.227 |
| W | M | 4 | 0.850 |
| W | N | 3 | 0.120 |
| W | P | 0 | – |
| W | Q | 2 | -1.767 |
| W | R | 8 | -0.262 |
| W | S | 3 | -0.587 |
| W | T | 0 | – |
| W | V | 2 | -1.676 |
| W | W | 0 | – |
| W | Y | 30 | -0.047 |
| Y | A | 134 | -0.885 |
| Y | C | 17 | 0.319 |
| Y | D | 20 | -0.855 |
| Y | E | 17 | -1.126 |
| Y | F | 188 | 0.389 |
| Y | G | 44 | -1.558 |

Continued on next page

**Table S17**: **Continued.**

| WT | Mutant | $n$ | Mean z-score |
|---|---|---|---|
| Y | H | 16 | -0.227 |
| Y | I | 10 | -0.205 |
| Y | K | 10 | -1.634 |
| Y | L | 43 | -0.234 |
| Y | M | 8 | -0.850 |
| Y | N | 15 | -1.568 |
| Y | P | 8 | -2.206 |
| Y | Q | 13 | -0.808 |
| Y | R | 9 | -1.540 |
| Y | S | 22 | -0.844 |
| Y | T | 17 | -1.550 |
| Y | V | 14 | -0.853 |
| Y | W | 28 | 0.473 |
| Y | Y | 0 | – |

#### 2.8 Detailed numerical values for computational efficiency benchmarks

Supplementary Tables S18, S19 and S20 report the numerical values underlying panels a–c of the main-text computational-efficiency figure, respectively: prepared-input full-landscape scoring latency, effective non-wild-type substitution throughput, and input-preparation-plus-scoring latency. These comparisons include MAXWELL and eight GPU-based baselines evaluated on seven nested truncations of one protein. Effective throughput is calculated as $19L$ divided by the mean prepared-input scoring time, excluding wild-type entries.

Supplementary Table S21 summarizes the matched-length comparison at $L = 957$, and Supplementary Table S22 reports the hardware, software and timing configuration for the nine-method benchmark.

Supplementary Table S23 reports dataset-level completion times for MAXWELL, zero-shot ProteinMPNN and ThermoMPNN, corresponding to panel d. Each run generated 1,222,574 non-wild-type substitution scores across all 308 Test12K proteins. Processing times and total times including one-time model loading are reported separately.

The candidate-request analysis in panel e is described in the supplementary speed methods and results. The dataset-level and candidate-request benchmarks used ThermoMPNN's mutation-list scoring interface, whereas the matched-length benchmark reused its residue-wise amino-acid outputs. Timing comparisons therefore remain specific to their respective workloads and implementations.

**Table S18 Prepared-input full-landscape scoring latency.** Values are mean ± sample s.d. (ms) across repeated complete-landscape requests after warm-up; at least 30 repetitions per point. Input features were prepared before timing. All nine methods used the same GPU and seven nested N-terminal truncations of one protein. Column headings give sequence length $L$. Native model precision was retained.

| Method | 64 | 128 | 256 | 384 |
|---|---|---|---|---|
| MAXWELL | 6.46 ± 0.12 | 6.49 ± 0.15 | 6.52 ± 0.12 | 6.536 ± 0.037 |
| ProteinMPNN zero-shot | 6.47 ± 0.15 | 6.513 ± 0.097 | 6.584 ± 0.092 | 6.746 ± 0.081 |
| ThermoMPNN | 25.58 ± 0.33 | 44.46 ± 0.27 | 82.18 ± 0.41 | 120.00 ± 0.25 |
| HERMES | 181.20 ± 1.11 | 181.06 ± 0.90 | 181.40 ± 0.53 | 360.17 ± 1.69 |
| Mutate Everything | 14.91 ± 0.19 | 16.663 ± 0.081 | 24.821 ± 0.038 | 31.046 ± 0.040 |
| ESM-IF1 | 20.44 ± 0.34 | 25.34 ± 0.25 | 35.908 ± 0.091 | 46.21 ± 0.11 |
| VenusREM | 12.86 ± 0.17 | 13.08 ± 0.15 | 13.47 ± 0.19 | 13.21 ± 0.14 |
| ProtSSN | 162.76 ± 0.22 | 339.96 ± 1.11 | 845.64 ± 4.26 | 1334.26 ± 8.77 |
| VespaG | 16.611 ± 0.054 | 16.72 ± 0.23 | 16.579 ± 0.083 | 21.55 ± 0.17 |

| Method | 512 | 768 | 957 |
|---|---|---|---|
| MAXWELL | 6.89 ± 0.11 | 9.190 ± 0.029 | 10.84 ± 0.13 |
| ProteinMPNN zero-shot | 7.231 ± 0.062 | 9.515 ± 0.033 | 11.111 ± 0.026 |
| ThermoMPNN | 158.10 ± 0.57 | 234.76 ± 0.40 | 292.21 ± 0.58 |
| HERMES | 360.89 ± 1.00 | 542.17 ± 1.10 | 725.53 ± 0.92 |
| Mutate Everything | 38.426 ± 0.070 | 53.367 ± 0.067 | 64.075 ± 0.069 |
| ESM-IF1 | 58.027 ± 0.069 | 81.68 ± 0.47 | 102.92 ± 0.11 |
| VenusREM | 16.836 ± 0.063 | 24.34 ± 0.12 | 38.789 ± 0.038 |
| ProtSSN | 1897.26 ± 13.25 | 2974.83 ± 20.26 | 3423.41 ± 19.15 |
| VespaG | 28.949 ± 0.083 | 40.942 ± 0.095 | 44.93 ± 0.11 |

**Table S19 Effective non-WT substitution throughput.** Values are in $10^6$ substitutions/s, calculated as $19L$ divided by the mean prepared-input latency, not the mean of inverse latencies. All nine methods used the same GPU and seven nested N-terminal truncations of one protein. Column headings give sequence length $L$. Native model precision was retained.

| Method | 64 | 128 | 256 | 384 |
|---|---|---|---|---|
| MAXWELL | 0.188 | 0.375 | 0.746 | 1.116 |
| ProteinMPNN zero-shot | 0.188 | 0.373 | 0.739 | 1.082 |
| ThermoMPNN | 0.048 | 0.055 | 0.059 | 0.061 |
| HERMES | 0.007 | 0.013 | 0.027 | 0.020 |
| Mutate Everything | 0.082 | 0.146 | 0.196 | 0.235 |
| ESM-IF1 | 0.059 | 0.096 | 0.135 | 0.158 |
| VenusREM | 0.095 | 0.186 | 0.361 | 0.552 |
| ProtSSN | 0.007 | 0.007 | 0.006 | 0.005 |
| VespaG | 0.073 | 0.145 | 0.293 | 0.339 |

| Method | 512 | 768 | 957 |
|---|---|---|---|
| MAXWELL | 1.412 | 1.588 | 1.677 |
| ProteinMPNN zero-shot | 1.345 | 1.534 | 1.636 |
| ThermoMPNN | 0.062 | 0.062 | 0.062 |
| HERMES | 0.027 | 0.027 | 0.025 |
| Mutate Everything | 0.253 | 0.273 | 0.284 |
| ESM-IF1 | 0.168 | 0.179 | 0.177 |
| VenusREM | 0.578 | 0.600 | 0.469 |
| ProtSSN | 0.005 | 0.005 | 0.005 |
| VespaG | 0.336 | 0.356 | 0.405 |

**Table S20 Input preparation and complete-landscape scoring latency.** Values are mean ± sample s.d. (s) across three repetitions with resident weights. Timing starts from supplied sequence, structure and MSA as applicable; model loading, MSA search and structure prediction are excluded. All nine methods used the same GPU and seven nested N-terminal truncations of one protein. Column headings give sequence length $L$. Native model precision was retained.

| Method | 64 | 128 | 256 | 384 |
|---|---|---|---|---|
| MAXWELL | 0.00836 ± 0.00029 | 0.00918 ± 0.00025 | 0.011696 ± 0.000086 | 0.01352 ± 0.00032 |
| ProteinMPNN zero-shot | 0.01105 ± 0.00036 | 0.01405 ± 0.00047 | 0.02202 ± 0.00041 | 0.03190 ± 0.00053 |
| ThermoMPNN | 0.02957 ± 0.00024 | 0.05119 ± 0.00015 | 0.09665 ± 0.00044 | 0.14429 ± 0.00086 |
| HERMES | 0.4814 ± 0.0044 | 0.7704 ± 0.0060 | 1.608 ± 0.012 | 2.3651 ± 0.0088 |
| Mutate Everything | 0.01555 ± 0.00019 | 0.01759 ± 0.00016 | 0.02628 ± 0.00016 | 0.03295 ± 0.00017 |
| ESM-IF1 | 0.03169 ± 0.00034 | 0.04510 ± 0.00030 | 0.07349 ± 0.00049 | 0.10067 ± 0.00011 |
| VenusREM | 0.0856 ± 0.0065 | 0.1420 ± 0.0050 | 0.440 ± 0.011 | 1.248 ± 0.015 |
| ProtSSN | 0.7482 ± 0.0026 | 1.5238 ± 0.0082 | 3.6770 ± 0.0035 | 5.3988 ± 0.0081 |
| VespaG | 0.01816 ± 0.00023 | 0.01842 ± 0.00016 | 0.01861 ± 0.00014 | 0.02179 ± 0.00021 |

| Method | 512 | 768 | 957 |
|---|---|---|---|
| MAXWELL | 0.01575 ± 0.00019 | 0.02233 ± 0.00033 | 0.02711 ± 0.00022 |
| ProteinMPNN zero-shot | 0.0478 ± 0.0011 | 0.0872 ± 0.0015 | 0.1238 ± 0.0013 |
| ThermoMPNN | 0.1975 ± 0.0011 | 0.3122 ± 0.0020 | 0.40321 ± 0.00077 |
| HERMES | 3.412 ± 0.012 | 5.8306 ± 0.0046 | 7.256 ± 0.031 |
| Mutate Everything | 0.03972 ± 0.00022 | 0.05447 ± 0.00020 | 0.06431 ± 0.00011 |
| ESM-IF1 | 0.13086 ± 0.00087 | 0.18761 ± 0.00088 | 0.23477 ± 0.00045 |
| VenusREM | 2.61 ± 0.14 | 7.579 ± 0.014 | 12.905 ± 0.065 |
| ProtSSN | 7.900 ± 0.023 | 12.937 ± 0.076 | 15.766 ± 0.025 |
| VespaG | 0.02913 ± 0.00013 | 0.041942 ± 0.000094 | 0.04439 ± 0.00020 |

**Table S21 Matched complete-landscape scoring at $L = 957$.** Each request comprised 18,183 non-WT substitutions. Prepared-input and input-preparation-plus-scoring measurements are separate timing scopes. Latency ratios are baseline prepared-input latency divided by the corresponding MAXWELL latency. All measurements used the same NVIDIA RTX PRO 5000 72GB Blackwell GPU.

| Method | Prepared (ms) | Throughput ($10^6$/s) | Prep.+score (s) | Latency/MAXWELL |
|---|---|---|---|---|
| MAXWELL | 10.841 | 1.677 | 0.027 | 1.00 |
| ProteinMPNN zero-shot | 11.111 | 1.636 | 0.124 | 1.02 |
| VenusREM | 38.789 | 0.469 | 12.905 | 3.58 |
| VespaG | 44.933 | 0.405 | 0.044 | 4.14 |
| Mutate Everything | 64.075 | 0.284 | 0.064 | 5.91 |
| ESM-IF1 | 102.924 | 0.177 | 0.235 | 9.49 |
| ThermoMPNN | 292.209 | 0.062 | 0.403 | 26.95 |
| HERMES | 725.531 | 0.025 | 7.256 | 66.93 |
| ProtSSN | 3423.408 | 0.005 | 15.766 | 315.79 |

**Table S22 Computational environment for the GPU efficiency benchmark.** These settings describe the nine-method inference timing experiment, not all experiments in the study. The seven inputs are nested truncations of a single protein. Timing uses GPU-synchronized wall-clock measurements.

| Item | Setting |
|---|---|
| GPU | NVIDIA RTX PRO 5000 72GB Blackwell |
| Reported GPU memory | 73,415 MiB |
| NVIDIA driver | 595.84 |
| PyTorch / CUDA runtime | 2.12.1+cu132 / 13.2 |
| Protein batch size / allocated CPUs | 1 / 8 |
| TF32 | Disabled |
| Native precision | FP32; VespaG: FP16 encoder and FP32 head |
| Prepared-input repetitions | 10 warm-ups; at least 30 measurements and 5 s |
| Preparation-plus-scoring repetitions | 3; weights resident |
| Sequence lengths | 64, 128, 256, 384, 512, 768, 957 |
| Input protein | `fireprotdb_1UZC_ddG` |
| Non-WT request size | $19L$ |
| Ensembles | HERMES: 10 members; ProtSSN: 9 members |

**Table S23 Dataset-level completion of Test12K mutation landscapes.** Each run generated 1,222,574 non-wild-type substitution scores across all 308 proteins. Values are medians across three complete runs. Processing includes per-protein input preparation, scoring, score extraction and output writing. Total time additionally includes one-time model loading. ThermoMPNN used its mutation-list scoring interface; these timings should not be combined with the residue-wise-output-reuse measurements in panels a–c.

| Method | Processing time (s) | Total time incl. loading (s) |
|---|---|---|
| MAXWELL | 9.03 | 16.98 |
| ProteinMPNN zero-shot | 11.45 | 11.67 |
| ThermoMPNN | 354.89 | 363.34 |

**Table S24 Sensitivity of melting-temperature screening performance to the stabilizing-mutation threshold.** AUROC, AUPRC and Enrich@10 are reported for MAXWELL and the baseline methods at $\Delta T_m \geq 0.5$, $\geq 1$ and $\geq 2\,^\circ$C on the FireProt2 dataset, with $1\,^\circ$C as the primary threshold. Values are mean ± s.d. across eligible proteins. Bold means mark the highest value for each metric within each threshold. Group headings give the numbers of proteins contributing to AUROC/AUPRC and Enrich@10.

| Method | AUROC↑ | AUPRC↑ | Enrich@10↑ |
|---|---|---|---|
| $\Delta T_m \geq 0.5\,^\circ$C; AUROC/AUPRC: $n = 70$; Enrich@10: $n = 50$ | | | |
| MAXWELL | **0.750**±0.209 | 0.635±0.278 | 1.875±1.040 |
| ThermoMPNN | 0.748±0.209 | 0.636±0.285 | **1.895**±1.113 |
| HERMES | 0.722±0.230 | 0.637±0.279 | 1.840±1.088 |
| Rosetta | 0.741±0.209 | **0.642**±0.288 | 1.613±0.964 |
| Mutate Everything | 0.655±0.251 | 0.577±0.285 | 1.442±0.986 |
| FoldX | 0.654±0.222 | 0.557±0.283 | 1.498±0.826 |
| ProteinMPNN zero-shot | 0.693±0.229 | 0.600±0.273 | 1.853±1.079 |
| ProtSSN | 0.662±0.231 | 0.593±0.269 | 1.656±1.006 |
| VenusREM | 0.670±0.248 | 0.600±0.295 | 1.551±1.047 |
| RSALOR | 0.631±0.223 | 0.532±0.273 | 1.504±0.978 |
| VespaG | 0.627±0.233 | 0.546±0.279 | 1.458±0.953 |
| ESM-IF1 | 0.648±0.219 | 0.573±0.264 | 1.696±0.931 |
| $\Delta T_m \geq 1\,^\circ$C; AUROC/AUPRC: $n = 66$; Enrich@10: $n = 49$ | | | |
| MAXWELL | **0.737**±0.218 | 0.578±0.287 | **2.103**±1.598 |
| ThermoMPNN | 0.733±0.206 | 0.579±0.288 | 1.988±1.153 |
| HERMES | 0.716±0.234 | **0.597**±0.295 | 2.044±1.428 |
| Rosetta | 0.703±0.227 | 0.577±0.306 | 1.630±1.105 |
| Mutate Everything | 0.648±0.260 | 0.535±0.300 | 1.528±1.115 |
| FoldX | 0.630±0.234 | 0.509±0.299 | 1.606±1.198 |
| ProteinMPNN zero-shot | 0.687±0.238 | 0.572±0.285 | 1.971±1.322 |
| ProtSSN | 0.643±0.231 | 0.547±0.299 | 1.612±1.106 |
| VenusREM | 0.651±0.242 | 0.541±0.312 | 1.562±1.147 |
| RSALOR | 0.602±0.214 | 0.480±0.282 | 1.397±1.084 |
| VespaG | 0.595±0.249 | 0.477±0.292 | 1.412±1.080 |
| ESM-IF1 | 0.625±0.234 | 0.518±0.275 | 1.830±1.674 |
| $\Delta T_m \geq 2\,^\circ$C; AUROC/AUPRC: $n = 58$; Enrich@10: $n = 43$ | | | |
| MAXWELL | **0.748**±0.219 | 0.568±0.295 | **2.709**±3.406 |
| ThermoMPNN | 0.733±0.211 | 0.544±0.320 | 1.975±1.054 |
| HERMES | 0.727±0.231 | **0.574**±0.317 | 2.094±1.723 |
| Rosetta | 0.679±0.239 | 0.520±0.336 | 1.405±0.848 |
| Mutate Everything | 0.639±0.243 | 0.490±0.327 | 1.374±1.149 |
| FoldX | 0.651±0.235 | 0.499±0.302 | 1.755±1.479 |
| ProteinMPNN zero-shot | 0.672±0.224 | 0.521±0.279 | 1.794±1.118 |
| ProtSSN | 0.602±0.244 | 0.501±0.310 | 1.490±1.361 |
| VenusREM | 0.628±0.250 | 0.496±0.323 | 1.537±1.276 |
| RSALOR | 0.546±0.223 | 0.412±0.279 | 1.229±1.116 |
| VespaG | 0.550±0.237 | 0.412±0.260 | 1.714±2.736 |
| ESM-IF1 | 0.614±0.213 | 0.464±0.284 | 1.884±2.566 |

### References

[1] Dieckhaus, H., Brocidiacono, M., Randolph, N.Z., Kuhlman, B.: Transfer learning to leverage larger datasets for improved prediction of protein stability changes. Proceedings of the National Academy of Sciences **121**(6), 2314853121 (2024)

[2] Visani, G.M., Pun, M.N., Galvin, W., Daniel, E., Borisiak, K., Wagura, U., Nourmohammad, A.: Hermes: Holographic equivariant neural network model for mutational effect and stability prediction. arXiv preprint arXiv:2407.06703 (2024)

[3] Dauparas, J., Anishchenko, I., Bennett, N., Bai, H., Ragotte, R.J., Milles, L.F., Wicky, B.I., Courbet, A., Haas, R.J., Bethel, N., *et al.*: Robust deep learning–based protein sequence design using proteinmpnn. Science **378**(6615), 49–56 (2022)

[4] Schymkowitz, J., Borg, J., Stricher, F., Nys, R., Rousseau, F., Serrano, L.: The foldx web server: an online force field. Nucleic acids research **33**(suppl_2), 382–388 (2005)

[5] Alford, R.F., Leaver-Fay, A., Jeliazkov, J.R., O'Meara, M.J., DiMaio, F.P., Park, H., Shapovalov, M.V., Renfrew, P.D., Mulligan, V.K., Kappel, K., *et al.*: The rosetta all-atom energy function for macromolecular modeling and design. Journal of chemical theory and computation **13**(6), 3031–3048 (2017)

[6] Ouyang-Zhang, J., Diaz, D.J., Klivans, A.R., Krähenbühl, P.: Predicting a protein's stability under a million mutations. In: Advances in Neural Information Processing Systems, vol. 36 (2023)

[7] Hsu, C., Verkuil, R., Liu, J., Lin, Z., Hie, B., Sercu, T., Lerer, A., Rives, A.: Learning inverse folding from millions of predicted structures. In: International Conference on Machine Learning, pp. 8946–8970 (2022). PMLR

[8] Tan, Y., Wang, R., Wu, B., Hong, L., Zhou, B.: From high-throughput evaluation to wet-lab studies: advancing mutation effect prediction with a retrieval-enhanced model. Bioinformatics **41**(Supplement 1), 401–409 (2025) https://doi.org/10.1093/bioinformatics/btaf189

[9] Tsishyn, M., Hermans, P., Rooman, M., Pucci, F.: Residue conservation and solvent accessibility are (almost) all you need for predicting mutational effects in proteins. Bioinformatics **41**(6), 322 (2025) https://doi.org/10.1093/bioinformatics/btaf322

[10] Tan, Y., Zhou, B., Zheng, L., Fan, G., Hong, L.: Semantical and geometrical protein encoding toward enhanced bioactivity and thermostability. eLife **13**, 98033 (2025) https://doi.org/10.7554/eLife.98033

[11] Marquet, C., Schlensok, J., Abakarova, M., Rost, B., Laine, E.: Expert-guided protein language models enable accurate and blazingly fast fitness prediction. Bioinformatics **40**(11), 621 (2024) https://doi.org/10.1093/bioinformatics/btae621

[12] Steinegger, M., Söding, J.: Mmseqs2 enables sensitive protein sequence searching for the analysis of massive data sets. Nature biotechnology **35**(11), 1026–1028 (2017)

[13] Rost, B.: Twilight zone of protein sequence alignments. Protein engineering **12**(2), 85–94 (1999)

[14] Likic, V.: The needleman-wunsch algorithm for sequence alignment. Lecture given at the 7th Melbourne Bioinformatics Course, Bi021 Molecular Science and Biotechnology Institute, University of Melbourne, 1–46 (2008)
